# Shape-centered representations of bounded regions of space mediate the transformation of retinotopic representations into conscious perception of objects

**DOI:** 10.1101/2020.05.29.123901

**Authors:** G. Vannuscorps, A. Galaburda, A. Caramazza

## Abstract

The primary visual cortex represents the retinotopic orientation of visual primitives (edges, blobs, bars), but our conscious perception is of orientated objects (e.g., dogs, forks) in the environment. How this transformation operates remains unknown. We report here the study of a young woman presenting with an extraordinarily clear and informative visual disorder that affects highly specific aspects of object perception allowing precise inferences about the type and properties of visual representations that mediate this transformation. Davida perceives sharp-edged 2D bounded regions of space of medium to high contrast as if they were plane-rotated by 90, 180 or 270 degrees around their center, mirrored across their own axes, or both. In contrast, her perception of strongly blurred or very low contrast shapes, and of compound shapes emerging from a collection of bounded elements, is intact. The nature of her errors implies that visual perception is mediated by a representation of each bounded region of space in a shape-centered coordinate system aligned on either the shape’s most elongated part or on the shape’s axis of symmetry and centered either at the midpoint of the shape’s most elongated part or at the shape’s centroid. The selectivity of her disorder to sharp-edged medium to high-contrast stimuli additionally suggests that duplicate shape-centered representations are computed in parallel from information derived from the parvocellular and magnocellular subcortical channels and integrated precisely at the level at which shape representations must be mapped onto a behaviorally relevant frame of reference.

## Introduction

The primary visual cortex represents visual primitives (local spatial frequency patches, edges, blobs, bars, terminators) in a retinotopically organized map of the visual field (De Valois, Albrecht, & Thorell, 1982; Felleman & Van Essen, 1991; Hubel & Wiesel, 1962; Wandell, Dumoulin, & Brewer, 2007) but we eventually perceive objects (e.g., dogs, faces, forks) and their relative location and orientation with respect to our bodies and other objects in the environment (Colby, 1998; Connor & Knierim, 2017; McKyton & Zohary, 2007; Melcher & Morrone, 2015; Milner & Goodale, 2006; Rock, 1973). Our conscious perception of the world remains stable across eye movements and although a stationary vertical line moves and rotates in retinotopic coordinates when we move and rotate our head, *phenomenally*, it remains a vertical and stationary line. A fundamental question concerns the mechanisms involved in this transformation of visual information from primitives to objects and, for their associated coordinate systems, from retinocentric to ego and allocentric. Although much progress has been made in addressing this question, much remains to be learned about the nature of the representations that characterize this process (Cadieu et al., 2007; DiCarlo, Zoccolan, & Rust, 2012; Marr & Nishihara, 1978; Palmer & Rock, 1994; Pasupathy & Connor, 2001; Peirce, 2015; Serre, Oliva, & Poggio, 2007; Yamins et al., 2014).

Progress in understanding the levels of representations involved in the transformation of retinotopic representations into conscious perception of objects is hindered by the extreme degree of complexity and interactions between multiple levels of representations in the visual system, making it extremely difficult to isolate and study the nature of one particular level. Nevertheless, nature occasionally provides the opportunity to peer inside extremely complex neural systems by isolating components of a system through accidental damage or genetic modification of neural components.

We report here the detailed study of a young woman (Davida), who has no remarkable medical, neuropsychological, neurological, psychiatric or ophthalmological history (see Appendix Case History), but presents with an extraordinarily clear and informative visual disorder that affects a highly specific aspect of object perception. Davida reports perceiving any sharp-edged 2D medium to high-contrast bounded region of space (e.g., black letters, arrows, abstract shapes on white background) alternating through piece-meal gradual transition between their correct orientation and all the other orientations that would result from their mirroring across one or both of their own axes, their rotation by 90, 180 or 270 degrees around their center, or both (see Movie S1 online for a description of what she perceives when shown these types of stimuli). The results of the experiments probing Davida’s perception of orientation through verbal judgments, visual illusions, direct copy, and directed movements fully corroborated this difficulty (see, for examples, Movies S2 – S7). In contrast, (a) the processing of orientation from auditory, tactile and kinesthetic information is intact (see, for example, Movies S8); (b) visual judgments about the identity, shape, distance, color, size, movement and location of the same kind of stimuli are intact; and (c) the perception of the orientation of the same shapes (letters, arrows, abstract shapes) shown in 3D, or under very low luminance contrast or very low spatial frequencies, and of compound shapes composed of a collection of bounded elements is intact. This highly selective deficit in the perception of the orientation (and not of other characteristics) of sharp-edged 2D medium to high-contrast (and not 3D, blurred or low contrast) bounded region of space (and not compound shapes) forces several new conclusions about the nature of the mechanisms involved in transforming retinotopic into spatiotopic representations of visual information.

## Experimental study

### Participants

A detailed case report of Davida’s medical, neuropsychological, neurological, psychiatric and ophthalmological history is provided in Appendix (Case History). Some of the experimental tasks were also presented to control participants. The control group was composed of 14 females (11 were right-handed), slightly older (mean age = 19.6; range = 18-21) and more educated (mean years of college education = 2.15; range = 1-4) than Davida. The control participants had normal or corrected visual acuity and reported no antecedent developmental disorders.

### Material and procedure

The experimental investigations were carried out from October 2016 to March 2019 during sessions lasting between 60 and 120 minutes. The study was approved by the Committee on the Use of Human Subjects, Harvard University (Protocol # IRB16-1124). Written informed consent (control participants), assent (Davida) or permission (Davida’s parents) were obtained prior to the study. Unless otherwise indicated, in all experiments participants were seated in front of a laptop computer at 50 cm from the screen. The room was dimly illuminated from the ceiling. All experiments were controlled with the Psychopy software (Peirce, 2007, 2009), and all visual stimuli were displayed on a Lenovo T460s 14 inch, 16:9, 1920 x 1080 pixels (157 PPI), 60Hz screen controlled by an Intel® HD Graphic 520 graphics card.

A detailed description of the material and procedures of all the experiments is provided in the Appendix. Supplementary Movies can be accessed on the Open Science Framework platform (link: https://osf.io/pf56m/?view_only=bda3dcc0b9ea4d62ac122e23d8227463).

### Results

The main conclusions afforded by Davida’s behavioral profile, schematized in Figure 1, follow from 6 sets of results, §1-§6.

**Figure 1.**
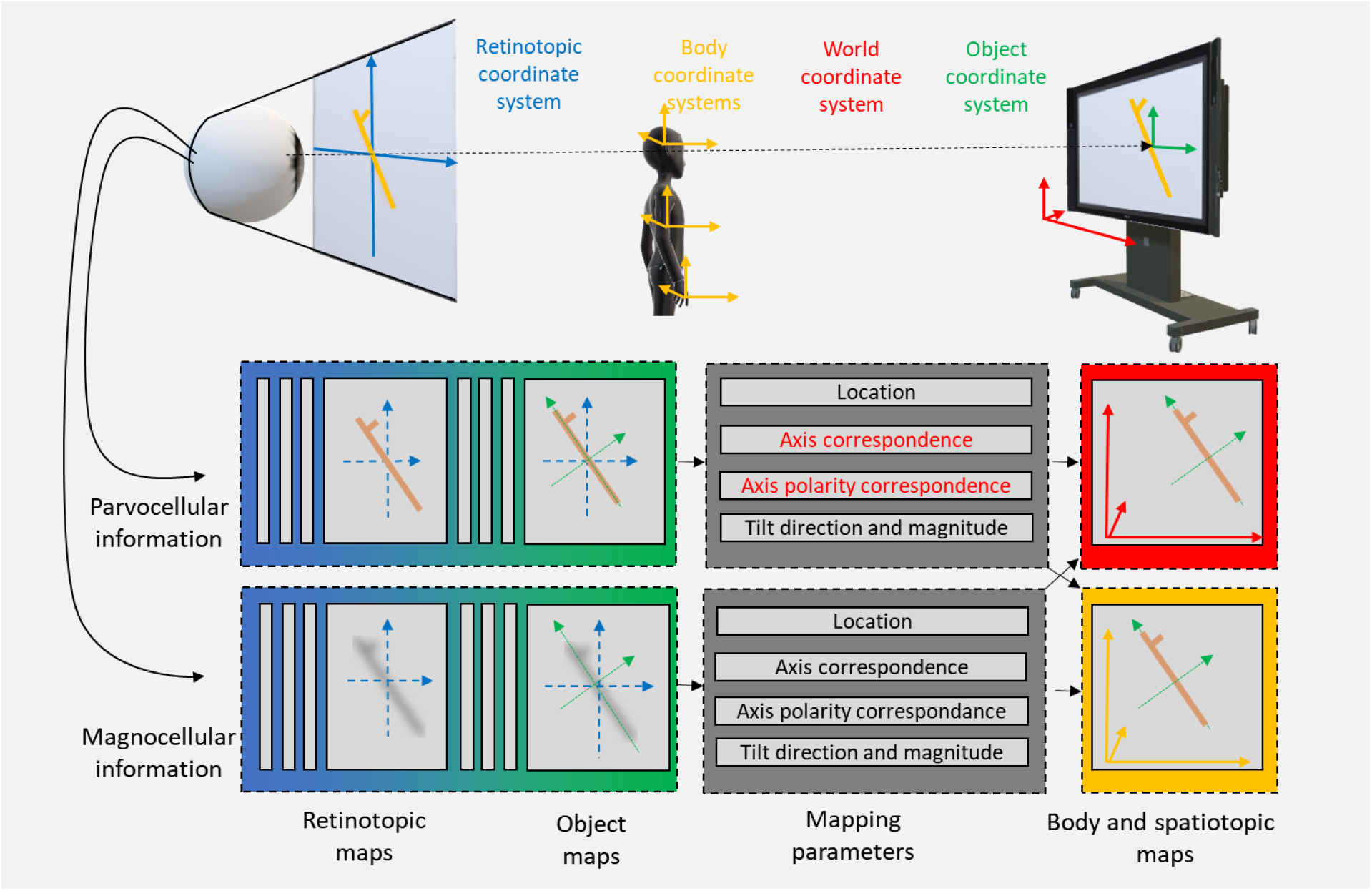
Schematic representation of the main conclusions drawn from Davida’s behavioral profile. Observed objects are projected onto the retina in retinotopic space (in blue). From the retina, information is conveyed to the brain through a parvocellular pathway composed of cells mostly sensitive to sharp-edged, fine, stationary, and high-contrast stimuli and a magnocellular pathway mostly activated by stimuli with complementary characteristics (coarse, large, moving, brief, and low-contrast). The primary visual cortex represents this information in retinotopic coordinates (in blue). Behavior requires a transformation from retinotopic coordinates to non-retinotopic coordinates (e.g., spatiotopic and body-centered, in red and yellow). The results reported here show that this transformation is mediated by an intermediate, unconscious, stage of processing where the visual system represents bounded regions of space in their own “object-centered” coordinate system composed of orthogonal axes aligned either on the shape’s most elongated part (i.e., for elongated objects, as displayed in the Figure) or on the shape’s axis of symmetry (for non-elongated symmetrical objects, not displayed in the Figure), and centered either at the center of elongated shapes’ most elongated part or at the centroid of symmetrical shapes. Davida’s behavioral profile also suggests that these “Shape-Object-centered representations” are computed in parallel from information derived from the parvocellular and magnocellular channels and are integrated precisely at the level at which Shape-Object-centered representations must be mapped onto a behaviorally relevant frame of reference. Her disorder affects selectively two of the parameters – the axis correspondence and axis polarity correspondence parameters (in red; McCloskey, Valtonen, & Cohen Sherman, 2006) required to map Shape-Object-centered representations computed (correctly) from information carried in the parvocellular pathway onto behaviorally relevant coordinate frames.

[§1] Upon initial questioning, Davida reported seeing letters and other 2-dimensional (2D) stimuli (e.g., numbers and road signs), but not daily life’s 3-dimensional (3D) stimuli, in different orientations rapidly alternating through piece-meal gradual transitions “as if the letter was fading in, fading out in different orientations” (see Movie S1). This description, which is similar to that typically reported during rivalry (Blake, 2001), suggested the visual system’s attempt to resolve a perceptual problem. We tested Davida in a series of experiments probing her perception of the orientation of 2D shapes either explicitly through verbal judgments and direct copy or implicitly through naming, visual after-effects, visual illusions, stimulus-response compatibility effects and immediate and delayed directed movements. We had three objectives. The first was to characterize the set of orientations that she perceives when shown different types of stimuli. The second was to explore whether Davida’s disorder similarly affects explicit (Appendix 1.1 – 1.5) and implicit (Appendix 1.6 – 1.13) perceptual judgment tasks, which are widely assumed to be resolved based on a spatiotopic representation of visual information (Appendix 1.1 – 1.7), and action tasks, which call into play body-centered representations of visual information (Appendix 1.9 and 1.10). Davida’s performance and response profile in this series of experiments revealed a clear and coherent pattern: Davida consciously perceives 2D stimuli to be inverted (e.g., b → p), reversed (b → d), or plane-rotated by 90 or 180 degrees (e.g., Figure 2A-E; Appendix 1.1-1.13; Movie S2-7), and her disorder is quantitatively and qualitatively independent of the nature of the task (e.g., implicit, explicit) and of the nature of the high-order coordinate frame called into play to solve the task (body-centered or spatiotopic). When presented with a black arrow and asked to carefully place her finger on the tip of that arrow, for instance, she almost systematically pointed to where the tip of the arrow would have been if the arrow were rotated by 90 or 180 degrees around its center (Figure 2E; Appendix 1.7, 1.10; Movie S5, S7).

**Figure 2.**
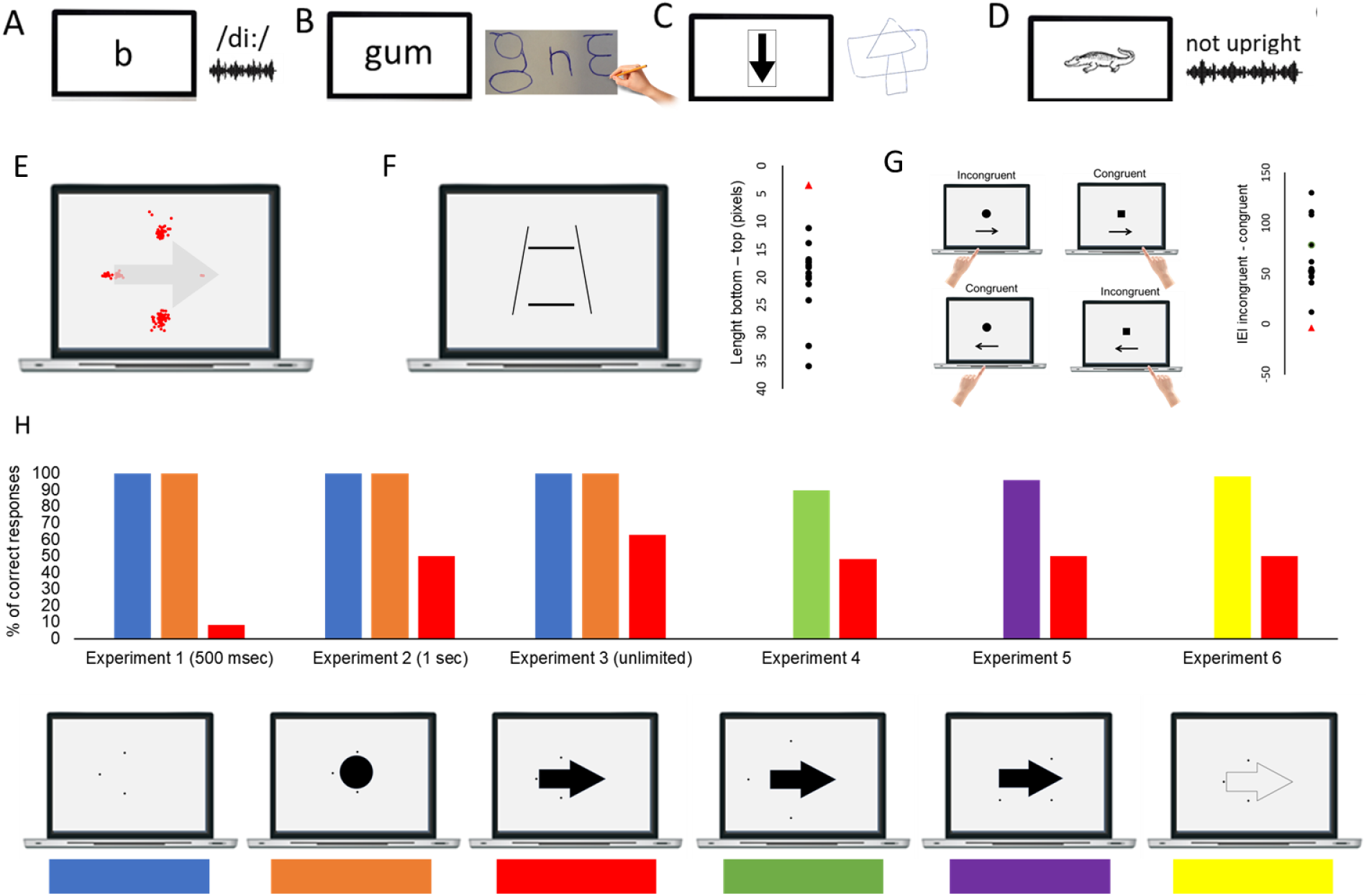
Davida named (A), copied (B, C) and judged (D) the orientation of letters and line drawing as if they were inverted or rotated by 90 or 180 degrees. (E) Asked to place her finger or the mouse cursor on the tip of a displayed arrow, Davida typically pointed to where it would have been if the arrow were inverted or rotated by 90 or 180 degrees (the red dots, Appendix 1.7; see also Movie S5). (F) When asked to match the size of the two horizontal lines in the Ponzo illusion display, control participants (black dots) typically underestimated the length of the lower horizontal line (*t* (13) = 11.5, *p* < 0.001; the Ponzo illusion), but not Davida (red triangle; *t* (19) = 0.72), who instead was significantly more accurate than controls (Crawford & Howell’s 1998 modified *t* test: *t* (13) = −2.45, *p* = 0.01; Appendix 1.12). (G) In responding as fast as possible to a circle or a square with the left or right index finger, respectively, the controls’ (black dots) inverse efficiency index (IEI; mean response latency divided by accuracy rate) showed the typical advantage for congruent trials. Davida (red triangles), however, differed significantly from the controls and showed no congruency effect (Appendix 1.11). (H) Presented with one, two or three small dots for various amounts of time across 6 experiments, Davida systematically failed to indicate the correct number of dots (1, 2 or 3) presented on the screen when they were presented together with a large black arrow that would overlap with their location if it were rotated by 90 or 180 degrees (red condition), but she was excellent in the other conditions (Appendix 3.6).

A third goal was to test two predictions derived from Davida’s report of the orientation of the 2D stimuli: (1) if she perceives shapes in inaccurate orientations, then, Davida should perform far better than control participants in tasks, such as visual illusions and stimulus-response compatibility tasks, in which accurate orientation perception typically hinders performance; (2) if she sees 2D stimuli randomly fluctuating between different orientations alternating through piecemeal gradual transitions, then, Davida should be slow at identifying 2D objects. These two predictions were confirmed (see Appendix 1.11 – 1.14). Davida, for instance, was extraordinarily efficient in the Ponzo illusion task (Figure 2F) and, unlike control participants, she was not influenced by the orientation of an arrow during a typical stimulus-response compatibility task (Figure 2G).

[§2] Davida reports no other visual difficulty. This was confirmed in two series of experiments. The first series comprised experiments probing her perception of the shape, size, location, distance, movement and tilt of 2D stimuli (2.1 – 2.6). The goal of these experiments was to explore whether her disorder affected other aspects of visual processing. Davida’s performance in these experiments was as good as control participants, including her ability to discriminate the tilt of shapes. Davida had no difficulty discriminating abstract shapes when one had an edge 0.05 degree of visual angle longer than the comparison ones (Appendix 2.1) or to discriminate shapes tilted less than one degree of visual angles from each other (Appendix 2.5), for instance. The second series of experiments examined her ability to process the orientation/location of kinesthetic, tactile, and auditory stimuli, and her ability to form and use internal representations of oriented shapes (2.7 – 2.9). She performed these experiments flawlessly. For instance, she had no difficulty to name orientation sensitive letters (b, p, d, q) traced on her hand (Appendix 2.8, Movie S8) or to write these letters to dictation, and hence from memory, on a sheet of paper (Appendix 2.9). All this implies that Davida’s disorder is specific to vision and consists only in perceiving 2D shapes as if they were inverted, reversed, or plane-rotated by 90 or 180 degrees.

The sets of results §1-2 severely constrain hypotheses about the functional locus of Davida’s perceptual deficit. That Davida literally sees 2D shapes in incorrect orientations and has a consistent proportion and type of errors in all (but only visual) tasks implies that her deficit is at a stage in the visual processing stream that is common (and thus preliminary) to the different types of “higher” representational frames (e.g., spatiotopic, body-centered) involved in perception and action tasks. This pattern of performance contrasts with the fact that she was as good as control participants in judging the shape, size, location, distance, tilt, and movement of 2D stimuli, thus implying that her disorder arises at a level in the visual system at which, or beyond which, the shape of these stimuli has been computed accurately. Thus, her disorder affects representations in the visual system involved in transforming intact representations of shapes into higher-level frames of reference underlying action and conscious perception (Figure 1). In all this, Davida differs sharply and instructively from previous reports of neurological individuals who suffered from difficulties in reporting, naming, judging, memorizing, reproducing and/or comparing the orientation of objects. A majority of these cases had difficulties in only some visual tasks (Cooper & Humphreys, 2000; Davidoff & Warrington, 1999; Davidoff & Warrington, 2001; Harris, Harris, & Caine, 2001; Karnath, Ferber, & Bülthoff, 2000; Martinaud et al., 2016, 2014; Priftis, Rusconi, Umiltà, & Zorzi, 2003; Riddock et al., 2004; Robinson, Cohen, & Goebel, 2011; Turnbull, Beschin, & Della Sala, 1996; Turnbull, Laws, & McCarthy, 1995; Turnbull & McCarthy, 1996). Other patients displayed either orientation errors in several modalities (e.g., visual, motor, tactile) or a visual deficit that was not selective to orientation (McCloskey, 2009; McCloskey et al., 2006; Pflugshaupt et al., 2007; Valtonen, Dilks, & McCloskey, 2008). Davida’s disorder offers a unique opportunity to investigate the nature of the representations and mechanisms involved in the course of transforming retinotopic coordinates into environmental ones.

[§3] That Davida sees 2D objects reversed, inverted or plane rotated with respect to their own center (see Figure 2 A-E) suggests that Davida’s disorder emerges at a level at which each object is represented in a spatial coordinate system located at the center of the objects, independently of their background, of other objects, and of their retinotopic representation – a shape- or object-centered coordinate system. Three predictions of this conclusion were tested and confirmed: (1) her subjective report, error rates, and error distributions in experiments assessing her perception of sharp-edged 2D stimuli were independent of the eye(s) used, the location of the stimulus in the visual field, and where she focuses her visual attention (Appendix method and results 3.1, 3.2); (2) when presented simultaneously with two bounded objects, Davida reported perceiving them as the result of different, independent transformations (see Figure 2C and Appendix method and results 3.3 – 3.5); (3) Davida has difficulty detecting stimuli located in an area that would be covered by another object (e.g., black solid arrow) if that object were rotated by 90 degrees or inverted (Appendix methods and results 3.6, 3.7 and Figure 2H). Hence, as shown on Figure 2H, Davida was able to correctly report whether one, two or three black dots were presented on the screen when the dots were displayed alone (blue condition), when they were displayed together with a large black circle (orange condition) or with a “transparent” arrow defined only by its contour (yellow condition), and when they were placed outside the area that would be covered by a large black arrow if that arrow were rotated by 90 or 180 degrees (green and purple condition), but not when they were placed in an area that would be covered by the same large black arrow if it were rotated by 90 or 180 degrees (red condition). All her errors in the latter condition consisted in underestimating the number of dots that had been displayed in that condition.

[§4] That Davida’s disorder affects a level of processing where objects are represented with respect to their own, intrinsic, frame (an “object-centered” representation) offers the opportunity to explore what is an “object” at that stage of processing. Davida’s response profile in a series of experiments conducted to address this issue indicated that her disorder affects the perception of the orientation of areas in the visual field bound by sharp (luminance or chromatic) borders (Appendix method and results 4.1 – 4.14, Movie S9-12; see Figure 3 for examples). When asked to copy words, for instance, Davida misrepresented the orientation of individual letters when the letters were unconnected but also of the whole word when the letters were connected (Appendix 4.1). When shown a series of arrows made of two colors separated by a sharp edge and asked to use the computer mouse to move a small round cursor and click as precisely as possible on the tip of the arrow, Davida almost systematically (in 78.12% of the trials) clicked approximately (i.e. less than 50 pixels away) where the tip of that arrow would have been if only the colored part of the arrow of the same color as the tip had been rotated by 90, 180 or 270 degrees (Appendix 4.5; Figure 3B and S17A). In contrast, as shown on Movie S10, when shown a series of arrows made of two colors transitioning very progressively from one to another, the bicolor arrow was almost always perceived as a single rotated object (Appendix 4.5; Figure 3B and S17C). Thus, her disorder affects a stage of processing in which bounded areas of the visual field separated by clear edge are represented independently of each other. Additional evidence in support of this conclusion is provided by the finding that Davida has no difficulty to perceive the orientation of compound shapes emerging from an arrangement of bounded elements, such as arrows composed of non-connected dots or of multiple parts of different colors (Appendix 4.8 – 4.14; Movie S11, 12; Figure 3C-F). For instance, Davida has no difficulty copying, judging or naming orientation-sensitive letters (b, d, p, q) or the orientation of objects when the letters and objects are composed of non-connected dots or of multiple parts of different colors and, while this was not the case with a solid black arrow (Figure 2G), her response latencies in a stimulus-response compatibility task were significantly influenced by the presence of a to-be ignored dotted arrow (Appendix 4.11; Figure 3F).

**Figure 3.**
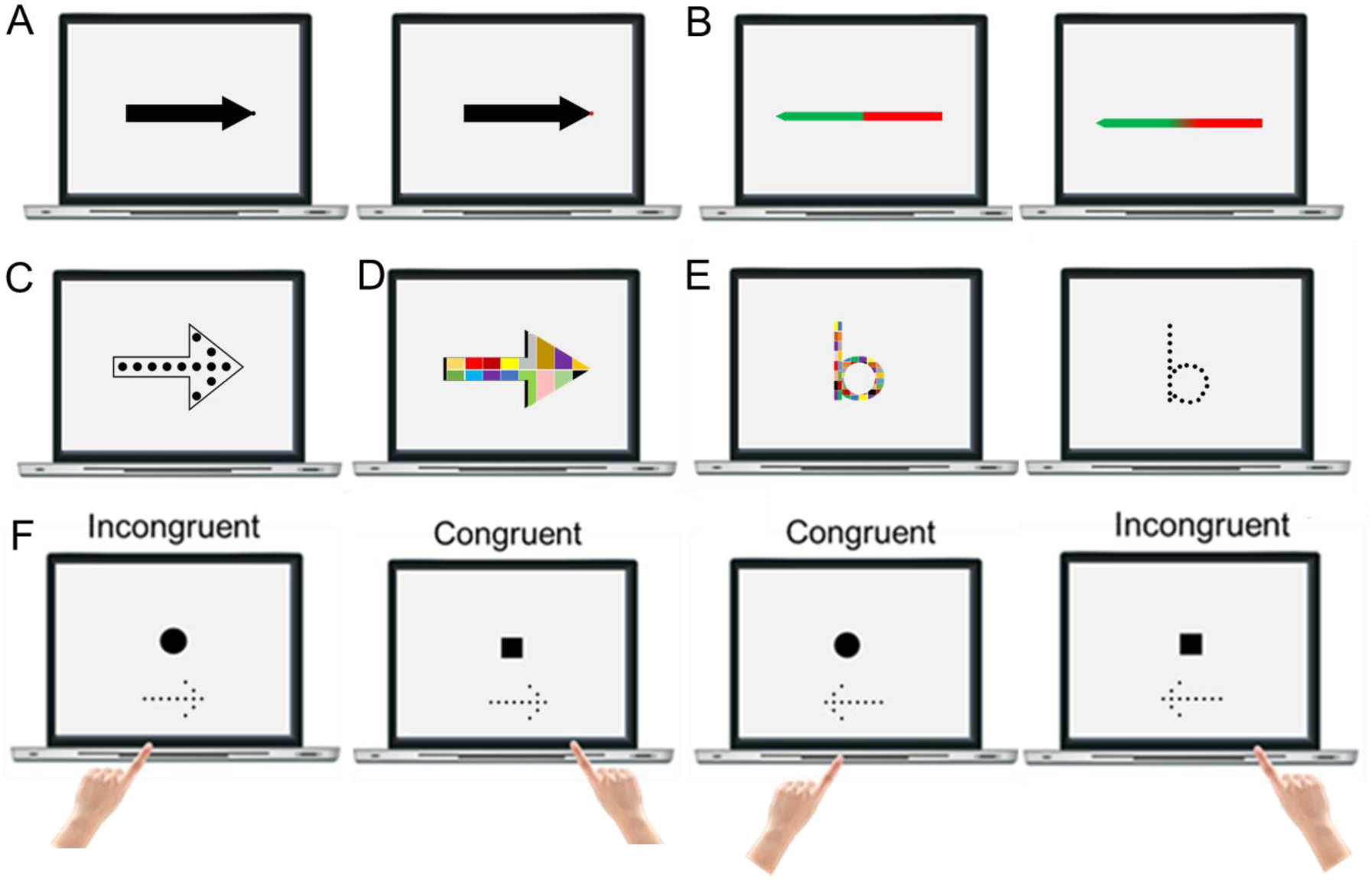
(A) Asked to click as precisely as possible with a mouse-controlled cursor on the dot at the tip of the arrow, Davida clicked on the dot in only 1/20 trials when the dot was black but in 100% of the trials when the dot was of a different color (Appendix 4.2). (B) Asked to point to the tip of these arrows, Davida’s errors consisted mostly of clicking where the tip would have been if only the colored part of the arrow of the same color as the tip had been rotated by 90, 180 or 270 degrees when the colors were separated by a sharp edge (19/26 errors), but where the tip would have been if the whole arrow had been rotated by 90, 180 or 270 degrees when the colors were blended over a large area (23/27 errors) (see also Appendix 4.5 and Movie S10). (C) When shown an arrow implied by a series of unconnected small dots within an arrow composed of solid black lines, Davida mislocated the tip of the arrow in 87.5% of the trials when asked to click on the tip of the solid arrow, but was flawless when asked to click on the tip of the dotted arrow (see also Appendix 4.10 and Movie S11 – 12). (D). Davida was flawless when asked to click on the tip of a large arrow composed of segments of different colors (Appendix 4.10). (E) Davida named flawlessly orientation sensitive letters composed of connected parts of different colors or composed of small black dots (Appendix 4.14). (F) In responding as fast as possible to a circle or a square with the left or right index finger, respectively, Davida showed the typical advantage for congruent trials when shown dotted arrows (one-tailed *t* (53) = 2.05, p = 0.02), but not when shown solid arrows (Appendix 1.10 and 4.11).

The findings in §4 introduce a distinction between two levels of object representation: objects defined strictly by bounded regions of space, which we will refer to as Shape-Object (S-Object), and compound objects, composed of independent parts but perceived as a single object (e.g., a shape composed of unconnected dots; Figure 3C-F; Figure S20-26), which we will refer to as Perception-Object (P-Object).

This distinction parallels that proposed on theoretical grounds by Palmer and Rock (Palmer & Rock, 1994; see also Tse & Palmer, 2012) between entry-level “uniform connected regions” (UCRs) and postconstancy levels of representations. Like the S-object-centered representations affected by Davida’s disorder, the UCRs are defined as connected regions of uniform image properties (e.g., luminance, color) and were hypothesized to serve as the fundamental first unit of perceptual organization, emerging from the processes of edge detection in early vision and laying the foundation on which all later perceptual organization rests. Consciously perceived organizations of UCRs derived from parsing and grouping operations (Wagemans et al., 2012) were hypothesized to emerge at later “postconstancy” levels of representation.

[§5] That Davida’s disorder occurs at the level of mapping correctly computed “S-Object-centered” representations onto behaviorally relevant non retinotopic frames affords the additional opportunity to explore the geometric properties that determine how a coordinate frame is assigned to S-Objects. In two series of experiments, we aimed to characterize the geometrical parameters used by the visual system to ascribe coordinate axes to elongated asymmetrical stimuli (5.1 – 5.8; Figure 4) and to symmetrical stimuli deprived of a straight segment (5.9 – 5.11; Figure 5). We used tilted stimuli because, unlike the upright objects used in the previous experiments (e.g., Figure 2), they allow discriminating reflections across retinotopic, body-centered, allocentric (spatiotopic, gravitational) and object-based reference frames (McCloskey et al., 2006). When presented with tilted, asymmetrical, elongated stimuli in 8 experiments (Appendix 5.1 – 5.8; Figure 4 for examples), Davida systematically made 7 types of errors (e.g., Figure 4; see also Figure 6 E-K; Movies S13-16). All these errors resulted from transformations of the stimulus (rotations, mirror reflection or both) within a frame constituted by an axis aligned precisely on the shape’s longest straight segment (Appendix 5.6) and a perpendicular axis intersecting the elongation axis precisely at its geometrical center (Appendix 5.7 – 5.8). For example, when shown a tilted, asymmetrical, elongated, target stimulus and three probe stimuli that were mirror reflections of the target across either an axis aligned on the shape’s longest straight segment (Chaisilprungraung, German, & McCloskey, 2019), an axis relating the two most distant points of the shape (longest span axis; Sekuler & Swimmer, 2000), or the axis that minimizes the sum of squared distances to all points of the shape (the axis of least second moment; Haralick & Shapiro, 1991) and asked to indicate whether one of these probes corresponded to a perceived orientation of the target, Davida pointed to the probe corresponding to a mirror reflection of the target across the shape’s longest straight segment in 100% of the trials (Appendix 5.6; Figure 4C). To explore whether the center of the representational frame of an elongated shape is the center of the shape’s longest straight segment or the shapes’ centroid (mean coordinate of all the points in the shape), we showed Davida an elongated asymmetrical shape and asked her to click on the screen where she saw the different perceived orientations of that shape intersecting (Appendix 5.8, see also Appendix 5.7). The coordinates at which she indicated seeing two lines crossing were on average 14.9 pixels from the center but 45 pixels from the centroid of the asymmetrical elongated shape. When presented with symmetrical shapes devoid of a straight part (e.g., circles, semicircles and arcs; see Figure 5), all of Davida’s errors were the result of rotations and mirror reflections of the object in a coordinate frame composed of an axis aligned on the object’s axis of symmetry and/or a perpendicular axis intersecting it precisely at the shape’s centroid (Appendix 5.9 – 5.11; Figure 5 for examples; Movie S17). To account for these errors, we are thus required to assume that at some stage in the visual system object shapes are represented in a coordinate frame composed of orthogonal axes, aligned and centered onto the most elongated segment of elongated shapes and, for symmetrical shapes, aligned to their axis of symmetry and centered on their centroid– the S-Object-centered representation.

**Figure 4.**
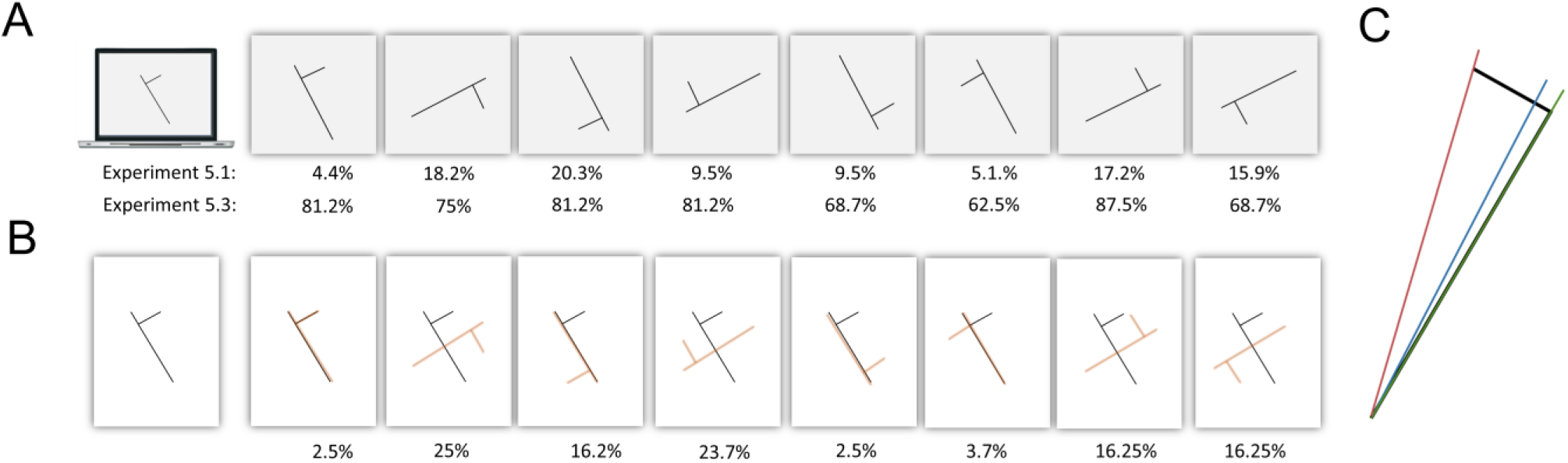
(A) Shown asymmetrical shapes in different orientations (here tilted 30 degrees counterclockwise from the vertical) and asked to draw on a separate sheet of paper either the most likely orientation of that shape given what she perceives (Experiment 5.1; see also Movie S13) or all the orientations of that shape that she perceives (Experiment 5.3; see also Movie S15), Davida systematically made the same 7 types of errors, whose proportions are reported here in percentages. (B) Shown asymmetrical shapes tilted 15 degrees from the vertical or horizontal on a sheet of paper and asked to trace the shape with ink, Davida made the same 7 types of errors, whose proportions are reported here in percentages. She made no other type of errors (Appendix 5.4. see also Movie S16). (C) A tilted asymmetrical elongated shape (in black ink), together with the axis corresponding to the shape’s longest straight segment (in green ink), the axis relating the two most distant points of the shape (longest span axis; in red ink) and the axis that minimizes the sum of squared distances to all points of the shape (the axis of least second moment; in blue ink).

**Figure 5.**
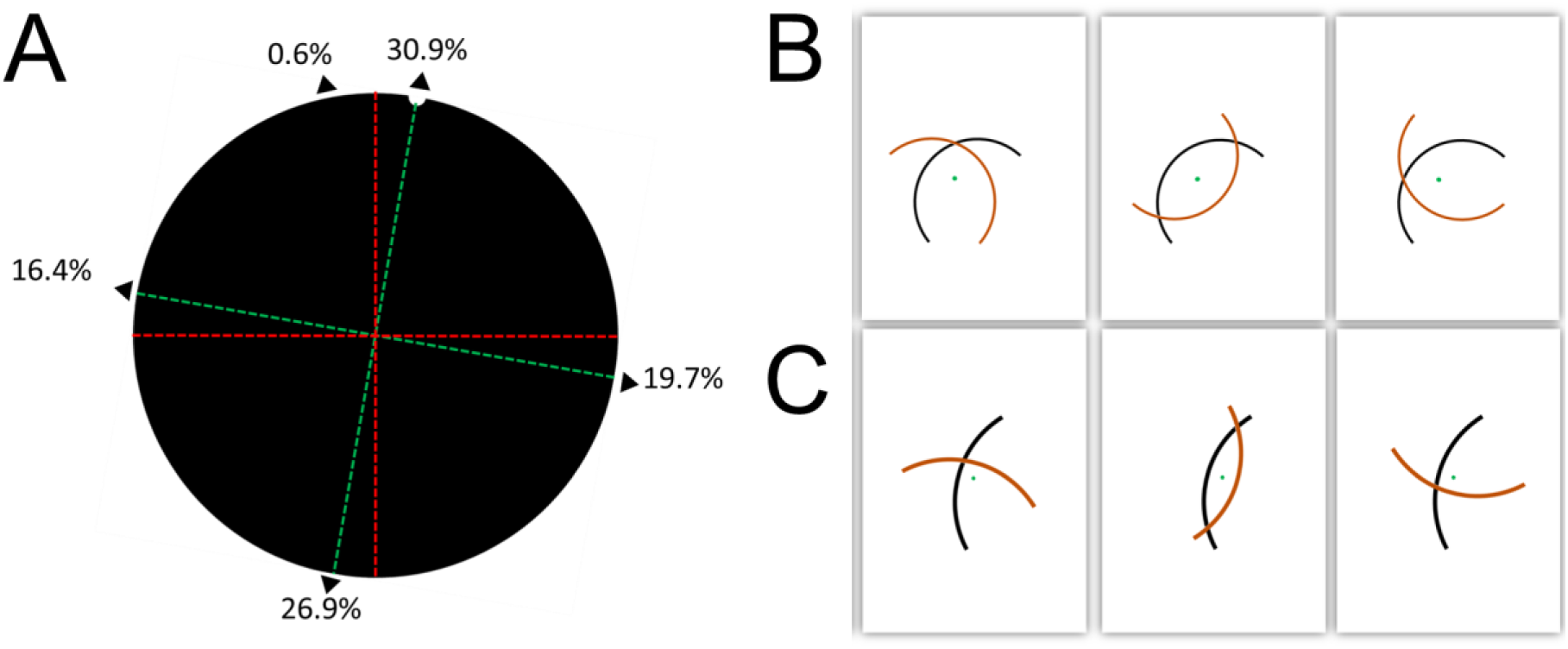
(A) Shown a large black disk with a small semicircular indent of the same color as the background for as long as needed, and asked to use the computer mouse to move a small round cursor and click as precisely as possible on the place(s) “where she sees the indent”, her errors consisted mainly in localizing the indent erroneously at less than 1 degree of visual angle from where it would have been if the disk had been rotated by 90 (19.7%), 180 (26.9%) or 270 (16.4%) degrees (Movie S17, online, is a recording of Davida performing this task). The green dotted lines, not shown during the experiment, illustrate a putative shape-centered representational frame composed of an axis aligned on the disk’s axis of symmetry, engendered by even a minimal deformation of a perfectly circular shape, and of its perpendicular. The red dotted lines, not shown during the experiment, illustrate another putative shape-centered representational frame composed of extrinsic vertical and horizontal axes (in red). The distribution of errors in this experiment clearly resulted from transformations of the stimulus within a frame intrinsic to the object (in green ink). (B, C). Illustrations of a semicircle (B) and of an arc (C) stimulus used in Experiment 5.10 (in black ink), of their centroid (green dot, not shown during the experiment), and of Davida’s different types of responses (in red ink) when shown these stimuli on a sheet of paper and asked to trace the shape with ink. Her errors consisted almost exclusively (98.5%) of shapes that were rotated by 90, 180 or 270 degrees around stimulus’ centroid (as illustrated on the Figure).

**Figure 6.**
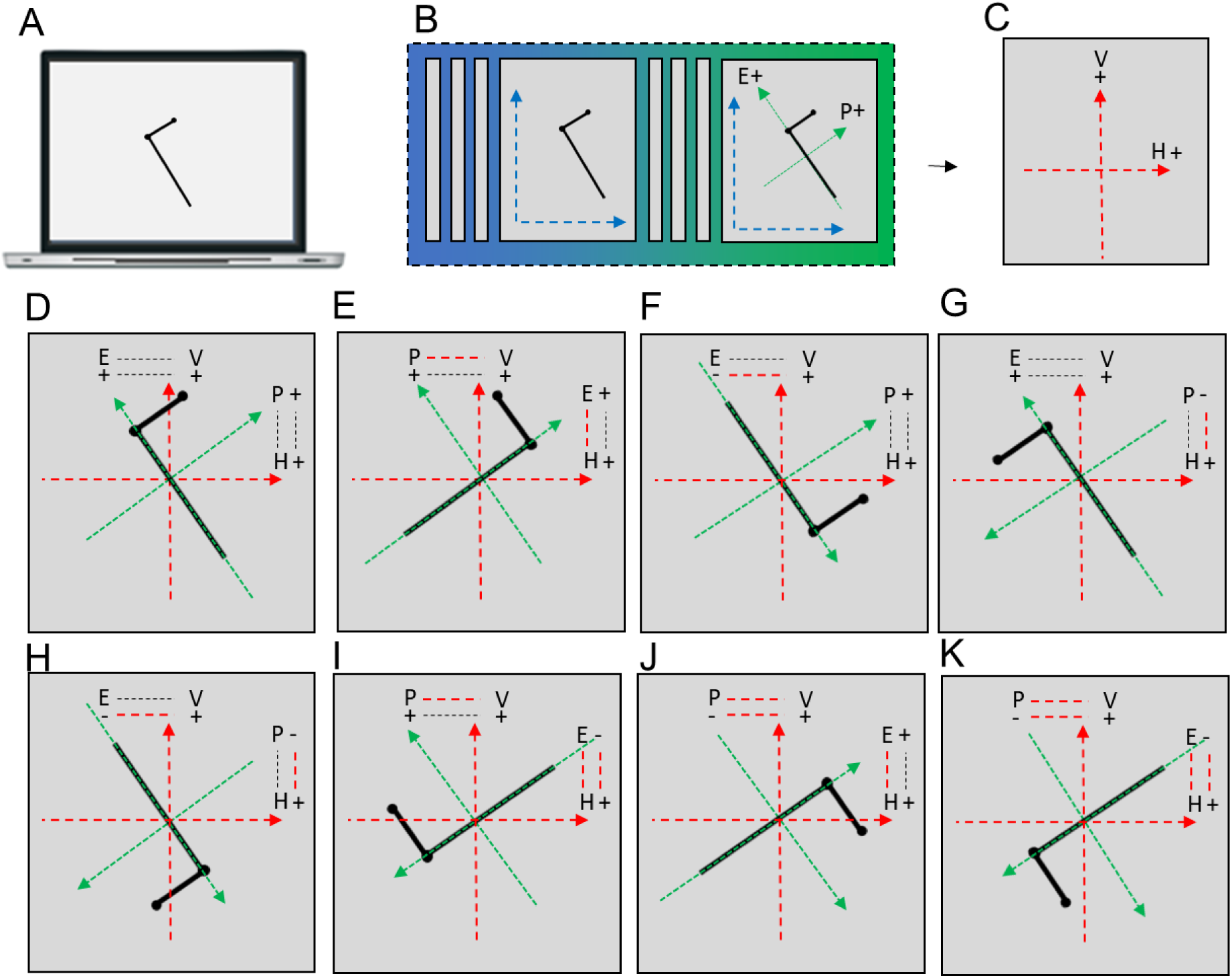
Illustration and interpretation of Davida’s 7 different types of errors with tilted, asymmetrical, elongated shapes in terms of a mapping deficit between an S-object-centered representation and a higher-order frame. **A**. A tilted, asymmetrical, elongated shape target. **B**. Schematic representation of the emergence, from the earliest cortical representation (blue), of an S-object-centered coordinate system (green) composed of a polar axis aligned with the object’s elongation axis (E) and a perpendicular polar axis crossing the shape through the center of its longest straight segment (Perpendicular axis; P). **C.** Schematic representation of a hypothetical higher order representational frame (red) composed of a polar vertical axis (V) and a polar horizontal axis (H). **D**. Illustration of the parameters specifying the relation between the two frames during a successful mapping process (20): their axis correspondence (in dotted lines: the shape’s elongation axis is related to the extrinsic vertical axis) and axis polarity correspondence (in dotted lines: the positive ends of the objects’ E and P axes are related to the positive ends of the scene-based V and H axes, respectively). **E-K**. Illustration and interpretation of Davida’s 7 different types of errors with this type of stimuli. The parameter(s) misrepresented during the mapping process are indicated by dotted lines in red ink. **E.** An error resulting from a misrepresentation of the axis correspondence: the object’s E and P axes are represented with respect to the wrong extrinsic axis. **F-H**. Errors resulting from a misrepresentation of the correspondence between the polarity of the objects’ E axis (**F**), P axis (**G**) or both (**H**) and the polarity of the extrinsic frame to which they relate. **I-K**. Combinations of an axis correspondence error and an axis polarity correspondence error concerning the objects’ E axis (**I**), P axis (**J**) or both (**K**).

The hypothesis that at one or several stage(s) of processing the primate visual system represents “objects” with respect to their own coordinate system is not new (Caramazza & Hillis, 1990; Driver, Baylis, Goodrich, & Rafal, 1994; Marr & Nishihara, 1978; McCloskey, 2009; McCloskey et al., 2006; Olson, 2003; Subbiah & Caramazza, 2000; Tipper & Behrmann, 1996), including the specific hypotheses that one axis is aligned on the most elongated part of elongated objects (Chaisilprungraung, German, & McCloskey, 2019; Gregory & McCloskey, 2010; Marr & Nishihara, 1978) or on the axis of symmetry of symmetrical objects (Palmer, 1985), and although they have been contested by some (Driver & Pouget, 2000; Mozer, 2002), there are numerous observations that have been interpreted as pointing to the existence of object-centered representations. Neurophysiological recording studies in monkeys have shown that some neurons in the supplementary eye field respond selectively to particular locations within a reference object (Olson, 2003); and, ventral-temporal object-responsive areas have been shown to compute representations of objects that are increasingly independent of their position, size and orientation in any coordinates (Logothetis, Pauls, & Poggio, 1995; Pasupathy & Connor, 2001; Rollenhagen & Olson, 2000). Behavioral studies have reported that humans tend to confuse objects’ orientations resulting from reflections across object-axes (Chaisilprungraung et al., 2019).

Furthermore, neuropsychological studies have reported brain-damaged patients who suffered from an attentional disorder whereby they ignore one half of a stimulus independently of its egocentric location or orientation (Tipper & Behrmann, 1996), one half of a stimulus separated in two parts by a gravitational vertical axis (Gainotti, Messerli, & Tissot, 1972), or one half of a stimulus separated in two by the stimulus’s own elongation axis (Driver et al., 1994). However, the different construals of “object” in these studies, the geometric properties of the center of the coordinate frame, and the corresponding stages of processing in the visual system, have remained largely underspecified.

The results in sections §3-§5 allow clear conclusions about the exact form of S-object-centered representations, their functional role, and their locus in the visual system: there is a stage of processing in the visual system, preliminary to the transformation of visual information in the different types of “higher” representational frames (e.g., spatiotopic, body-centered) underlying conscious visual perception, action and object recognition, which represents sharp-edged high contrast bounded areas of the visual field independently of their background and of each other in a perceptual frame composed of orthogonal axes, aligned on either the shape’s most elongated part or on the shape’s axis of symmetry, and centered either at the center of the shape’s most elongated part or on the shape’s centroid – the S-Object-centered representation.

The selectivity of Davida’s types of errors imposes a further constraint on our understanding of the nature and functional organization of the mechanisms involved in mapping S-object-centered representations onto higher frames. The existence of S-Object-centered representations implies that perceiving the orientation of a shape requires specifying the relation of that representation to “higher” representational frames. This entails specifying four parameters necessary for coordinates matching (McCloskey et al., 2006), Figure 6): (1) which coordinate frame axes relate to each other (axis correspondence); (2) the axes polarities correspondences (polarity correspondence); (3) the angular disparity between the axes (tilt magnitude) and (4) the direction of the tilt (tilt direction). Davida’s 7 types of errors can be interpreted in this framework as a consequence of a specific failure of the mechanisms that specify the axis correspondence and axis polarity correspondence between the two frames, leading to axis correspondence errors (Figure 6 E), axis polarity correspondence errors (Figure 6 F-H) and their combination (Figure 6 I-K). In line with this componential view, previous studies of brain-damaged humans and monkeys have reported cases showing disproportionate difficulties either in discriminating mirror images and 90 degrees rotations of objects (Davidoff & Warrington, 1999; Davidoff & Warrington, 2001; Eacott & Gaffan, 1991; Harris et al., 2001; Martinaud et al., 2014; Priftis et al., 2003; Turnbull et al., 1996; Turnbull & McCarthy, 1996) or two versions of the same shape rotated by a few degrees (Cowey & Gross, 1970; Holmes & Gross, 1984).

[§6] Davida’s selective difficulty in perceiving the orientation of the type of 2D stimuli used in the experiments reported so far – sharp-edged, stationary, defined by high luminance contrast from the background – contrasted with otherwise normal perception of the physical environment. Unlike the stimuli used in the experiments reported so far, physical environments under naturalistic viewing conditions are dynamic scenes populated with 3D objects of lower contrast separated by edges that are often blurred or shaded (Sebastian, Burge, & Geisler, 2015). To delineate more precisely this dissociation, we explored the influence of movement, contrast (chromatic and luminance), blur and depth on Davida’s performance. Davida had severe difficulty with isoluminant stimuli (Appendix 6.1, 6.2) but her performance improved and often became flawless when the stimuli were defined by very low luminance contrast with the background (Figure 7 A – B; Appendix 6.3 – 6.9, Movies S18-20), when the stimuli were implied by motion (Appendix 6.10), when the stimuli were strongly blurred (Figure 7 C – D; Appendix 6.11 – 6.14, Movies S21-23) or when stimuli were shown in 3D (Appendix 6.15). Interestingly, her performance worsened (normalized) when presented with low luminance contrast stimuli in the visual illusion task in which perception of accurate orientation hinders performance (the Ponzo illusion; Appendix 6.7).

**Figure 7.**
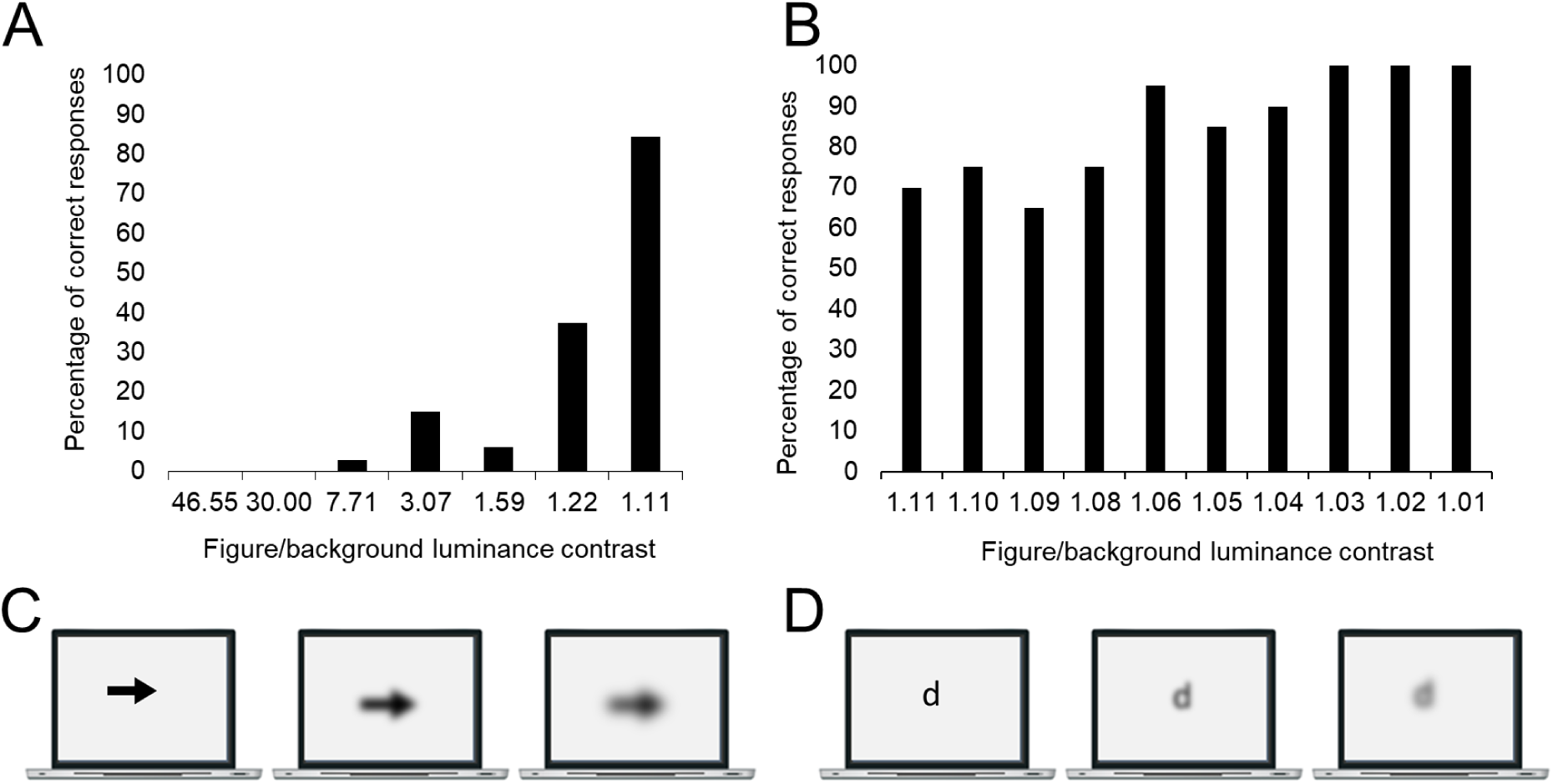
(A, B) Davida’s percentage of correct responses when shown arrows pointing up, down, left or right (randomly) and asked to indicate the orientation of these arrows by pressing on the corresponding key on a computer keyboard for arrows of different levels of luminance contrast with the background (Appendix 6.3, see also Appendix 6.4 - 6.9 and Movies S18 – 20). C. When shown arrows of 3 different levels of gaussian blur (0, 50, 80) pointing up, down, left or right and asked to indicate the orientation of these arrows Davida produced 0% correct responses at the two lowest levels of blur and 100% correct responses at the highest level (Appendix 6.11, see also Appendix 6.12 and Movie S21). D. When shown orientation sensitive letters (b, d, p, q) edited with different levels of gaussian blur (0, 50, 80), Davida read 2/40 and 0/40 letters accurately at the two lowest levels of blur but she read all the letters correctly (40/40) when the they were edited with a gaussian blur with a radius of 80 pixels. The Movie S22 illustrates Davida’s performance in this type of experiment.

The dissociation between stimulus properties affecting Davida’s performance seems to parallel that between the information that the two main subcortical channels carry from the retina to the primary visual cortex: the chromatic parvocellular (P) channel is specialized for processing sharp-edged, fine, stationary, and high-contrast stimuli, such as those impaired in Davida; whereas the achromatic magnocellular (M) channel is mostly sensitive to stimuli with complementary characteristics – 3D, coarse, large, moving, brief, and low contrast (Livingstone & Hubel, 1987; Merigan & Maunsell, 1993) – which were clearly spared in Davida. Thus, Davida’s disorder appears to result from a specific deficit in setting the axis correspondence and axis polarity correspondence between a correctly computed S-Object-centered representation computed from information derived, at least largely, from the parvocellular channel and an extrinsic representational frame. This finding implies that conscious visual perception of objects is the result of the integration of independent parallel mappings of shape-centered representations computed from information derived from the parvocellular and magnocellular channels into higher frames (e.g., spatiotopic) (see Figure 1).

This interpretation may seem surprising given the documented considerable mixing of the M- and P- channels in the primary visual cortex (Merigan & Maunsell, 1993; Nassi & Callaway, 2009; Sincich & Horton, 2005) and the popular view that the two channels lack independent contribution to vision beyond the primary visual cortex (Sincich & Horton, 2005). However, there is evidence that information derived from the M and P subcortical channels (e.g., the processing of color vs. luminance, sharp vs. blurred edges) remains segregated at least to some degree in several areas of the extrastriate cortex such as V2, V3, V4 and MT (Bushnell, Harding, Kosai, Bair, & Pasupathy, 2011; Ferrera, Nealey, & Maunsell, 1994; Merigan & Maunsell, 1993; Nassi & Callaway, 2009; Oleskiw, Nowack, & Pasupathy, 2018; Tanigawa, Lu, & Roe, 2010; Tootell & Nasr, 2017; Yabuta, Sawatari, & Callaway, 2001). Additional evidence that the information derived from the M and P channels remains at least partly segregated comes from two previous neuropsychological cases who suffered from a significantly more severe disorder in perceiving the orientation and location of stimuli biased toward the P- than the M- channel (McCloskey et al., 1995; Pflugshaupt et al., 2007). McCloskey and colleagues (McCloskey, 2004, 2009; McCloskey et al., 1995), in particular, reported an individual, A.H., whose severe disorder in perceiving objects’ location and orientation was modulated by the visual features of the stimuli. For instance, she was better at judging the orientation of arrowheads presented for 50 msec (94%) or with a low contrast difference with the background (72%) than those presented at high contrast for an unlimited amount of time (33-36%); she was severely impaired at pointing to a stationary visual stimulus (an “X” or an “O”; 59%) but flawless when the same stimulus oscillated up and down (6°) at 1 Hz; and, she was also better at copying shapes and at reading letters flickering at 25 or 10 Hz (100%) than when presented continuously (36% and 80%). Those observations led the authors to propose that objects’ location and orientation are jointly computed in separate M-based transient and P-based sustained visual subsystems (McCloskey, 2009). The findings reported here additionally allow specifying a locus of integration of shape-centered representations derived from the M- and P- channels precisely to the stage of processing where the visual system maps S-Object-centered representations onto extrinsic frames of reference resulting in Perceptual Objects (Figure 1).

### Discussion

Davida has a particularly clear and highly selective visual disorder: she perceives any 2D sharp-edged high-contrast bounded region of space alternating between its correct orientation and all other orientations that would result from a failure to specify the correct axis correspondence and/or axis polarity correspondence in the course of mapping an S-object-centered representation – composed of orthogonal axes aligned on either the shape’s most elongated part or on the shape’s axis of symmetry and centered either at the center of the shape’s most elongated part or on the shape’s centroid – onto higher coordinate frames (e.g., spatiotopic, body-centered; see Figure 6). To cope with this longstanding disorder, Davida early on developed her own “personal font” in which orientation-sensitive letters are attributed different shapes (Appendix Figure S8). Other particularly telling consequences of her disorder are (1) that Davida typically locates the tip of a straight arrow almost exactly where it would be located if that arrow was rotated by 90 degrees around the geometric center of its elongation axis (e.g., Appendix 1.7, 5.7 and 5.8) and the tip of a curved arrow almost exactly where it would be if the arrow was mirrored across its axis of symmetry or rotated by 90 degrees around its centroid (Appendix 5.11); (2) that Davida sometimes perceives non-overlapping objects as if they were overlapping (e.g., Appendix 3.5.), which often hinders the perception of objects located in an area that would be covered by another object if that object were rotated by 90 degrees or inverted (e.g., Appendix 3.6. and 3.7); (3) that Davida performs far better than control participants in tasks, such as visual illusions and stimulus-response compatibility tasks, in which accurate orientation perception typically hinders performance (see Appendix 1.11-1.13).

In contrast to the set of visual properties that determine Davida’s perceptual deficit, she has intact perception of the shape, size, location, distance, tilt, and movement of the same 2D stimuli, and of the orientation of shapes that are either strongly blurred, defined by very low luminance contrast with the background, implied by motion, shown in 3D or that emerge from a collection of non-connected elements.

This highly selective visual disorder forces three main conclusions about the nature of the mechanisms involved in transforming retinotopically represented visual primitives into conscious perception of objects in environmental coordinates (Figure 1): (1) There is an unconscious stage of processing where the visual system represents each sharp-edged bounded area in the visual field in their own “shape-centered” perceptual frame composed of orthogonal axes aligned on either the shape’s most elongated part or on the shape’s axis of symmetry, and centered either at the center of the shape’s most elongated part or at the shape’s centroid. We refer to this new type of visual representation as “Shape-Object-centered representation” to distinguish it from conscious object representations (Perceptual-Object representation). (2) S-Object-centered representations of objects characterized by different visual features (e.g., sharp-edges vs blurred) derived from the properties of the subcortical M- and P- channels are computed in parallel and integrated precisely at the level at which such object representations must be mapped onto higher frames of reference representing Perceptual Objects. (3) This mapping involves computing several parameters (McCloskey et al., 2006) of which at least two (axis correspondence and axis polarity correspondence) are independent and receive separate inputs from representations computed in the Parvocellular and Magnocellular visual streams.

These findings corroborate and complement previous proposals regarding the nature of object-centered coordinate frames (Driver et al., 1994; Marr & Nishihara, 1978; Olson, 2003; Quinlan & Humphreys, 1993; Sekuler, 1996; Sekuler & Swimmer, 2000; Subbiah & Caramazza, 2000; Tipper & Behrmann, 1996), the segregation of processing of information derived from the M- and P- channels in mid-level vision (Bushnell et al., 2011; Flanagan, Cavanagh, & Favreau, 1990; Livingstone & Hubel, 1987; McCloskey et al., 1995; Pflugshaupt et al., 2007; Tanigawa et al., 2010; Tootell & Nasr, 2017), and the division of labor within the visual system among the processes involved in different aspects of orientation processing (Clifford, Spehar, Solomon, Martin, & Zaidi, 2003; Eacott & Gaffan, 1991; Goodale, Milner, Jakobson, & Carey, 1991; Holmes & Gross, 1984; McCloskey et al., 2006; Valtonen et al., 2008), and tie these proposals to a particularly clear level of representation within the visual system where objects defined strictly by bounded regions of space (or “uniform connected regions”; Palmer & Rock 1994) are processed in parallel.

The neural correlates of the different mechanisms described in this proposal remain unclear. Nevertheless, the concept of duplicate P- and M-derived Shape-Object-centered representations is broadly consistent with several known properties of the visual area referred to as LO1-LO2 in humans (Kolster, Peeters, & Orban, 2010; Larsson & Heeger, 2006) and V4d in monkeys (Roe et al., 2012). First, and in line with our definition of S-object-centered representations, V4d is assumed to play an important role in figure-ground segmentation through the detection of discontinuities of color and/or luminance, and to encode isolated shapes’ (i.e., bounded regions of space) boundary features in an object-centered frame of reference (Kim, Bair, & Pasupathy, 2019; Pasupathy, 2015; Roe et al., 2012). Second, and in line with the hypothesis that duplicate S-object-centered representations are computed from information derived from the M-cellular and P-cellular visual channels, V4d/LO1-LO2 receive similarly potent inputs from P- and M-neurons (Ferrera et al., 1994) but retains some degree of segregation between clusters of neurons specialized in the processing of information derived from the P- and M- channels (Ferrera et al., 1994), such as between the processing of shapes with sharp or blurred edges (Oleskiw et al., 2018) or defined by color or luminance (Bushnell et al., 2011; Tanigawa et al., 2010; Tootell & Nasr, 2017). It is so far unknown whether the P-dominated sites within V4 are are also those specialized in the processing of color and sharp-edges and whether the M-dominated sites are those specialized for luminance and blur, but our general hypothesis suggests that it could be the case. Third, LO1-LO2 (or V4d in monkeys) are situated at an intermediate position between the early retinotopic representation characterizing V1–V3 and more abstract non-retinotopic object representations in LO/IT (McKyton & Zohary, 2007; Vernon, Gouws, Lawrence, Wade, & Morland, 2016). In fact, there is evidence that a retinotopic-to-non-retinotopic transition may occur near or within LO-1 and LO-2. Studies of monkeys’ V4d have reported the coexistence of a retinotopic topographical organization (Roe et al., 2012) and some degree of tolerance to changes in retinotopic location (Gallant, Braun, & Van Essen, 1993; Gallant et al., 1996; Pasupathy & Connor, 2001; Rust & DiCarlo, 2010) and size (El-Shamayleh & Pasupathy, 2016) of the stimulus suggesting a coding of objects’ shapes in object-based coordinates. Studies in humans have further documented a shift between LO1, where retinotopy and shape-centered representations seem to coexist, and LO2, which seems to encode object-centered shape representations only (Vernon et al., 2016). Finally, it may be worth mentioning that the hypothesis that remapping retinotopic to spatiotopic representations is limited to objects with high attention priority (Burr & Morrone, 2011) is compatible with V4d’s known role in visual attention and selection (Roe et al., 2012).

Whether and if so how LO1-LO2-V4d is involved in the mapping of S-object-centered representations onto higher frames also remains unclear. There is some evidence that in humans LO1 plays a role in the ability to discriminate the orientation of gratings tilted a few degrees from each other (Silson et al., 2013) and that in monkeys IT is necessary for discriminating tilted shapes (for instance 30 or 45 degrees apart; Gross, 1978; Holmes & Gross, 1984), suggesting a role in at least the tilt component of orientation representation. That the well-documented patient D.F. (Goodale et al., 1991), who has a bilateral lateral occipital cortex (LO) lesion, could grasp objects accurately, additionally suggests that tilt information for action is computed independently in the dorsal stream. Whether ventral stream regions are also responsible for the computations underlying axis correspondence and axis polarity correspondence is less clear. To our knowledge, the impact of lesions in LO1-LO2 on these aspects of orientation processing has never been tested in humans, and the type of IT lesions that affect monkeys’ tilt discrimination does not impact their ability to discriminate stimuli differing from one another in terms of axis correspondence or axis polarity correspondence (Gross, 1978; Holmes & Gross, 1984). In contrast, damage to the visual dorsal stream has been found to affect both monkeys’ (Eacott & Gaffan, 1991) and human patients’ ability to discriminate mirror images of objects (a condition termed “mirror agnosia”; Davidoff & Warrington 1999, 2001; Turnbull & McCarthy 1996; Priftis et al., 2003; Turnbull et al., 1997; Martinaud et al., 2014; Harris et al., 2001; Vinckier et al., 2006). Altogether, these observations invite three inferences: (1) tilt is computed in both the ventral stream (for visual perception) and the dorsal stream (for actions); (2) the dorsal stream cortex critically contributes to axis correspondence and axis polarity correspondence; (3) thus, computing axis correspondence and axis polarity correspondence for the mapping of S-object-centered representations onto higher frames may require a dorsal-to-ventral flow of information. This interpretation would be in line with the longstanding hypothesis that the dorsal stream plays a critical role in spatial vision and coordinates matching (Colby, 1998; Duhamel, Colby, & Goldberg, 1992; Olson, 2003).

In conclusion, Davida’s pattern of spared and impaired performance provides clear constraints to plausible answers to longstanding questions about the nature of the processes that result in conscious perception of the world from the 2D image captured by the retina. These include evidence that the primary units of shape information in defining objects consist of bounded regions of space akin to Palmer and Rock’s (Palmer & Rock, 1994; see also Tse & Palmer, 2012) “uniform connected regions”, which are represented in a shape-centered coordinate frame computed from information derived from the P- subcortical channel and in another shape-centered coordinate frame computed from information derived from the M- subcortical channel. We have also provided evidence that the coordinate systems for S-Object-centered representations are aligned and centered on the longest straight part of elongated shapes and on the axis of symmetry and centroid of non-elongated symmetrical shapes. Finally, the evidence from Davida’s performance invites the conclusion that conscious visual experience results from the mapping of these P- and M- derived S-object-centered representations onto behaviorally relevant frames.

## Acknowledgments

We thank Rick Born, Patrick Cavanagh, Bevil Conway, Jack Gallant, Mel Goodale, Michael McCloskey and James Pomerantz for their helpful suggestions, Eric Falke for referring Davida to us for further study, Sarah Carneiro for collecting part of the data from control participants, and Davida and her family for their time, motivation and kindness. This research was supported by the Mind, Brain and Behavior Interfaculty Initiative provostial funds.

## Author contributions

G.V administered the experiments and analyzed the data. The three authors conceptualized the study and wrote the manuscript.

## APPENDIX

**This APPENDIX file includes:**

Supplementary text

Figures S1 to S46

Tables S1 to S4

Legends for Movies S1 to S23

**Other supplementary materials for this manuscript include the following:**

Movies S1 to S23, available online on the Open Science Framework platform (link: https://osf.io/pf56m/?view_only=bda3dcc0b9ea4d62ac122e23d8227463).

### Appendix

#### Case history

Davida is a right-handed (Oldfield’s Laterality Index of 80), athletic (she is skillful at many sports, including soccer and basketball) and very cooperative young woman. She was 15 years old when this study began in October 2016 and 17 years old when it ended in March 2019. Information regarding her early history was obtained from her parents through a developmental and family history questionnaire and by reviewing her medical record.

#### Medical and developmental history

Davida was born at 33.5 weeks by cesarean section and was twin B of a twin pregnancy. The pregnancy was notable for preeclampsia. The cesarean section occurred after premature rupture of the membranes of Twin A and an increased fetal heart rate. Davida was 2,385 grams at birth (normal for gestational age), was immediately alert and active, and had Apgar scores of 8 and 9 at 1 and 5 minutes, respectively. Davida stayed in the hospital for 4 weeks after birth to gain weight. She required nasal CPAP support for a few days after delivery, but the newborn period was otherwise uneventful. Early developmental milestones were achieved within the usual time frames and she revealed no other medical or developmental issues. Although Davida tested normal in vision and hearing exams, she struggled with reading starting in the 2nd grade. She achieved partial compensation for her reading difficulties though a strong work ethic and extensive support through private tutoring and other programs.

#### Neurological history

In December 2014 she had a normal neurological examination. A 1.5T brain MRI conducted in late 2015 revealed a normal brain with no evidence of cortical malformations or early injury. An electroencephalogram from early 2016 at rest and with photic stimulation was normal.

#### Psychiatric history

There were no issues to report in the psychiatric history. A psychological evaluation of her social and emotional functioning carried out in May 2014 using the Behavior Assessment System for Children, second Edition(Sandoval & Echandia, 1994) (BASC-2), found average scores in all scales, suggesting that there is no problematic anxiety, depression, sense of inadequacy, somatization, inattention or hyperactivity, and that her relationship with her parents, her interpersonal relations, self-esteem, locus of control and self-reliance were normal.

#### Neuropsychological history

A first evaluations performed in May 2014 concluded that Davida displayed: (1) average or above average performance on all the subtests of the WISC-IV ((Vaughn-Blount et al., 2011); all tests 37 < percentile < 84); (2) typical verbal (percentile 34) and visual (percentile 27) memory on the Wide Range Assessment of Memory and Learning-2 (Sheslow & Adams, 2003); (3) no clinically relevant executive function deficits on the Behavior Rating Inventory of Executive Function(Gioia, Isquith, Guy, & Kenworthy, 2000) (all Teacher’s and Parents’ T-scores < 65) or on the Tower test of the Delis-Kaplan Executive Function System(Delis, Kaplan, & Kramer, 2001) (Percentile 37); and (4) no other clinically relevant behavioral abnormalities on the Conners’ 3-T rating scale(Conners, 2010), other than the presence of learning problems (Teacher’s and Parents’ T-scores > 80; all other scales Teacher’s and Parents’ T-scores < 55).

A second evaluation, in June 2014, which focused on possible underpinnings of her reading difficulties, found: (1) intact phonological awareness and meta-phonological skills at the CTOPP-2(Wagner, Torgesen, Rashotte, & Pearson, 1999) (Phonological Awareness Index, Elision, Blending Words and Phoneme Isolation tests, all 16 < Percentile < 92); (2) below average speed on rapid naming of letters, numbers (she was at Percentile 1 or below at the Rapid Letter and Rapid Digit Naming tests of the CTOPP-2) and words (she was at the percentile 1 for her age on the TOWRE-2(Torgesen, Wagner, & Rashotte, 2012)); (3) excellent reading comprehension (percentile 95th on the untimed Gray Diagnostic Reading Test(Bryant, Wiederholt, & Bryant, 1991)), thus suggesting that, despite her lack of fluency, she was able to accurately extract meaning from text; (4) age-appropriate listening comprehension on the Understanding Spoken Paragraphs subtest of the CELF-5(Wiig, Semel, & Secord, 2003); and (5) strong abstract thinking skills on the D-KEFS 20(Delis et al., 2001). Davida had an atypical performance on the Test of Variables of Attention(McCarney & Greenberg, 1990), characterized by slow and inconsistent response times and a fast decline in her ability to inhibit incorrect responses over the time of testing, but the interpretation of these results is difficult given her visual disorder (see main text).

#### Ophthalmological history

The ophthalmological history was negative. An evaluation carried out in May 2017 revealed: (1) A normal pupillary exam without significant anisocoria or relative afferent pupillary defect; (2) Full eye movements with the patient’s being orthotropic in all directions of gaze at both distance and near. Smooth pursuit and horizontal and vertical saccades were intact; (3) An unremarkable anterior segment; (4) A dilated fundus with pink, sharp, normal, healthy-appearing optic nerves with 0.25 cup-to-disc ratio in each eye, flat healthy-appearing maculae with good foveal reflexes bilaterally, and a normal periphery; (5) Normal cyclopegic refraction with approximately plano in each eye. (6) Bilaterally normal Optical Coherence Tomography of the macula and the retinal nerve fiber layer; (7) Visual acuity of 20/15 binocularly, and at least 20/20 for each eye individually with the Snellen chart and a green background. During the test, she was not able to read the letters but instead described them. When asked to explain how many lines she sees in the letter “H” for instance, she replied “three lines”. When asked about their orientation, she replied “two horizontal and one vertical line”; (8) A normal color perception assessed on the Ishihara Test(Ishihara, 1987) and on the Farnsworth D-15 color test; (9) Normal contrast sensitivity measured with the Pelli-Robson contrast sensitivity test(Pelli & Robson, 1988). On the Pelli-Robson, she was asked to copy the letters, which she did, but in the wrong orientation; (10) Abnormal stereopsis: She demonstrated no stereopsis on Titmus Vision testing and on the Worth 4 dots test. On a version of the Worth 4 dots test, for instance, Davida was asked to wear red-green glasses and to look at series of dots on a series of screens. There were always 1 or 2 green, 1 or 2 red and 1 or 2 black dots. Davida always reported the same experience—that of seeing stable black dots and rapidly flashing red and green dots in asynchrony. She would typically say “I see green-red-green-red”. Interestingly, she also reported the white background of the computer screen to flicker from green to red. This suggests a rapidly alternating ocular suppression.

#### 1. Materials and methods: set of results §1

##### 1.1. Arrow orientation judgment task

Davida was shown 80 arrows pointing up, down, left or right and asked to indicate the orientation of these arrows verbally (right, left, up or down) and simultaneously pointing with her finger in the same directions. These arrows were black and large (10 degrees of visual angle), and were displayed one at the time, at the center of the screen on white background, for as long as needed. Davida almost systematically (95% of the trials) responded as if the arrow pointed to 90 (20%), 180 (37.5%) or 270 (37.5%) clockwise degrees from the actual orientation.

##### 1.2. Arrow copy task

Davida was shown 20 black, large (10 degrees of visual angle) arrows pointing up, down, left or right and asked to “draw what she saw”, including multiple orientations if needed. The stimuli were displayed one at the time, at the center of the screen, on white background, for one second. Davida drew on average 3.1 differently oriented versions of each displayed arrows. Among these, she included the correct orientation in most of the trials (19/20 trials), but also the equivalent of the displayed arrow pointing 90, 180 or 270 clockwise degrees from the actual orientation in 10/20, 16/20 and 17/20 trials, respectively. An illustration of Davida’s copying of arrows is shown on Movie S2, online. See also Movie S1 online for a discussion of what she perceives.

##### 1.3. Object orientation decision task

Experiment 1: typical or atypical orientation? Davida was shown 40 line-drawings of objects from the Snodgrass and Vanderwart (1980) set once in their typical upright orientation and once upside-down. The stimuli were presented one at a time for as long as needed, and Davida was asked to decide whether each object was in its typical or atypical orientation. She responded incorrectly to 69/80 trials (35/40 errors for upright stimuli and 36/40 errors for inverted stimuli).

Experiment 2: typical or atypical orientation? Davida was shown 20 line-drawings of objects from the Snodgrass and Vanderwart (1980) set once in their typical upright orientation, once upside-down and once rotated by 90 degrees (10 objects clockwise and 10 objects counter-clockwise). The stimuli were displayed one at a time for as long as needed, and Davida was asked to decide whether each object was in its typical or atypical orientation. She responded incorrectly to 36/60 stimuli (11/20 errors for upright stimuli, 14/20 errors for inverted stimuli and 11/20 errors for rotated stimuli).

##### 1.4. Abstract shape copy

Davida was shown 50 different abstract shapes and asked to copy them as accurately as possible on a separate sheet of paper while the stimulus remained in view. She systematically copied the shapes as if the stimuli were inverted vertically, reversed horizontally, plane-rotated by 90 or 180 degrees (e.g., see Figure S1). See also Movie S1 online for a discussion of what she perceives when shown an abstract shape on the computer screen.

**Fig. S1.**
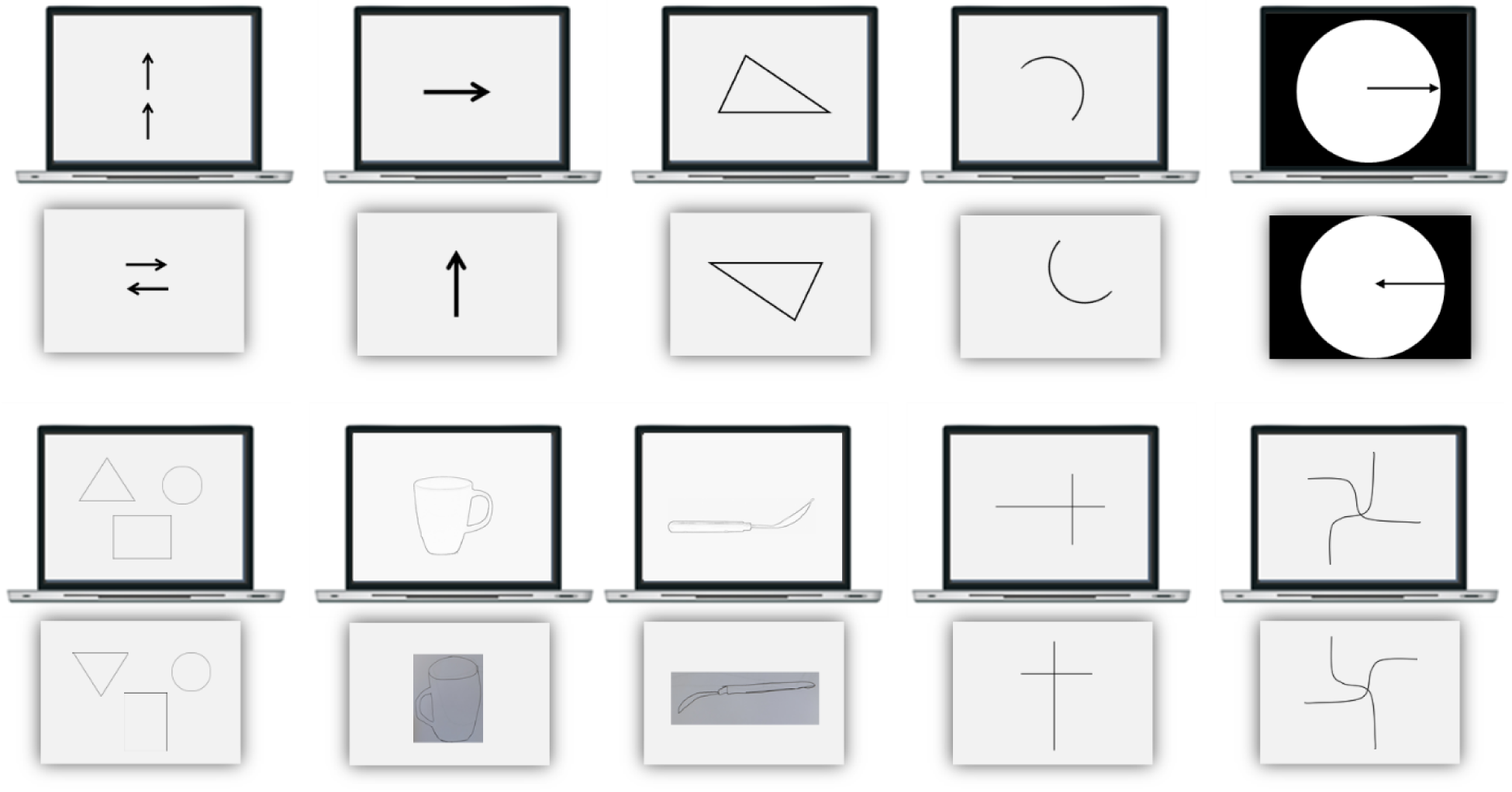
Examples of stimuli (displayed on the computer screen) and, below them, of Davida’s attempts to copy them as accurately as possible.

##### 1.5. Letters and words copy task

Davida was shown the letters b, d, p and q one at a time, 5 times each, and 6 short palindromes (mug, gum, live, evil, dog, god) once each and was asked to copy them as accurately as possible on a separate sheet of paper while the stimulus remained in view. Stimuli were presented one at the time at the center of the screen and were composed of large (± 5 degrees of vertical visual angle) lower-case black letters in Calibri font on white background. She systematically copied the single letters (Table S1) and the letters in the words as if they were inverted or rotated (see Figure 2B). An illustration of Davida’s copying of the single letters p, b, d and q is shown on Movie S3, online.

**Table S1.**
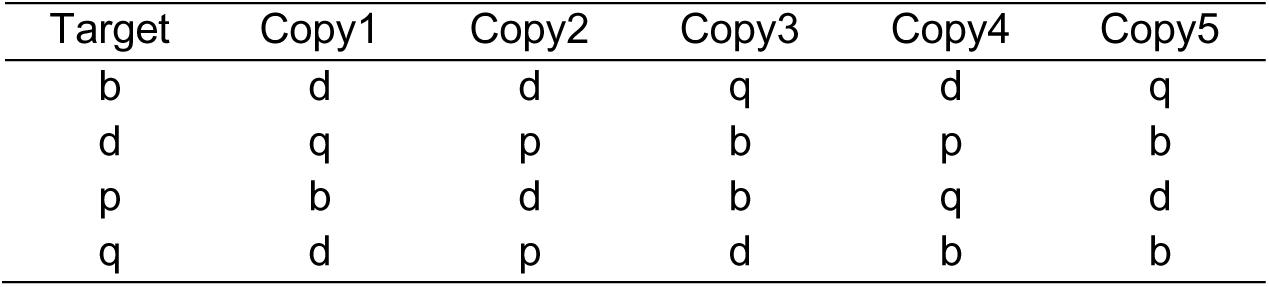
Davida’s errors in the 5 blocks of the letters copy task.

| Target | Copy1 | Copy2 | Copy3 | Copy4 | Copy5 |
| --- | --- | --- | --- | --- | --- |
| b | d | d | q | d | q |
| d | q | p | b | p | b |
| p | b | d | b | q | d |
| q | d | p | d | b | b |

##### 1.6. Letters, words and numbers reading task

Davida was asked to read five times the 26 letters of the alphabet, and, in a separate session, 40 short palindromes composed of 3 to 6 letters (e.g., mug, gum, live, evil). Stimuli were composed of large (± 10 degrees of vertical visual angle) lower-case black letters in ComicSans font on a white background. In both sessions, Davida sat approximately 60 cm from the computer screen and was asked to read as accurately as possible stimuli displayed in a random order, one at the time, at the center of the screen for as long as needed. The responses were encoded by the experimenter. The next trial was initiated by the experimenter after response encoding. In the letter reading task, Davida read without noticeable hesitation and without errors all the non-orientation-critical letters – letters that have a unique shape in the alphabet. However, she hesitated noticeably before naming all the orientation-critical letters – letters that differ from at least one other letter in the alphabet only by its orientation (b, d, p, q, n, u, c, z). She named correctly 15% of them, confused 75% of them with another letter of similar shape but different orientation and declined to respond 10% of the time (see table S2.). Davida was rather slow in reading the palindromes but read 77.5% of them (31/40) correctly: she read accurately all the 14 palindromes that contained no orientation-critical letter, all but one of the 18 containing at least one orientation-critical letter, but for which an orientation-sensitive letter confusion would have resulted in a non-word (she was unable to read “repaid”), but she made systematic errors in reading the 8 palindromes for which a letter confusion would result in plausible word (e.g., “deer” – “beer”, “doom” – “boom”, “peek” – “beek”, “raw” – “ram”). An illustration of Davida’s reading of the single letters p, b, d and q is shown on Movie S4, online.

**Table S2.** Davida’s errors in the 5 blocks of the letters reading task

| Target | Block1 | Block2 | Block3 | Block4 | Block5 |
| --- | --- | --- | --- | --- | --- |
| b | p | d | p | p | d |
| c | n |  |  |  |  |
| d | b | p | b | p | p |
| n | u | u | c | DNK |  |
| p | d | d | b | d | b |
| q | p | d | d | d | b |
| u | n | n | n | c | n |
| z | DNK | DNK | DNK | n |  |

On another task, Davida was asked to read the 10 digits and 20 2-digits numbers displayed one at a time at the center of the screen for as long as needed. The numbers were large (± 5 degrees of vertical visual angle) and drawn in black ComicSans on white background. Davida scored 22/30. Davida confused all the instances of the digits “6” and “9” (e.g., 68 « ninety-eight »), but made no other errors.

##### 1.7. Localizing the tip of an arrow by pointing with the computer mouse pointer

On a first task, a large (±10 x 6 degrees of visual angle) black arrow pointing left, right, up or down was displayed at the center of the computer screen for an unlimited duration. On each trial, Davida was asked to use the computer mouse to move a small round cursor and click as precisely as possible on the tip of the arrow. This task was presented multiple times for a total of 560 trials. As shown in Figure S2 and illustrated in the Movie S5, Davida almost systematically mislocated the position of the tip to approximately the place it would have been if the arrow were rotated by 90 degrees (38.8%), 180 degrees (19.3%) or 270 degrees (41.9%).

**Fig. S2.**
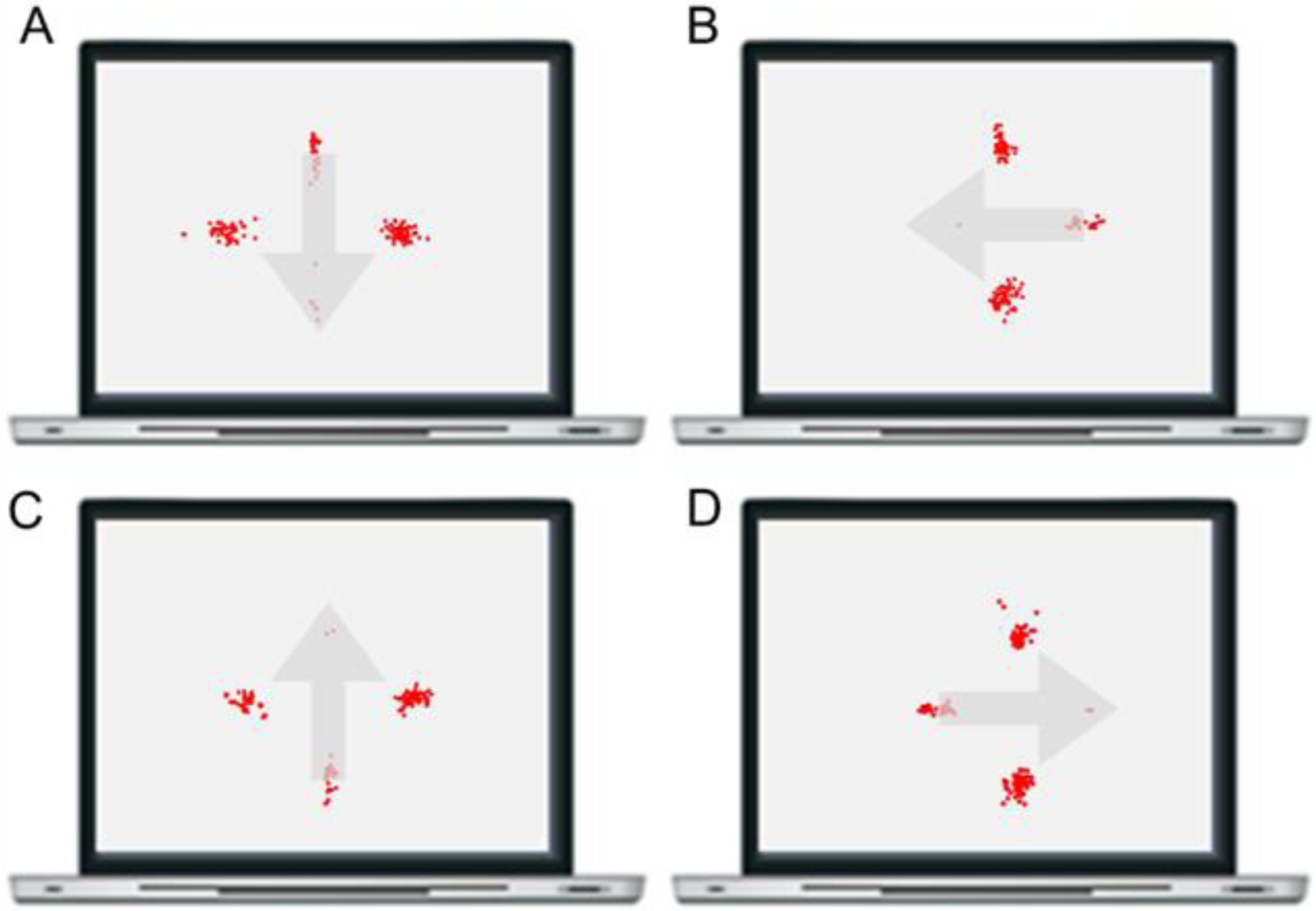
Each red dot represents the coordinates of one of Davida’s attempt at localizing the tip of an arrow pointing down (A), left (B), up (C) or right (D). Arrows appear in grey here for transparency, they were displayed in black during the experiment.

On a second task, a large (±10 x 6 degrees of visual angle) black arrow was displayed at the center of the computer screen for an unlimited duration and was either pointing upright (0 degrees) or at 20, 45, 70, 90, 110, 135, 160, 180, 200, 225, 250, 270, 290, 315 or 340 degrees clockwise. There were 10 trials for each 45 degrees steps (0, 45, 90, 135, 180, 225, 170, 315) and 5 trials for the other orientations for a total of 120 trials. On each trial, Davida was asked to use the computer mouse to move a small round cursor and click as precisely as possible on the tip of the arrow. Once again, as illustrated in the Movie S5 (part II), Davida almost systematically mislocated the position of the tip to approximately (+- 100 pixels) where it would have been if the arrow were rotated by 90 degrees (33.3%), 180 degrees (26.6%) or 270 degrees (30%).

##### 1.8. The color after-effect with striped stimuli

Davida was asked to look for 1 minutes at a red screen with black vertical stripes and then, following this induction period, to describe as precisely as possible what she saw when the screen turned white. She reported seeing “a grid made of white bars over blue”. The perceived white and blue are typical of color after-effects and correspond to approximatively the complementary of the colors perceived during the induction period (black and red). The “grid” perceived over the test screen is compatible with her perceptual report of seeing the vertical black and red stripes of the induction screen alternating between a vertical and horizontal orientations.

##### 1.9. Grasping the extremities of a line

In this task, three types of stimuli (black lines, black lines ending with a dot at each extremity or two dots) were displayed on a sheet of paper hanged at a comfortable distance in front of Davida (see Movie S6, online). On each trial, Davida was asked to place her right thumb and index finger on the extremities of the line or on the dots “as if she were grasping it/them”. Her fingers were inked in order to record her responses. In a first task the importance of accuracy was highlighted, and she carefully moved her fingers to the stimuli. In a second task, the importance of speed was stressed: she was asked to stand in front of the sheet of paper with her eyes closed and to open her eyes and move her fingers to the stimuli as fast as possible at the experimenter’s signal. Movie S6, online illustrates Davida’s performance in these experiments. When accuracy was stressed (Figure S3A), she almost always placed her fingers at approximately (a few millimeters from) the place where the extremities of the line would have been if the line (with or without the dots at the extremities) were rotated by 90 degrees (16/18) while this type of error was rare for the dots alone (1 error, 8/9 correct responses). When speed was stressed (Figure S3B), most of her errors consisted in placing her fingers at a few millimeters from the place where the extremities of the line would have been if the line were rotated by 90 degrees (14/18 trials), but she also made a few errors (3/18 trials, the three last errors illustrated in Figure S3B) consisting in placing her fingers at few millimeters from the place where the extremities of the line would have been if the line where rotated by approximately 45 degrees. As these errors were absent when accuracy was stressed (Figure S3A) and Davida never reported seeing stimuli rotated by 45 degrees, we interpret these errors as the result of hesitations between two percepts. Indeed, Davida reports seeing these lines continuously fluctuating between their accurate orientation and the equivalent of their rotation by 90 degrees. She made only two errors when shown two dots (Figure S3B).

**Fig. S3.**
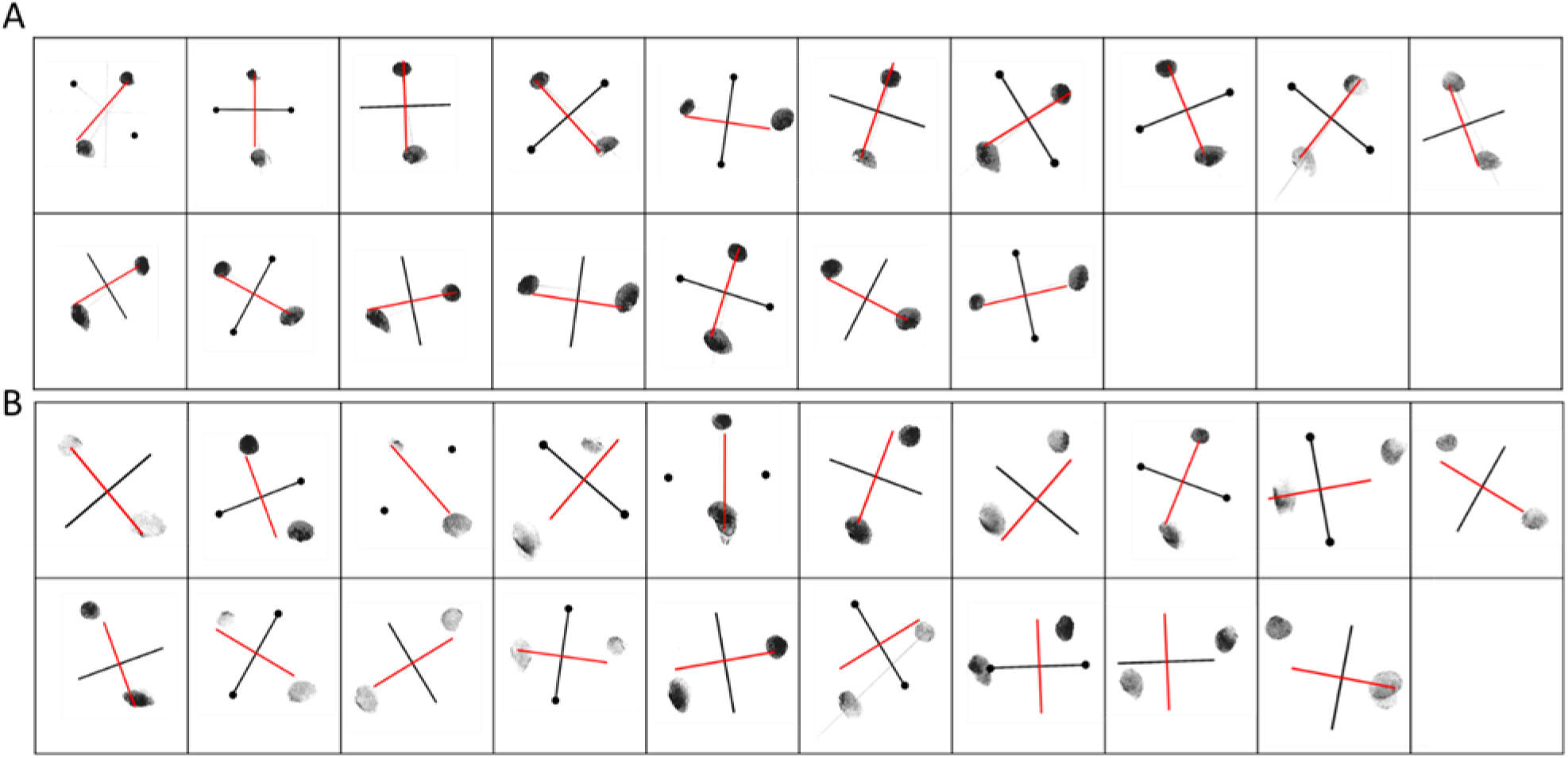
Davida’s fingerprints when she failed to carefully (A) or rapidly (B) move her thumb and index finger to either the extremities of a black line or two black dots. The line in red was added in this figure to illustrate where the line (or a line binding the two dots) would have been if it were rotated by 90 degrees.

##### 1.10. Pointing to the tip of an arrow with the index finger

While standing, Davida saw a large black arrow pointing left, right, up or down displayed on a sheet of paper hung at a comfortable distance in front of her (see Movie S7, online). She was asked to stand with her eyes closed, and, at the experimenter’s signal, to open her eyes and place her index finger as fast as possible on the tip of the arrow. Her index finger was inked in order to record her responses. She almost systematically (25/26 trials) placed her index finger where the tip of the arrow would have been if the arrow were rotated by 90 degrees (34.6%), 180 degrees (7.7%) or 270 degrees (53.8%) clockwise (see Movie S7 online).

##### 1.11. Stimulus–response compatibility task

In this task, participants were shown either a circle or a square on the computer screen and, below it, a black arrow pointing either toward the left or toward the right (see Figure S4). The participants were asked to press a button on the keyboard (the “z” key) as fast as possible with their left index finger when they saw a circle and with their right index finger (the “m” key) when they saw a square, while ignoring the arrow. In each trial, a fixation point was presented at the center of the screen for 500 ms; then the screen was cleared for 500 ms and the stimulus was displayed until a response was recorded. The next trial began after an interval of 1000 ms. The number of different stimuli was 4 (2 shapes x 2 arrow orientations). The experiment included 120 stimuli (30 random repetitions of each stimulus). Congruent trials were those in which the arrow pointed to the correct response key, i.e., an arrow pointing to the key that needed to be pressed for a correct response to a given stimulus. An incongruent trial was that in which the arrow pointed in the opposite direction from the desired key press response.

Analyses were performed to compare participants’ accuracy, latency and efficiency for congruent and incongruent trials. Responses faster than 200 ms (0%) and slower than 1000 ms (3.7%) were excluded from the analyses(Hommel, 1993). Response accuracy and response latency were analyzed separately. Response latency analyses were carried out over correct responses only. In addition, an efficiency score (expressed in ms) was computed for each participant by dividing the mean response latency by the proportion of correct responses in a given condition (thus, the higher the score the poorer the performance). This score allows combining the measures of accuracy and speed into a single measure of processing efficiency; also, it allows between-group comparisons unbiased by potential speed-accuracy tradeoffs(Townsend & Ashby, 1983). Figure S4 displays error rate, mean response latency, and mean efficiency score for Davida and for each control participant for the congruent and incongruent trials.

**Fig. S4.**
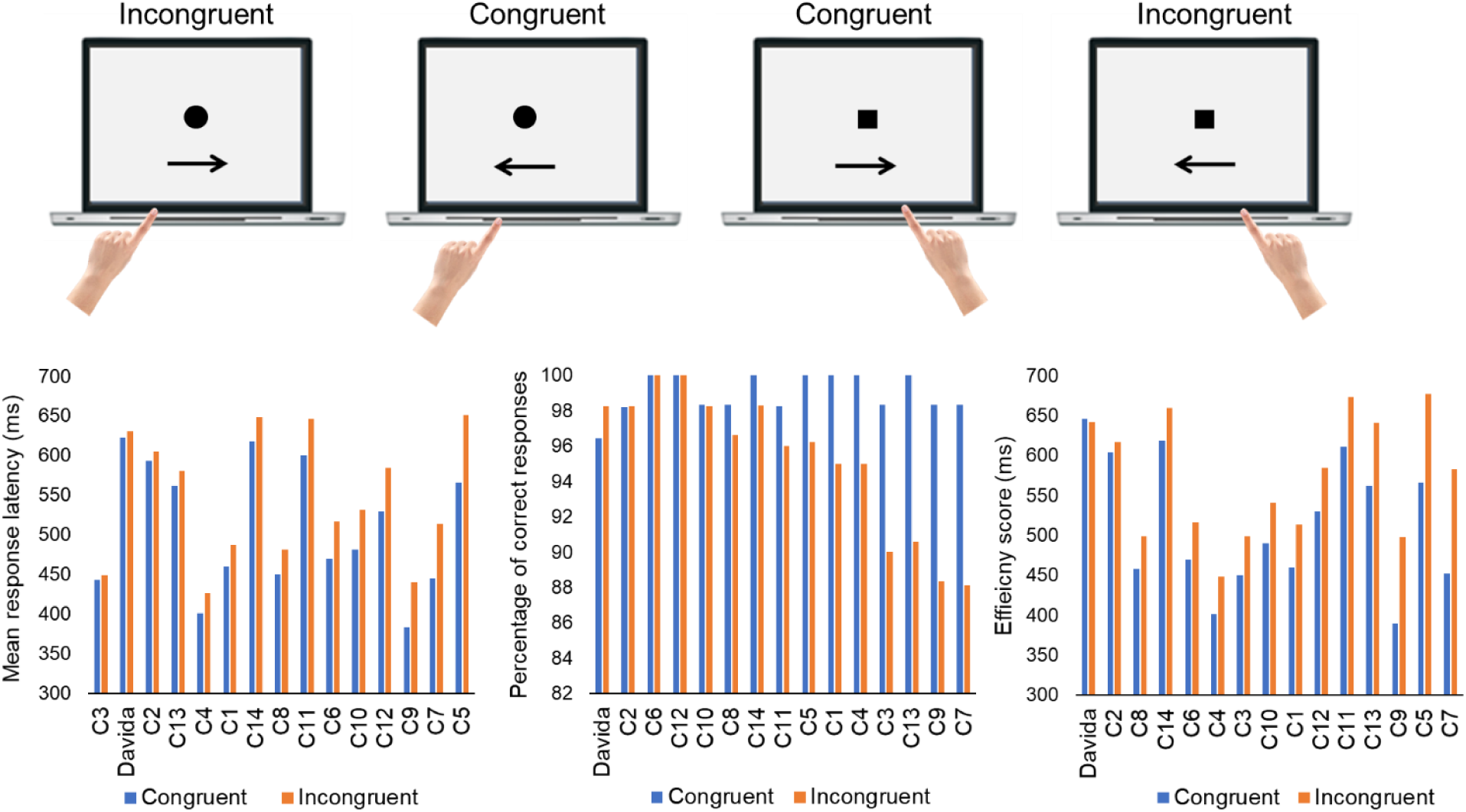
Davida and control participants’ (C1 – C14) accuracy (left), mean response latency (center) and inverse efficiency score (right) in the stimulus-response compatibility task for congruent and incongruent items. Individual data are aligned on the horizontal axis in ascending order as a function of the size of the advantage in speed, accuracy and efficiency for congruent trials.

We first computed paired-sample t-tests to test for an effect of congruency of the stimuli over control participants’ accuracy, RLs and efficiency score. The results of these analyses indicated the typical advantage for congruent over incongruent items in the three variables (accuracy: t(13) = 6.6, p < 0.001; response latency: t(13) = 3.9, p < 0.001; efficiency: t(13) = 7.3, p < 0.001). This typical finding indicates that control participants have an automatic tendency to be influenced by the orientation of the arrow displayed below the stimulus of interest even though the arrow is irrelevant to the actual task (Simon, 1969). To test whether Davida was also implicitly influenced by the orientation of the arrow, we first conducted an independent sample t-test analysis over Davida’s RLs for congruent and incongruent items. This analysis revealed no effect of congruency (t (108) = 0.28, p = 0.78). Finally, we computed unilateral Revised Standardized Difference Test(Crawford & Garthwaite, 2005) over each dependent variable (error rate, response latency, and efficiency score) in order to test whether the discrepancy in Davida’s performance between the congruent and incongruent items was significantly smaller than the discrepancy between both sets in control participants. The results of these analyses revealed that the discrepancy in Davida’s performance between the congruent and incongruent items was almost significantly smaller than that found in control participants when response latency [t (13) = 1.42, p = .09] was considered and significantly smaller when the accuracy [t (13) = .2.99, p < .01] and the efficiency score [t (13) = 1.78, p < .05] were considered as the dependent variable.

##### 1.12. The Ponzo illusion

In this task, participants were presented with the classic Ponzo illusion display, consisting of two internal horizontal lines and two external converging oblique lines (Figure S5). The oblique lines measured 566 pixels and were tilted 10 degrees from the vertical, the upper horizontal line (at the converging end of the display) was 152 pixels (± 2.5 degrees of visual angle) and the lower one (at the diverging end of the display) was of an initially random size composed of between 100 and 190 pixels (± 1.6 and 3.2 degrees of visual angle). The two internal horizontal lines were separated by 600 pixels (10 degrees of visual angle). In each trial, the Ponzo display appeared at the center of the screen and participants had to use the computer’s keyboard to adjust the length of the lower horizontal line (by steps of 2 pixels) to match the length of the upper one and to press the space bar when the two lines appeared identical in length. Participants performed a total of 20 trials. We calculated each participant’s average difference between the lengths of the two horizontal lines. The 14 control participants significantly underestimated the length of the lower horizontal line (t (13) = 11.5, p < 0.001), drawing it on average 20.35 pixels (±0.33 degrees of visual angle) longer than the upper line. This reflects the typical effect of the induced linear perspective in size perception (the Ponzo illusion). Davida, however, did not show the typical visual illusion (Mean difference = +3.5 pixels, SD = 20.8, t (19) = 0.72, p = 0.48) and was significantly better than the controls at matching the length of the two lines (modified t test(Crawford & Howell, 1998): t (13) = −2.45, p = 0.01; 59). See also Figure 2F.

**Fig. S5.**
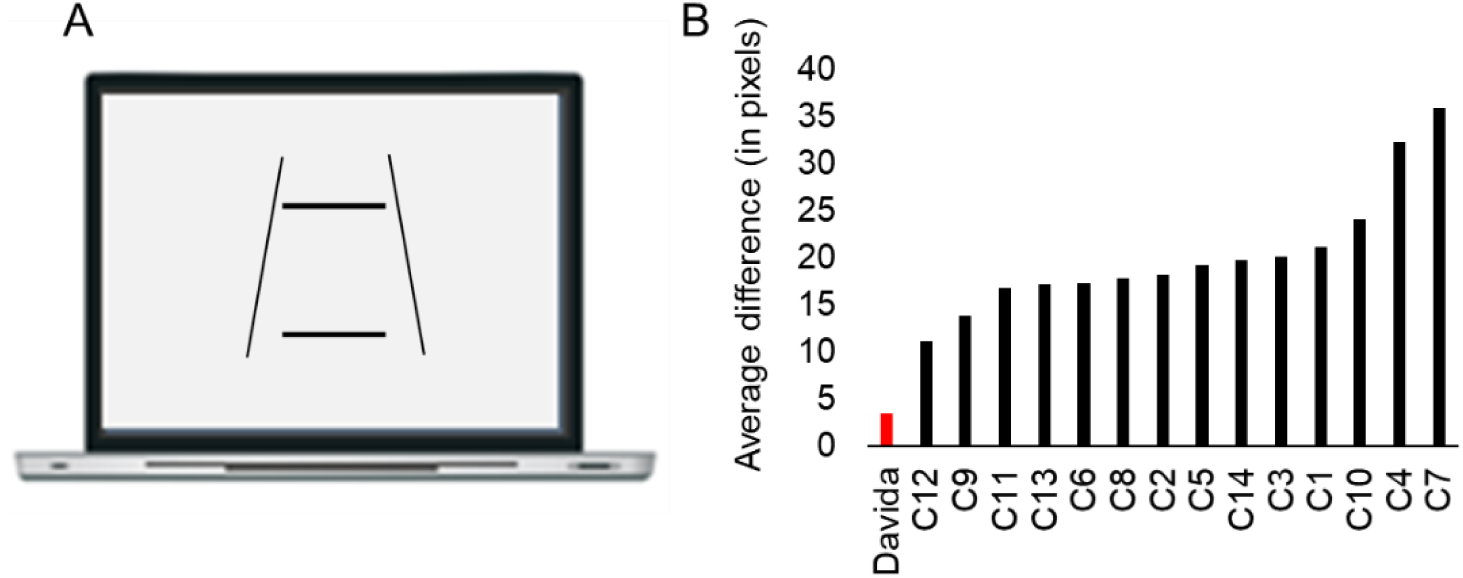
The Ponzo illusion display. B. Davida (in red) and control participants’ (in black) average difference between the length of the two horizontal lines in the Ponzo illusion task (in pixels). Individual data are aligned on the horizontal axis in ascending order as a function of the size of the absolute difference between the length of the two lines.

##### 1.13. The vertical-horizontal illusion

In this task, participants were shown the classic Vertical-Horizontal illusion display, consisting of (1) a black vertical line 10 pixel thick and 604 pixels long centered 500 pixels to the left of the center of the screen and of (2) a black horizontal line 10 pixel thick, between 575 and 620 pixels long at the beginning of each trial and centered at the center of the screen (Figure S6A). In each trial, the display appeared, and participants had to use the computer’s keyboard to adjust the length of the lower horizontal line (by steps of 2 pixels) to match the length of the vertical one and to press the space bar when the two lines appeared identical in length. Control participants performed 20 trials and Davida performed the same experiment twice for a total of 40 trials. We calculated each participant’s average horizontal line’s length. On average, the control participants significantly underestimated the length of the horizontal line (t (13) = 2.89, p = 0.012), drawing it 19.2 pixels (± 0.3 degrees of visual angle) longer than the vertical line. Davida, however, did not show the typical visual illusion (Mean difference = −1.45 pixels; t (39) = 0.13) and was among the two best participants at matching the length of the two lines (see Figure S6).

**Fig. S6.**
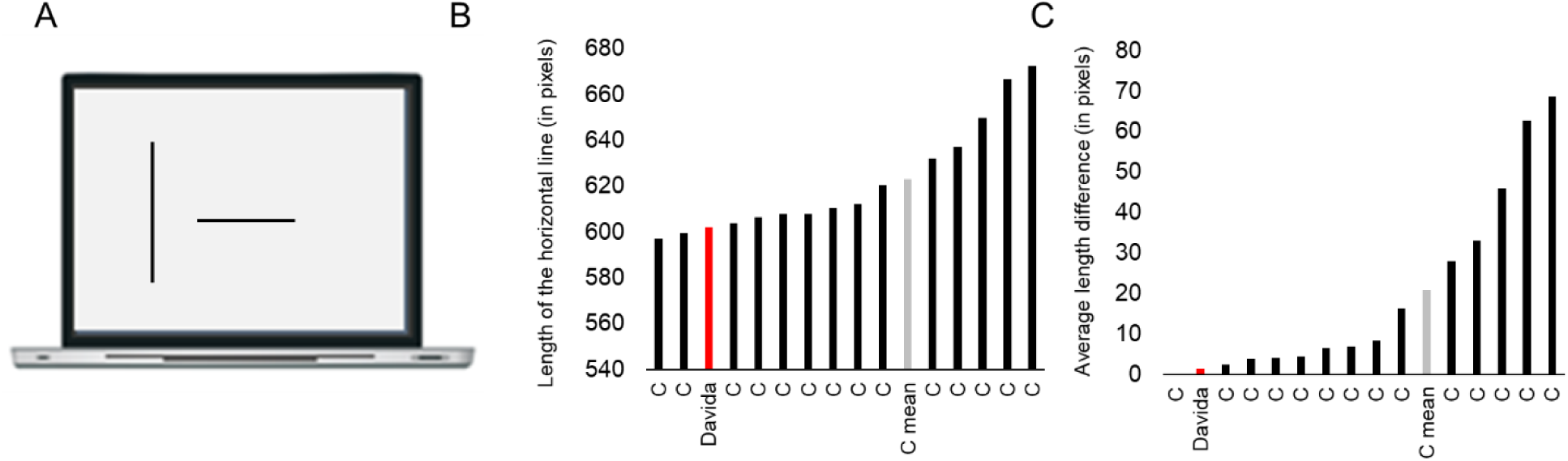
A. The Vertical-Horizontal illusion display. B. Average length of the horizontal line in Davida (in red) the control participants (Cs, in black) and the average of the control participants (C mean, in grey). C. Absolute value of the length difference between the horizontal and vertical line in Davida (in red), the control participants (Cs, in black) and the average of the control participants (C mean, in grey).

##### 1.14. Line drawing naming task with progressive unmasking

Davida was shown 20 line-drawings of objects from the Snodgrass and Vanderwart (60) set displayed on a white background (± 3.3 x 4.6 degrees of visual angle), either in their canonical orientation (10 stimuli) or rotated 180 degrees (10 stimuli). During the experiment, she sat in front of a computer screen located at a distance of about 60 cm, and each trial began with the presentation of a fixation cross for 1000 msec, followed by a stimulus displayed during two frames (32 msec). She was asked to either name the object if she recognized it, or to increase the presentation duration by one frame (+16 msec) by clicking on the space bar. She was invited to repeat this procedure until the presentation duration was long enough for the object to be recognized. There was no time constraint for responding. In line with her report of perceiving 2D stimuli randomly alternating between different orientations through piece-meal gradual transitions, Davida could not recognize any object at the shortest presentation time. Instead, she increased the duration of the stimuli on average 39.05 times (624 msec) before providing a response (which was then always correct). Interestingly, there was no difference between the stimuli displayed in their canonical orientation (606 msec) and those displayed 180 degrees rotated (643 msec).

#### 2. Materials and methods: set of results §2

##### 2.1. Assessment of shape perception and recognition

###### Shape discrimination task

In each trial of the shape discrimination task, participants were shown three shapes of different colors (black, red, green). The reference black shape and one of the probes (red or green, randomly) had the exact same shape (they had edges of 6, 6.1, 1 and 2 degrees of visual angle, see Figure S7), whereas the other probe (the target) had a slightly longer edge. In each trial, any of the four edges of the target could be slightly longer. Participants were asked to report verbally which of the green or red figures had a slightly different shape than the black one at the center. We used a staircase method to measure the threshold magnitude of size difference between the edges required for Davida to correctly discriminate the shapes. The experiment started with a large (+3 degrees of visual angle) difference in edge size, after which the difference decreased every three successive correct responses and increased after any incorrect response (step sizes of 8, 4, 4, 2, 2, 1, 1 db). The session was terminated after 10 reversals. Shape difference sensitivity was defined as the average threshold of the 6 last reversals. Davida had a sensitivity threshold of 0.052 degree of visual angle, that is, she correctly discriminated the shape 80% of the time when one had an edge 0.052 degree of visual angle longer than the comparison one (see, for instance, Figure S7). This performance was comparable to that of the control participants (Mean: 0.13; SD: 0.08; modified t-test(Crawford & Howell, 1998): t (13) = −0.87, p = 0.4; 59).

**Fig. S7.**
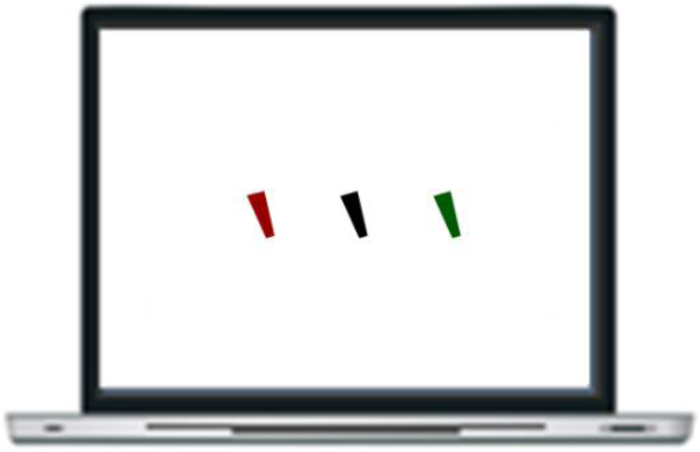
Shape discrimination task. In this example, the red shape’s left edge is slightly longer than that of the two other shapes. The magnitude of this difference (0.052 degree) corresponds to Davida’s sensitivity threshold.

###### Line drawing naming task

Davida was asked to verbally name as accurately as possible 60 line-drawings of objects from the Snodgrass and Vanderwart (60) set displayed on a white background (± 3.3 x 4.6 degree of visual angle) for as long as it took for her to respond. Davida named accurately all the line drawings but reported seeing the line drawings randomly fluctuating among different orientations alternating through piece-meal gradual transitions. We inferred that although her recognition of objects is preserved, it should be somewhat slower than normal. This was confirmed in the experiment reported below.

###### Line drawing naming task with progressive unmasking

Davida was shown 20 line-drawings of objects from the Snodgrass and Vanderwart (60) set displayed on a white background (± 3.3 x 4.6 degrees of visual angle), either in their canonical orientation (10 stimuli) or rotated 180 degrees (10 stimuli). During the experiment, she sat in front of a computer screen located at a distance of about 60 cm, and each trial began with the presentation of a fixation cross for 1000 msec, followed by a stimulus displayed during two frames (32 msec). She was asked to either name the object if she recognized it, or to increase the presentation duration by one frame (+16 msec) by clicking on the space bar. She was invited to repeat this procedure until the presentation duration was long enough for the object to be recognized. There was no time constraint for responding. In line with her report of perceiving 2D stimuli randomly alternating between different orientations through piece-meal gradual transitions, Davida could not recognize any object at the shortest presentation time. Instead, she increased the duration of the stimuli on average 39.05 times (624 msec) before providing a response (which was then always correct). Interestingly, there was no difference between the stimuli displayed in their canonical orientation (606 msec) and those displayed 180 degrees rotated (643 msec).

##### 2.2. Assessment of size perception

###### Line bisection task

Davida was shown 69 lines of different lengths (± 5.5, 9.1 or 12.8 degree of visual angle) and orientations (0, 45 or 90 degrees) displayed one by one randomly placed at various locations on the screen, and was asked to use the computer mouse cursor to click on the middle of each line. Davida reported seeing all lines in two alternating orientations 90 degrees off one another and clicked at the point where the two lines crossed. Davida clicked on average at an absolute distance of 9 pixels (SD: 7.34) off from the center of the lines (0.15 degree of visual angle) with a tendency to click very slightly above (1.6 pixels ± 8 SD) and to the right (4.2 pixels ± 7 SD) of the center. This performance was not different from that of the controls in both absolute distance (mean distance: 8.35 pixels; SD: 2.81; range: 0.014 – 0.262; modified t-test(Crawford & Howell, 1998): t (13) = 0.23, p = 0.82) and bias (Mean: 0.3 ± 3.5 SD pixels below and 1.5 ± 2.8 SD pixels to the left of the center; both modified t-tests(Crawford & Howell, 1998): ts (13) < 1.97, p > 0.05).

###### Size comparison task

In each trial of this experiment, two circles were displayed simultaneously on the computer screen for as long as needed, and Davida had to report verbally which one (the green or the red) was smaller. In each trial, one circle (the green or the red, randomly) had a diameter of 1 degree of visual angle and the other one was slightly larger. We used a staircase method to measure the threshold magnitude of size difference required for Davida to identify the smaller circle correctly. The experiment began with a large size difference of +3 degrees of visual angle, after which, the size difference decreased following three successive correct responses or increased after a single incorrect response (step sizes of 8, 4, 4, 2, 2, 1, 1 db). The session was terminated after 10 reversals. Davida’s sensitivity to size differences was defined as the average threshold of the 6 last reversals. Davida had a sensitivity threshold of 0.015 degree of visual angle, that is, she correctly indicated the smaller circle 80% of the time when the foil had diameter of 1.015 degree of visual angle. This performance was equivalent to the performance of the control participants (mean: 0.021; SD: 0.009; modified t-test(Crawford & Howell, 1998): t (13) = −0.61, p = 0.55).

##### 2.3. Assessment of distance perception

In each trial of this experiment Davida was shown three circles, one green, one red, a third black, aligned in the horizontal plane (each circle had a diameter of 1 degree of visual angle) and had to report verbally which of the two colored circles (green or red) was closer to the black circle (the reference), which was placed between the two targets. In each trial one of the colored probes was placed at a distance of 5 degree of visual angle and the other one slightly farther away. We used a staircase method to measure the threshold magnitude of distance difference required for Davida to correctly identify the circle that was closer to the reference circle. The experiment began with a large size difference of +3 degrees of visual angle, after which the size difference decreased after every three successive correct responses and increased after even one incorrect response (step sizes of 8, 4, 4, 2, 2, 1, 1 db). The session was terminated after 10 reversals. Davida’s sensitivity to distance difference was defined as the average threshold of the 6 last reversals. She had a sensitivity threshold of 0.17 degree of visual angle, that is, she correctly indicated which circle was the closer 80% of the time when the foil was at a distance of 5.17 degree of visual angle. This performance was similar to that of control participants (mean: 0.14; SD: 0.067; modified t-test(Crawford & Howell, 1998): t (13) = 0.36, p = 0.73).

##### 2.4. Assessment of location perception

On a first pointing task, Davida saw series of small black circles, each having a diameter of 0.69 degrees of visual angle (6mm, 40 pixels) appearing one at a time for 1 second at one of 15 possible locations on the screen. The 15 locations constituted a grid made of 5 columns and 3 rows covering the entire screen. After a circle disappeared, Davida heard a brief tone and was asked to use the computer mouse to click on the screen where the circle had appeared. There were three trials per location. She clicked on average at a distance of 20 pixels from the center of the circle (0.35 degree of visual angle, 3 mm). This performance was similar to that of control participants. (mean distance: 20.8 pixels; SD: 7; modified t-test: t (13) = −0.12, p = 0.91).

In a second pointing task, the procedure was the same, except that Davida was asked to fixate on a dot at the center of the screen until the to-be-localized circle appeared for 48 milliseconds (3 frames; 60Hz). She located the circle within an average distance of 54 pixels from its center (0.94 degrees of visual angle, 8.1 mm).

##### 2.5. Assessment of line tilt perception

Davida was presented with a series of three lines (each 10 degrees of visual angle long, 0.25 degrees of visual angle thick, centered 5 degrees of visual angle apart on the horizontal plane). In each trial, the central black reference line and one probe line (green or red) were vertically oriented and the third probe was slightly rotated away from the vertical. Davida’s task was to decide which of the two probe lines (the red or the green) was perfectly aligned with the central reference line. We used a staircase method to measure the threshold magnitude of orientation difference required for Davida to correctly identify the misoriented line. The experiment started with a large size difference of +10 degrees, after which the size difference decreased after every three successive correct responses and increased after any incorrect response (step sizes of 8, 4, 4, 2, 2, 1, 1 db). The session was terminated after 10 reversals. Davida’s orientation difference sensitivity was defined as the average threshold of the 6 last reversals. She had a sensitivity threshold of 0.88 degrees, that is, she correctly indicated which line was differently oriented 80% of the time when the foil was tilted 0.88 degrees from the vertical. This performance was similar to that of the control participants (mean: 0.58; SD: 0.27; modified t-test(Crawford & Howell, 1998): t (13) = 1.06, p = 0.3).

##### 2.6. Assessment of movement perception

###### Point-light walker

Davida saw one of two possible point-light walker animations (one facing to the left and one facing to the right) made of 14 dots placed on the main joints, and she was asked to judge in which direction the point-light walkers were facing. She performed this task flawlessly and easily (10/10).

###### Motion coherence

Davida was presented with a series of circular random dot kinematograms (RDKs) composed of black dots displayed on a uniform white background. The field size of the RDKs was 10° in diameter, dot size was 30 pixels, dot density was 100 dots/frame and dot velocity was 0.6°/s. A proportion of dots moved coherently toward the top, bottom, left or right of the screen and the remaining noise dots moved in random directions. On each trial, the RDK was shown for 1 s, and Davida was asked to tell the direction of coherent motion. We used a staircase method to measure the threshold proportion of signal dots required to correctly discriminate the direction of coherent motion. The session began with an RDK composed of 90% signal dots. Then, the signal to noise proportion was decreased after three successive correct responses and increased after one incorrect response (step sizes of 8, 4, 4, 2, 2, 1, 1 db). The session was terminated after 10 reversals. Coherence sensitivity was defined as the coherence threshold of the 6 last reversals. Davida had a coherence threshold of 15%. This performance was equivalent to that of the control participants (mean: 22.6; SD: 7.6; modified t-test(Crawford & Howell, 1998): t (13) = −0.96, p = 0.36).

##### 2.7. Assessment of auditory processing of orientation

Davida was exposed to one of three different pure tones (24, 250 or 337 Hz) lasting 1.9 seconds to either the right or left ear at a comfortable intensity and had to report whether the sound had been presented to her right or left ear. The task contained 30 trials (2 ears x 3 sounds x 6 repetitions). She scored 30/30.

##### 2.8. Assessment of tactile processing of orientation

###### Laterality judgment task

Davida was asked to position her two hands on a table in front of her, close her eyes, and report which hand had been gently touched by the experimenter with a pen. She was asked to respond by lifting the touched hand. She scored 20/20.

###### Stereognosis

On the first task Davida was asked to close her eyes, explore a 3D wooden arrow positioned in front of her with her two hands and decide in which direction the arrow was pointing (left, right, up or down). She was asked to respond verbally. She scored 20/20.

On the second task, Davida was asked to close her eyes, explore a 3D wooden letter positioned in front of her with her two hands and decide whether the letter was oriented so that it constituted a “b”, “d”, “p” or a “q”. She was asked to respond verbally. She scored 20/20.

###### Tactile integration

Davida was blindfolded, asked to place her left hand comfortably on a table in front of her and decide which of four possible letters (b, d, p, q) was traced on the dorsal surface of her hand (see Movie S8 online). She scored 20/20.

###### Transcoding

On the first task, Davida was first to place her left hand comfortably on a table in front of her, to close her eyes while one of four possible letters (b, d, p, q) was traced on the back of her hand by the experimenter. Then, she was asked to open her eyes and draw the letter on a sheet of paper placed in front of her. There were 20 trials and she performed the task perfectly (20/20) and without hesitation.

On the second task, Davida was asked to place her left hand comfortably on a table in front of her and to close her eyes while an arrow was traced on the back of her hand by the experimenter in one of four possible orientations (left, right, toward and away from her body). Then, she was asked to open her eyes and draw the arrow on a sheet of paper placed in front of her. There were 20 trials and she performed the task perfectly (20/20) without hesitation.

##### 2.9. Writing from memory

Davida was asked to write on dictation a series of 15 letters and 15 words. She made only one error, consisting of reversing the two last letters in the word “table” (“tabel”). Interestingly, as one can see in Figure S8, she used a personal font in which orientation-sensitive letters are attributed different shapes. She reported having developed this strategy to be able to read her own writing.

**Fig. S8.**
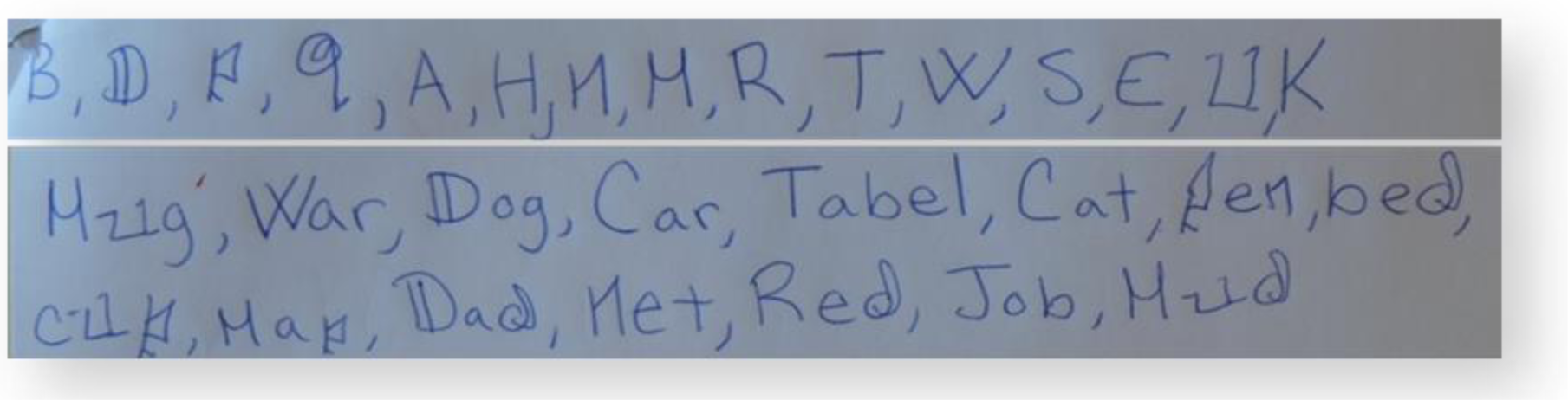
Davida writes letters (‘b’, ‘d’, ‘p’, ‘q’, ‘a’, ‘h’, ‘n’, ‘m’, ‘r’, ‘t’, ‘w’, ‘s’, ‘e’, ‘u’, ‘k’) and short words (‘mug’, ‘war’, ‘dog’, ‘car’, ‘table’, ‘cat’, ‘pen’, ‘bed’, ‘cup’, ‘map’, ‘dad’, ‘net’, ‘red’, ‘job’, ‘mud’) on dictation. She uses different shapes to discriminate orientation sensitive letters (e.g., ‘b’, ‘d’, ‘p’, ‘q’).

#### 3. Materials and methods: set of results §3

##### 3.1. Arrow orientation naming in various locations mono and binocularly

Davida was shown black arrows on white background and asked to decide whether the arrow pointed “up”, “down”, “left” or “right”. Stimuli were ± 1.5 x 3 degrees of visual angle and displayed at one of five different locations: at the center, upper left corner, upper right corner, lower left corner or lower right corner of the computer screen. The locations near the corner of the screen were at horizontal and vertical distance of 10 degrees of visual angle from the center position. The total number of different stimuli was of 20 arrows (5 positions × 4 orientations). During the experiment Davida was asked to fixate a cross located at the center of the screen, stimuli appeared 100 milliseconds and Davida was asked to respond verbally (“left”, “right”, “up”, “down”). The next trial was launched by the experimenter. She performed 5 blocks of 20 trials with both eyes, then with only the left eye, then with only the right eye. As shown on Table S3, Davida made only a few correct responses and her response profile was not affected by the location of the stimulus or the eye(s) used to solve the task.

**Table S3.**
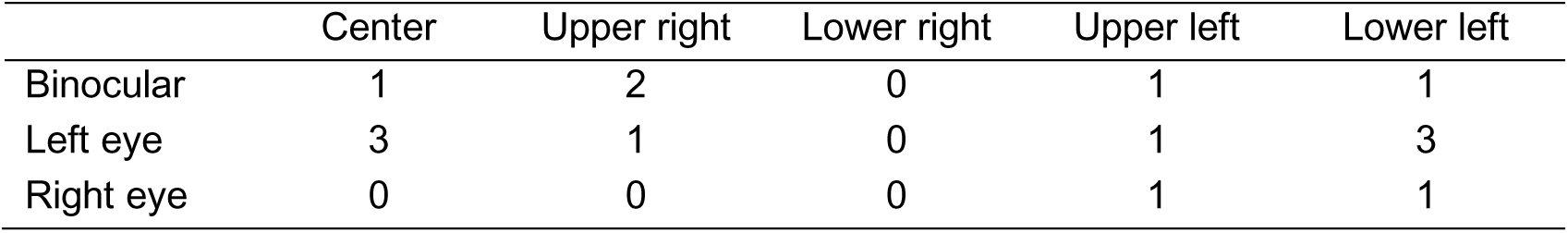
Davida’s number of correct responses (/20) in the arrow orientation naming task according to the location of the stimulus and eye(s) used.

##### 3.2. Localizing the tip of an arrow by pointing with short exposure duration, masking and various locations

In this task, a small (± 4 x 8 degrees of visual angle) black arrow was displayed at one of five different locations: at the center of the screen or at ± 4 degrees of visual angle above, below, on the left or on the right of the center. The arrow appearing at the center could be pointing left, right, up or down. The arrow appearing above or below the center could be pointing up or down. The arrow appearing on the left or the right of the center could be pointing left or right (see Figure S9). On each trial, a fixation cross appeared for 1 second at the center of the screen followed by the arrow for 80 msec and a visual mask (Figure S9A). Davida was asked to fixate the fixation cross and, then, to use the computer mouse to move a small round cursor and click as precisely as possible on the place where the tip of the arrow had appeared. The next trial was launched by the experimenter. There was a total of 40 trials in which the arrow was centered on the fixation cross and 20 trials for each other position (10 trial by orientation). As shown in Figure S9B, Davida almost systematically mislocated the position of the tip to approximately the place it would have been if the arrow were rotated by 90 degrees, 180 degrees, or 270 degrees with respect to its own center.

**Fig. S9.**
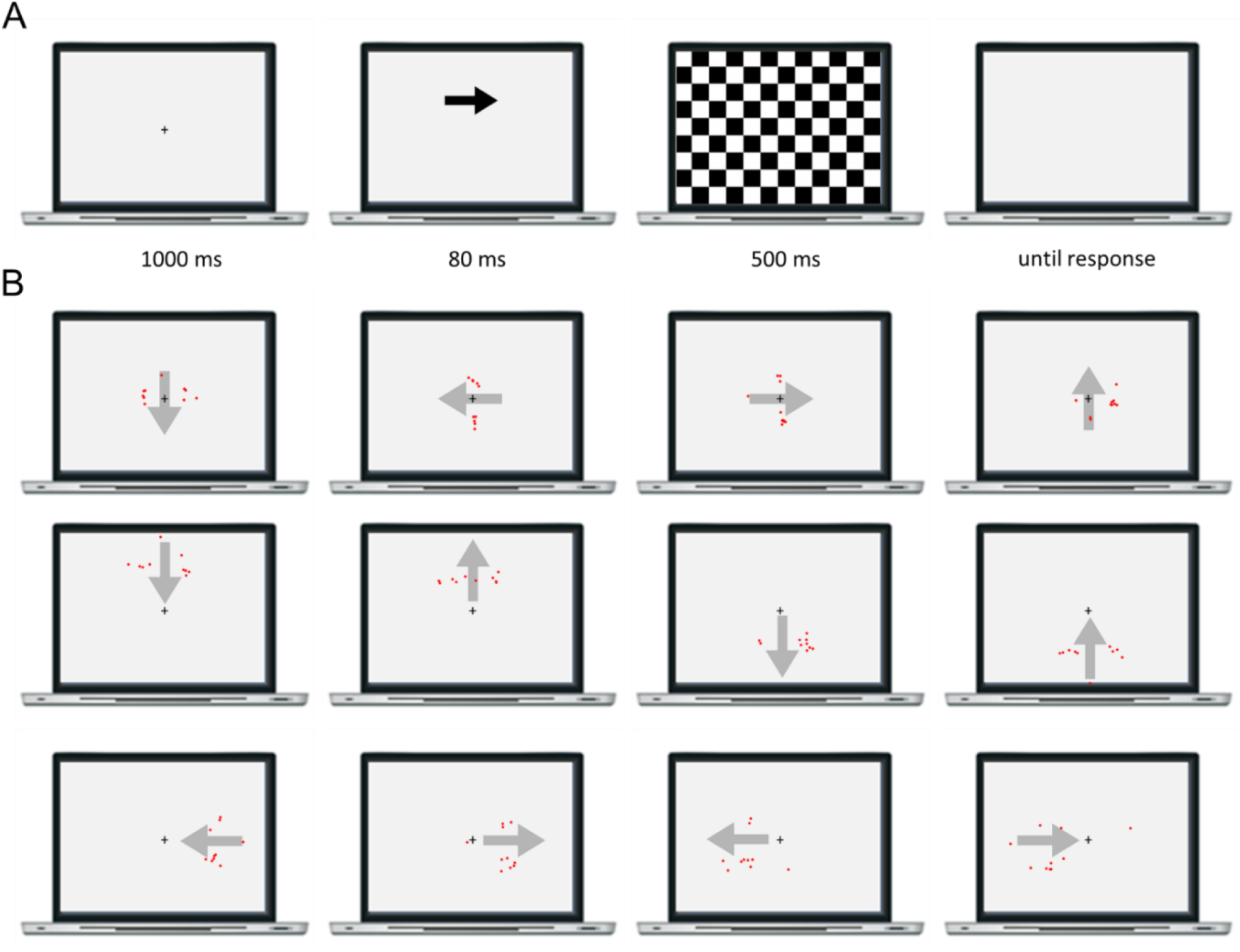
A. Illustration of the procedure. B. Each red dot represents the coordinates of one of Davida’s attempt at localizing the tip of an arrow. Arrows appear in grey here for transparency, they were displayed in black during the experiment.

##### 3.3. Copying two shapes presented simultaneously

Davida was shown 6 different series of 2 unconnected shapes (an arrow and a rectangle) and asked to copy them as accurately as possible on a separate sheet of paper while the stimulus remained in view. These shapes were copied 4 or 5 times. As shown in Figure S10, the two shapes were copied as if they were rotated or inverted independently from each other.

**Fig. S10.**
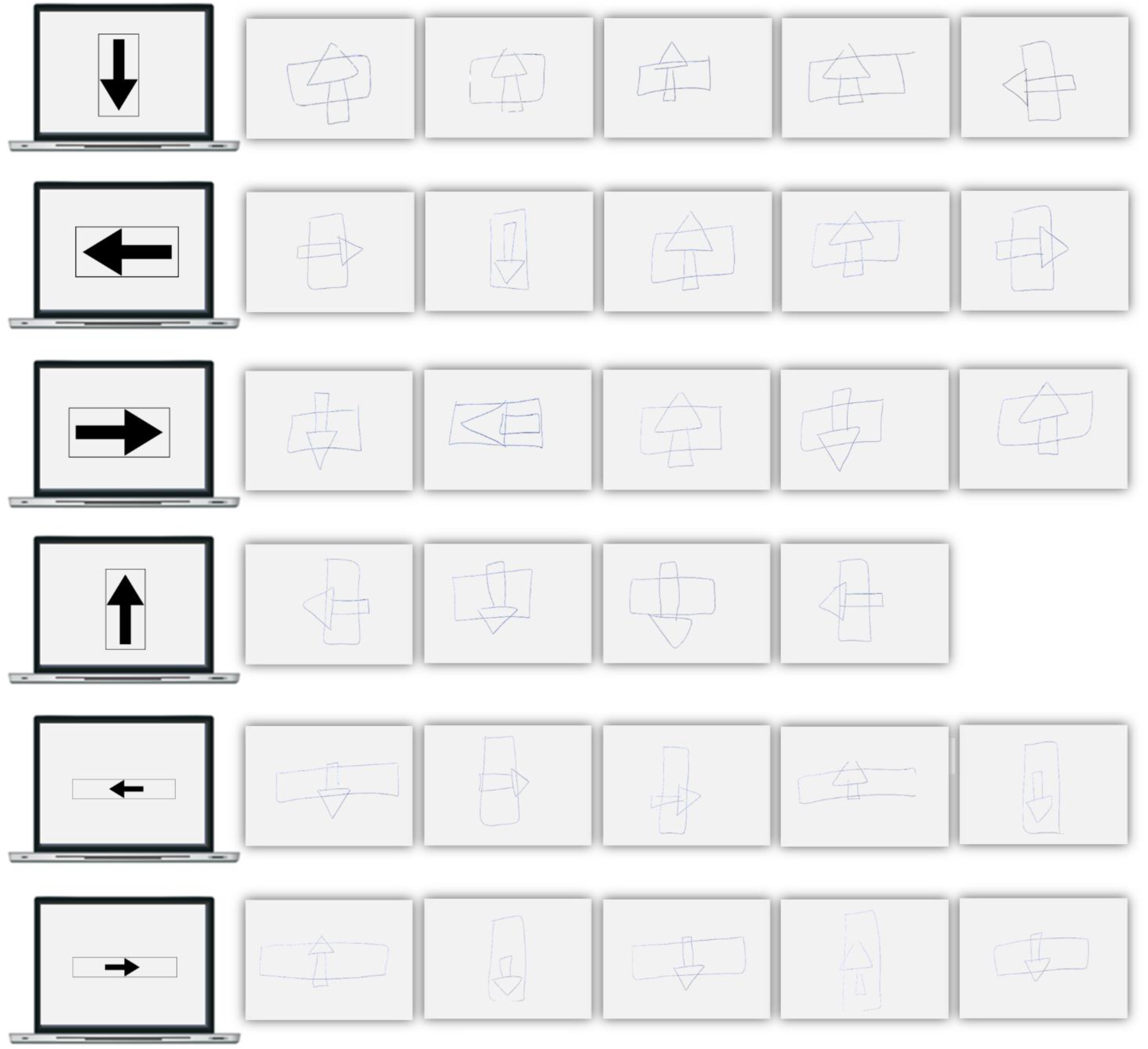
Davida’s multiple attempts at copying of 2 unconnected shapes (displayed on the computer screen).

##### 3.4. Copying two briefly displayed arrows

In this task, two small black arrows were displayed simultaneously (± 4 x 8 degrees of visual angle each) for 32 ms at ± 8 degrees of visual angle left and right of center of the screen. Davida was asked to look carefully and, then, to draw what she has seen on a separate sheet of paper. Each arrow was displayed 5 times in 4 possible orientations (left, right, up, down). Davida copied only 3/40 arrows accurately. All her errors consisted in rotating the arrows by 90, 180 or 270 degrees. Of particular interest in this task was whether Davida made the same or different orientation errors for the two arrows. The two arrows were drawn as if their perception resulted from the same orientation error in 6/20 trials and from different errors in 14/20 trials.

##### 3.5. Judging whether briefly displayed arrows are in the “same or different” orientation

In this task, two small black arrows (± 4 x 8 degrees of visual angle each) were displayed for 32 ms at ± 8 degrees of visual angle left and right of center of the screen. Davida was asked to look carefully at them and to decide whether the two arrows had the same direction or not. There were 80 trials. Among these, there was the same proportion of trials in which the arrows differed by 0, 90, 180 and 270 degrees. Davida scored 28/80. She failed to recognize that the two arrows were in the same orientation in 80% of the trials (16/20) and failed to recognize that the two arrows were in a different orientation in 60% of the trials (36/60).

##### 3.6. Counting tasks

In this series of task, Davida saw one, two or three black dots displayed alone, together with a large black circle, or with a large black arrow (e.g., Figure S11b-d). Thus, there were 9 different possible configurations (3 numbers of dots x 3 conditions). The dots had a diameter of 12 pixels and, in the default configuration, could be displayed at one of three positions: 140 pixels on the left of the center of the screen, 115 pixels above the center of the screen or 115 pixels below the center of the screen. The large black circle had a diameter of 188 pixels and, in the default configuration, was centered at the center of the screen. The arrow a length of 380 pixels, a maximal width of 220 pixels and, in the default configuration, was centered 70 pixels on the right of the center of the screen. During the experiment, Davida saw the default configuration and the equivalents of its rotation by 90, 180 and 270 degrees. Thus, there were 36 different stimuli (3 numbers of dots x 3 conditions x 4 orientations). In each trial, Davida had to count and then report verbally the number of small dots that she had seen. In three experiments, the stimuli were displayed for 500 msec, 1000 msec or for an unlimited time, respectively. Each experiment comprised 72 trials (each stimulus was seen twice). We counted the number of correct responses by condition. As can be seen in Figure S11a, in the three experiments Davida was able to correctly report the number of dots when the dots were displayed alone (Figure S11b) and when they were shown the large black circle (Figure S11c), but not when they were presented with the black arrow (Figure S11d). All her errors consisted in underestimating the number of dots that had been displayed in this condition.

Three additional counting experiments were performed to explore the specificity of this effect. In one experiment, Davida was presented for 1 second with a large black arrow surrounded by one, two or three black dots positioned either exactly as in the previous experiments (Figure S11d) or slightly above and below where the tip of the arrow would be if it were inverted or rotated by 90 or 270 degrees clockwise (Figure S11e). In a second experiment, Davida was presented for 1 second with a large black arrow surrounded by one, two or three black dots positioned either exactly as in the previous experiments (Figure S11d) or placed where the tip of the arrow would be if rotated by 45, 135 or 315 degrees (see Figure S11f). In a third experiment, Davida saw one, two or three black dots positioned either exactly as in the previous experiments displayed with either the same large black arrow (Figure S11d) or a “transparent” arrow of the same size and shape defined only by its contour (Figure S11g). In all experiments, Davida performed 96 trials (3 numbers of dots x 2 conditions x 4 orientations x 4 trials). We counted the number of correct responses by condition. As can be seen in Figure S11, in the three experiments Davida was unable to correctly report the number of dots when the dots were displayed with the black arrow in locations that the arrow would cover if it were seen as rotated or inverted but performed the task significantly better in the other conditions.

**Fig. S11.**
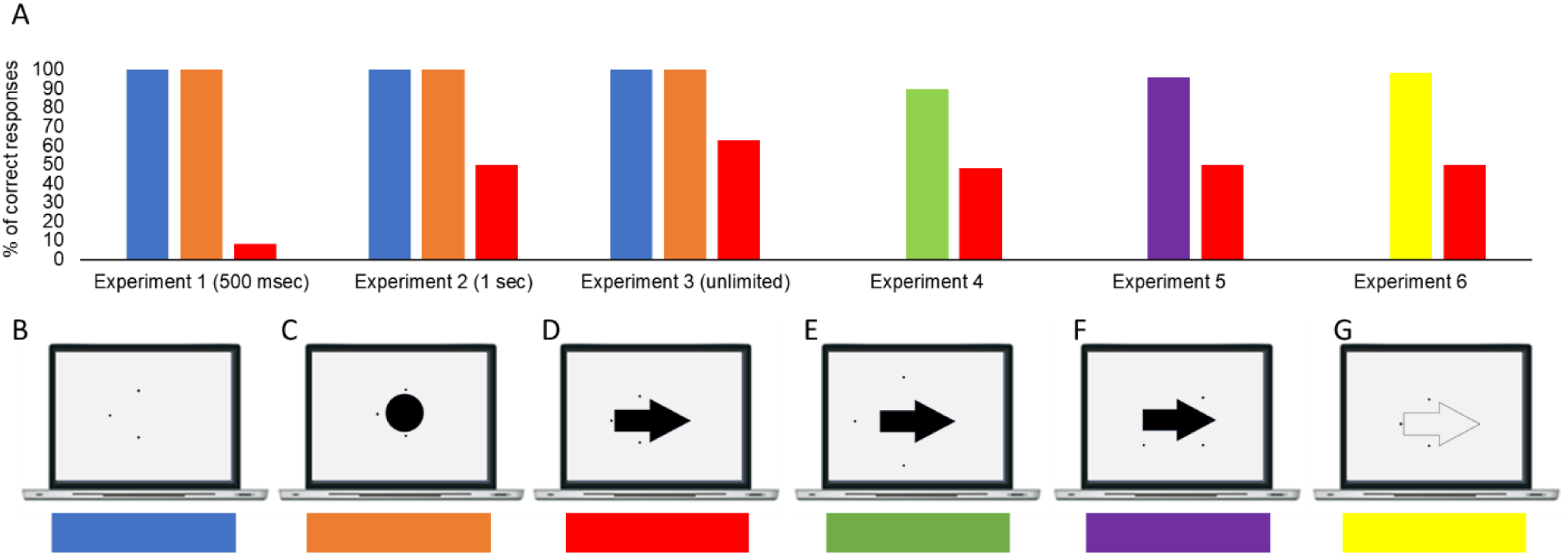
Davida’s percentage of correct responses in the six counting experiments for the different conditions (related by a color code).

##### 3.7. Object naming task below an arrow

This experiment comprised two sessions. In a first session, Davida was asked to name verbally as fast as possible 80 line-drawings of objects from the Snodgrass and Vanderwart (60) set displayed on white background. The line-drawings had a size of 200 x 280 pixels (± 3.3 x 4.6 degrees of visual angle). In each trial, a fixation point was presented at the center of the screen for 200 ms; then the screen was cleared for 500 ms and the stimulus was displayed until a voice key was triggered. The next trial began after an interval of 2000 ms. Malfunctioning of the voice key and Davida’s responses were registered on-line by the experimenter. In a second session, Davida performed a similar task with two differences: (1) she was presented only with the objects for which a valid response time had been collected during the first session (N=74; 4 errors, 2 voice key malfunctioning) and (2) these objects were separated in two sets (N = 37) matched in average naming latency collected during the first session (t (72) < 1) and were displayed either ± 200 pixels below a large black arrow (700 pixels long, maximal width of 370 pixels, see Figure S12a) or ± 200 pixels below three large black squares of same length and maximal width (see Figure S12b).

To explore a possible interference from the arrow on naming the objects displayed below it, we carried out a by-item analysis of variance over the response latencies of Davida for each item with item as the random factor, Session (session 1 vs session 2) as a within-item factor and Set of item (squares vs. arrow) as between-item factor. This analysis was performed on all but one item for which Davida made an error during the second session (in the squares condition). The results are displayed in Figure S12. The analysis disclosed no significant effect of the set of items [F (1, 71) = 1.42, p = 0.24] but a significant effect of the session [F (1, 71) = 16.7, p < 0.001] and a significant Set of items x Session interaction [F (1, 71) = 41.08, p < 0.001]. Independent samples t-tests performed to explore the interaction indicated a slight advantage of the second session for the set of items named in the squares condition (−104 msec; t (35) = 1.18, p = 0.24) but a large and significant disadvantage of the second session for the items in the arrow condition (+474 msec; t (36) = −20.72, p < 0.001).

**Fig. S12.**
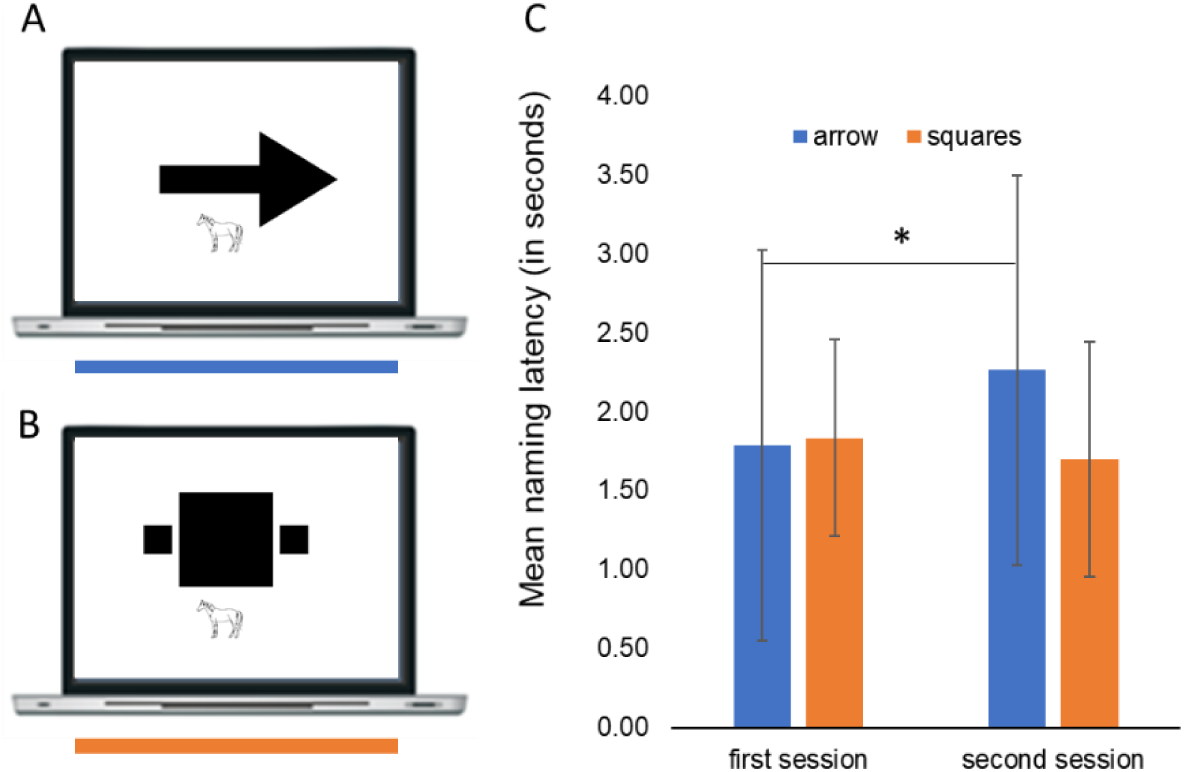
(A and B) Example of the arrow (A) and squares (B) conditions in which the objects were named in the second session. (C) Davida’s mean response latency and standard deviation for the two sets of items after the first and second session. * indicates a statistically significant difference at *p* < 0.001.

#### Materials and methods: set of results §4

##### 4.1. Copying words with connected or unconnected letters

Davida was shown 6 words composed of connected letters (Lucida Handwriting font, size 72) and 6 words composed of unconnected letters (Calibri font, size 72) and asked to copy them as accurately as possible on a separate sheet of paper while the stimulus remained in view. As can be seen in Figure S13, she misrepresented the orientation of the letters when the letters were unconnected but also of the whole word when the letters were connected.

**Fig. S13.**
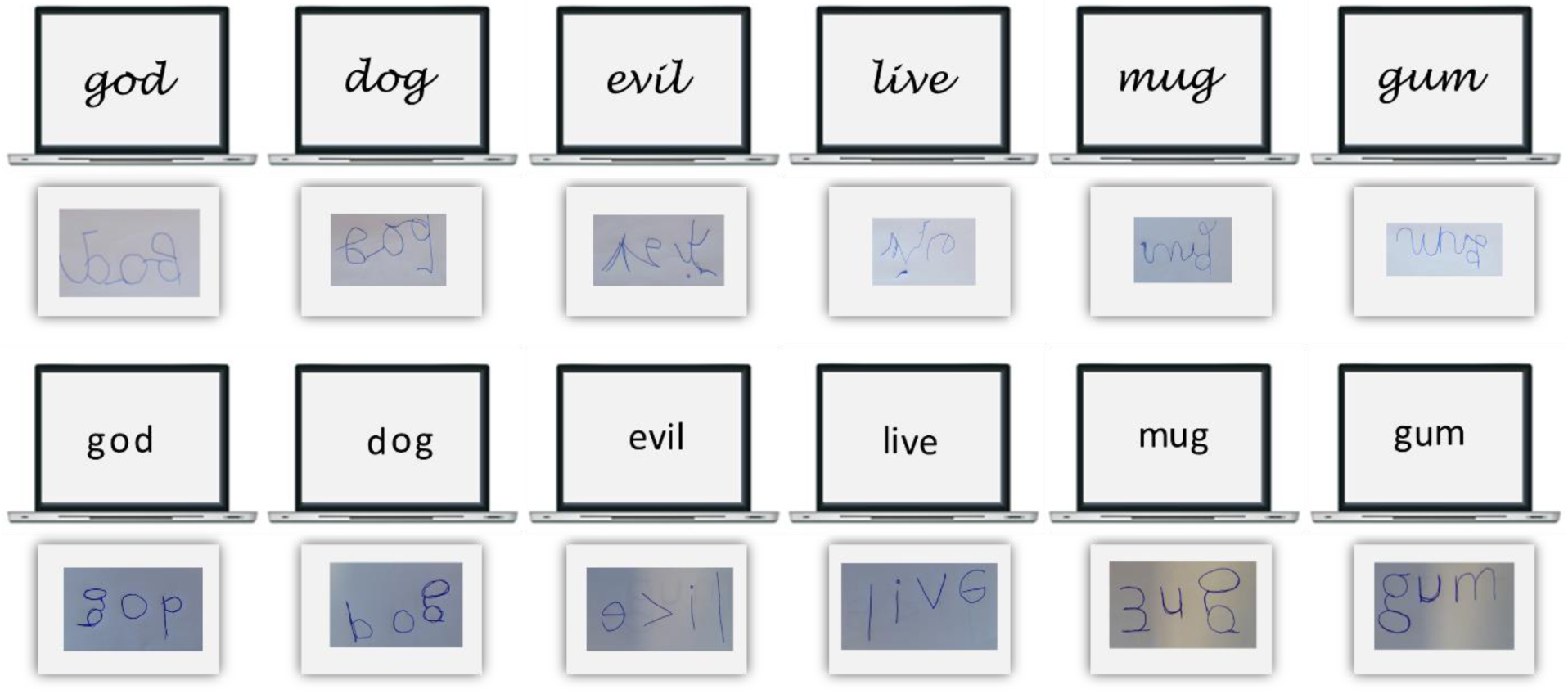
Davida’s copy of words with attached (top) and non-attached (bottom) letters.

##### 4.2. Clicking on a dot that is touching or not touching an arrow

Davida was shown a black or red dot positioned to the right or left of center of the screen alone (Figure S14, A), near to the tip of an arrow (Figure S14, B), touching the end of an arrowhead (Figure S14, C, D) or included within the arrow (Figure S14, E). In each trial, Davida was asked to use the computer mouse to move a small round cursor and click as precisely as possible on the dot. Each trial appeared 1 second after the response to the previous trial. There was no time constraint, as the emphasis was on accuracy. There were 20 trials per each condition. Davida clicked on the dot accurately in 100% of the trials in the first (A and B) and last (D and E) two conditions, but only 1/20 trials when the dot was black and connected to the tip of the arrow (C). In the latter condition, she clicked approximatively where the dot would have been if the arrow to which it was attached was rotated by 90, 180 or 270 degrees.

**Fig. S14.**
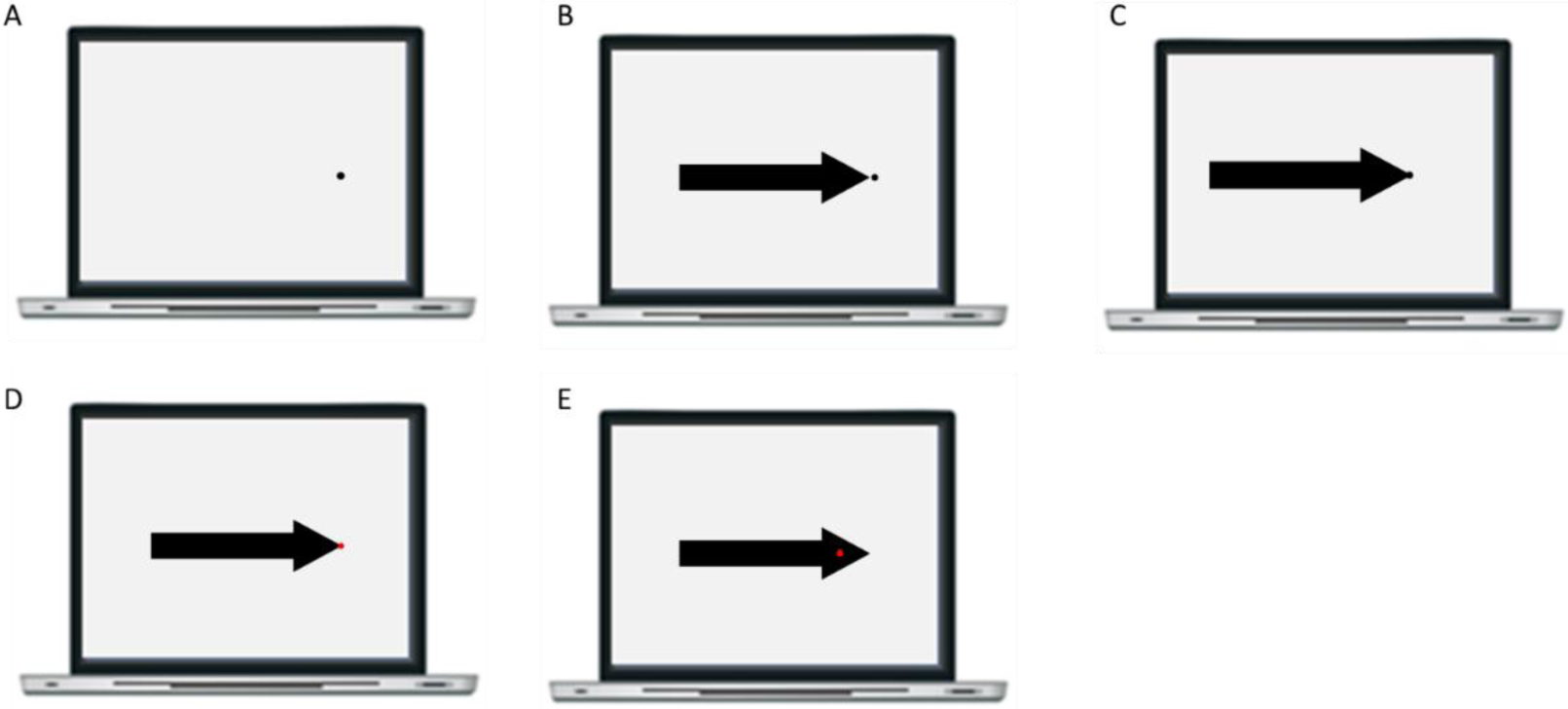
Illustration of the stimuli used in Experiment 4.2.

##### 4.3. Naming the color behind the tip of an arrow

In this task, Davida was presented with each of the stimuli displayed in Figure S15 20 times and had to name the color behind the tip of the arrow. Davida correctly identified the color behind the tip of the arrow in 2/80 trials and made 78/80 errors consisting of responding as if the arrow was rotated by 90 (42.5%), 180 (20%) or 270 (35%) degrees.

**Fig. S15.**
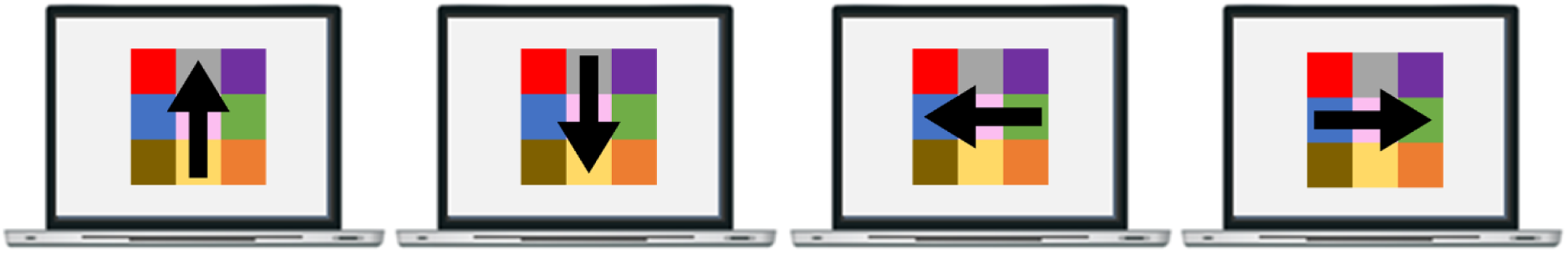
Stimuli used Experiment 4.3

##### 4.4. Pointing to the tip of an arrow

A long thin arrow (±6 x 0.5 degrees of visual angle) pointing left, right, up or down was displayed at the center of the computer screen for an unlimited duration (Figure S16). In half of the trials (N = 40), the arrow was fully depicted (Figure S16A). In the other half, the center of the arrow was hidden behind a mask (5 pixels) that had the same color as the background (Figures S16B). In each trial, Davida was asked to use the computer mouse to move a small round cursor and click as precisely as possible on the tip of the arrow. In the condition in which the arrow was fully depicted, Davida located the position of the tip of the arrow accurately (i.e., she clicked at less than 50 pixels from the accurate position of the tip) in 2 trials (5%) and mislocated the position of the tip to approximately (i.e., less than 50 pixels) where it would have been if the arrow had been rotated by 90 degrees (30%), 180 degrees (7.5%) or 270 degrees (57.5%). In the other condition, Davida located the position of the tip of the arrow accurately (i.e., less than 50 pixels) in 1 trial (2.5%) and mislocated the position of the tip to approximately (i.e., less than 50 pixels) where it would have been if only the part of the arrow connected to the tip had been rotated by 90 degrees (32.5%), 180 degrees (10%) or 270 degrees (55%). The Movie S9, online, is a recording of Davida performing this task and illustrate her response profile.

**Fig. S16.**
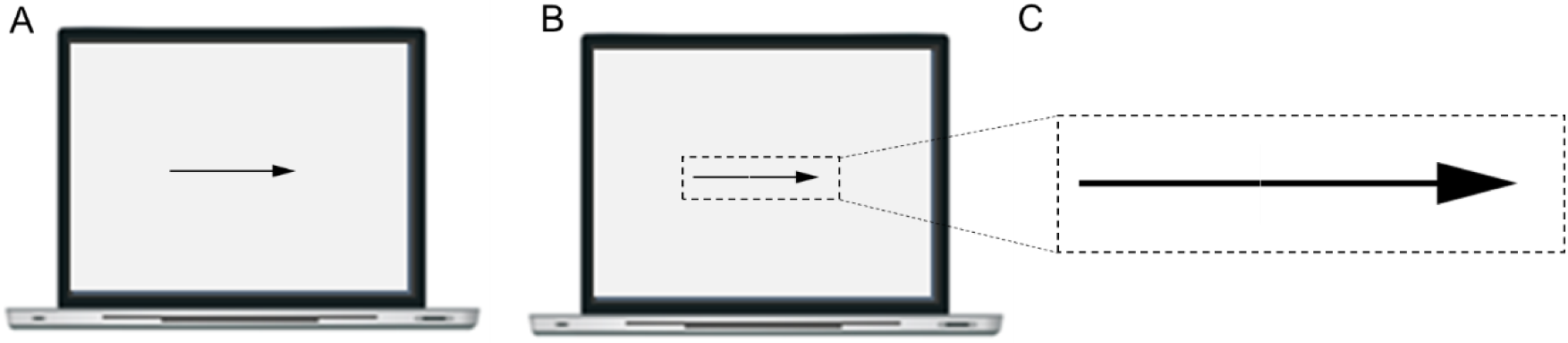
A, B. Stimuli used in Experiment 4.4. The dashed lines illustrate the area of the screen selected for the zoom (C) on the arrow composed of two unconnected parts, it was not displayed during the experiment.

##### 4.5. Pointing to the tip of a bicolor arrow

###### Colors separated by a sharp edge

In each trial of this experiment, Davida was shown a bicolor arrow and asked to use the computer mouse to move a small round cursor and click as precisely as possible on the tip of the arrow. The arrow was 368 x 35 pixels. The part of the arrow in the same color as the tip was 168 pixels long (e.g., the green part in Figure S17a) and the part in the other color 200 pixels long (e.g., the blue part in Figure S17a). There were 2 presentations of 12 different stimuli composed of different colors (black-blue, black-green, black-red, black-yellow, blue-black, blue-green, blue-red, blue-yellow, red-black, red-green, red-yellow) in 4 different orientations (up, down, left, right) for a total of 96 stimuli. In this task, Davida clicked approximately (i.e. less than 50 pixels away) on the tip of the arrow in 11.4% of the trials, approximately to the place the tip of the arrow would have been if the whole arrow (368 pixels) were rotated by 90, 180 or 270 degrees in 10.4% of the trials, and approximately the place the tip would have been if only the colored part of the arrow of the same color as the tip (168 pixels) had been rotated by 90, 180 or 270 degrees in 78.12% of the trials.

**Fig. S17.**
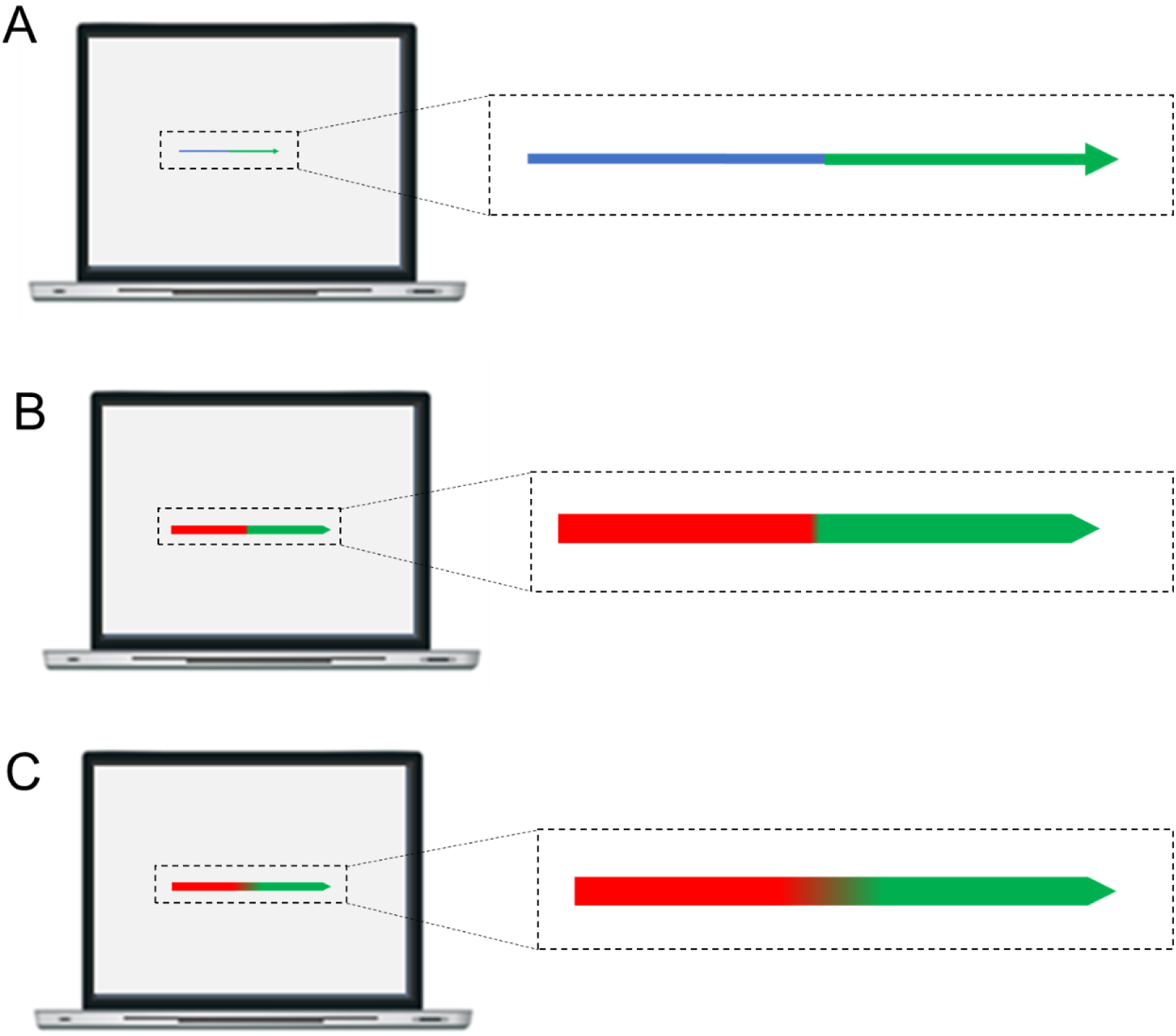
Example of a stimulus used on Experiment 4.5. The dashed lines illustrate the area of the screen selected for the zoom (right). They were not displayed during the experiment.

###### Colors blending more or less progressively

In each trial of this experiment, Davida was shown a bicolor arrow (red-green, see Figure S17B and C) and asked to use the computer mouse to move a small round cursor and click as precisely as possible on the tip of the arrow. The arrow was 450 x 25 pixels. The two colors were blended into one another over either a short or large ((blending of 8 or 60 pixels, see Figure S17B and C). There were 7 presentations of both arrows in 4 different orientations (up, down, left, right) for a total of 56 stimuli. In this task, Davida clicked approximately (i.e. less than 50 pixels away) on the tip of the arrow in 7.1% and 17% of the trials when the blending occurred over a short and large area, respectively. More importantly, she clicked approximately to the place the tip of the arrow would have been if the whole arrow (450 pixels) were rotated by 90, 180 or 270 degrees in 25% of the trials when the blending occurred over a short area but in 82% of the trials when the blending occurred over a large area and approximately the place the tip would have been if only the colored part of the arrow of the same color as the tip had been rotated by 90, 180 or 270 degrees in 67% of the trials when the blending occurred over a short area but never when the blending occurred over a larger area. The Movie S10, online, is a recording of Davida performing this task and illustrate her response profile.

##### 4.6. Pointing to the tip of a rectangle connected or not to another one

In each trial of this task Davida saw two red or two black rectangles either separated from each other by three pixels (Figure S18A, B) or connected by a 4-pixels line (Figure S18C, D), oriented toward the left, right, upper or lower side of the screen, and was asked to use the computer mouse to move a small round cursor and click as precisely as possible on the little “indent” at the extremity of one of the rectangles. When the rectangles were separated, Davida pointed either on the correct location of the indent (1/21 trials), or where the indent would have been if only the rectangle comprising the indent was rotated by 90 degrees (23.8%), 180 degrees (14%) or 270 degrees (57%). When the rectangles were connected by the thin line, Davida pointed either on the correct location of the indent (3/21 trials), or where the indent would have been if the whole shape composed of the two connected rectangles was rotated by 180 degrees (86%). The Movie S9 (Part II), online, is a recording of Davida performing this task and illustrate her response profile.

**Fig. S18.**
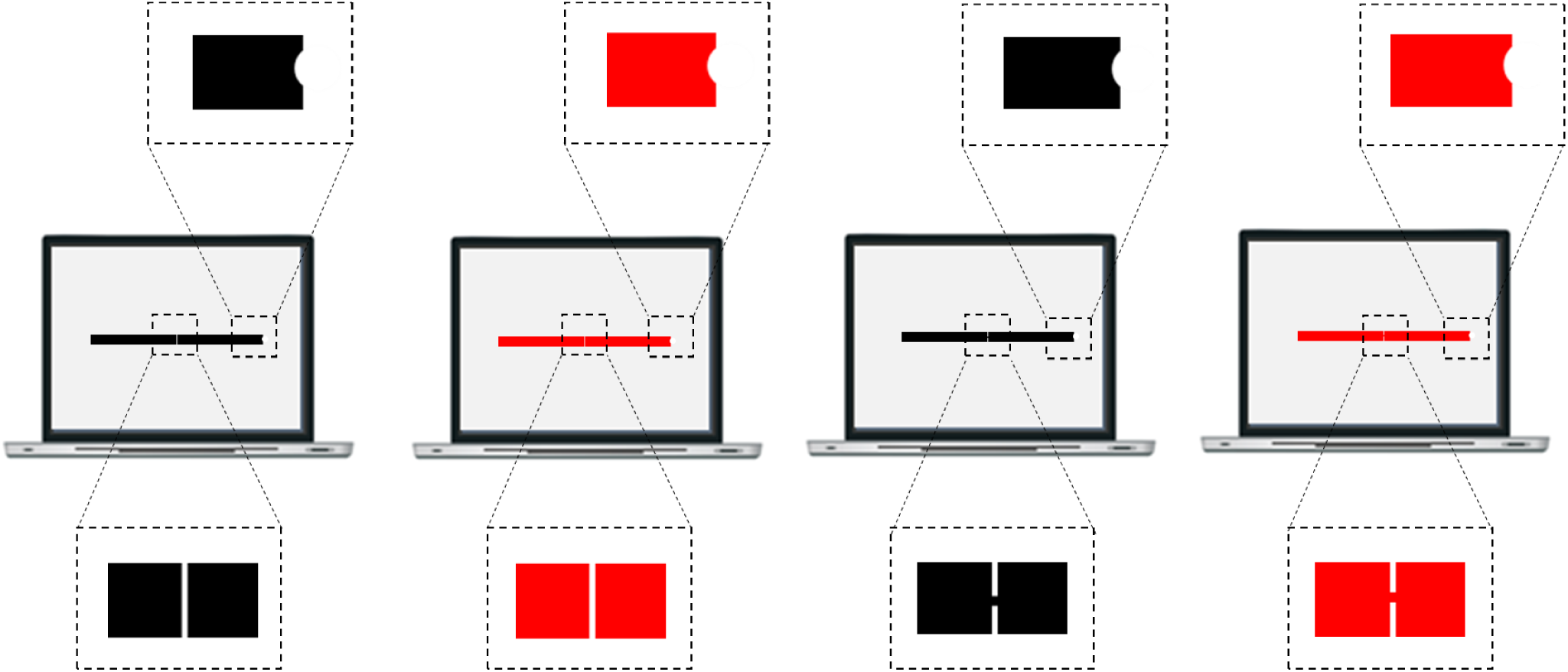
Stimuli used in Experiment 4.6. The dashed lines illustrate the areas of the screen selected for the zoom; they were not displayed during the experiment.

##### 4.7. Counting tasks

In each of the 48 trials of a first task, Davida was presented for 500 ms with a display composed of one, two or three black dots located above a white large square outlined in black ink (Figure S19A-C) or above the same square and a rectangle filled in black ink that would overlap with the position of the black dots if it were rotated by 90 degrees (Figure S19D-F). Davida was asked to name the number of small dots. In each of the 48 trials of a second task, Davida was presented for 500 ms with a display composed of one, two or three black dots displayed within a circle (Figure S19 G-I) or within a circle and a large black rectangle that would overlap with the position of the black dots if it were rotated by 90 degrees (Figure S19J-L). She was asked to name the number of small dots. We counted the number of correct responses by condition in both tasks. In both tasks, Davida was able to correctly report the number of dots when the dots were displayed without the black rectangle (100% RC) but erred in almost all trials in which the display included the black rectangle (12.5 and 25% correct responses in the first and second task).

**Fig. S19.**
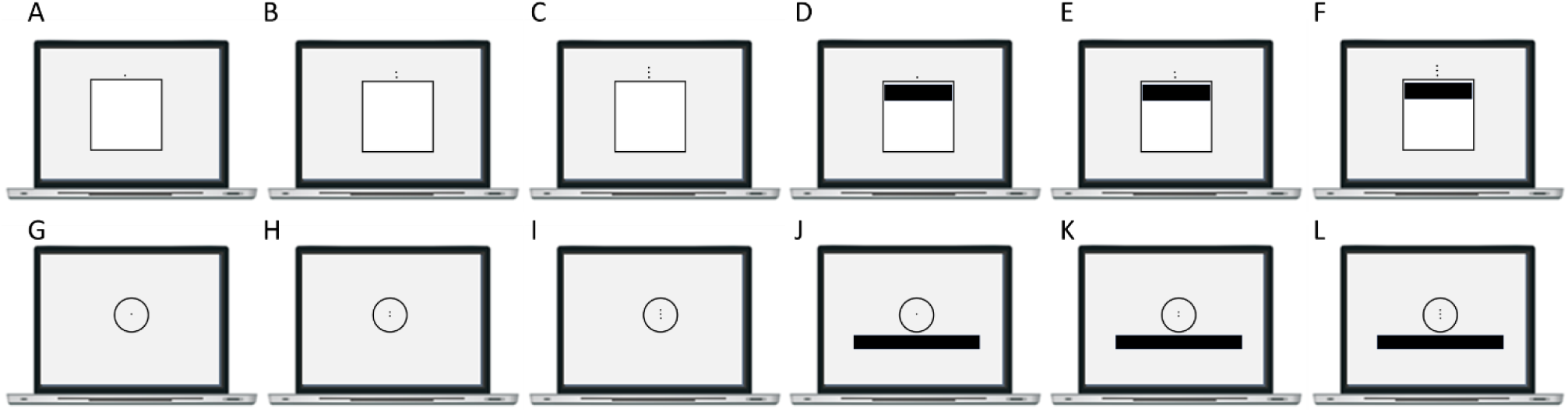
Stimuli used in Experiment 4.7.

##### 4.8. Copying stimuli composed of unconnected parts

Davida was shown 20 aligned dots (Figure S20, A), 10 aligned short horizontal lines (Figure S20, B), 20 aligned short lines vertical lines (Figure S20, C) or 10 aligned lines depicted in different orientations (Figure S20, D) and asked to simply copy what she sees. There were 5 trials in each of the four conditions. With the dotted line, Davida copied accurately the position of every dot in 100% of the trials. In the other conditions, Davida typically erred in reproducing the orientation of the local line segments but reproduced accurately the orientation of the global shape in 100% of the trials.

**Fig. S20.**
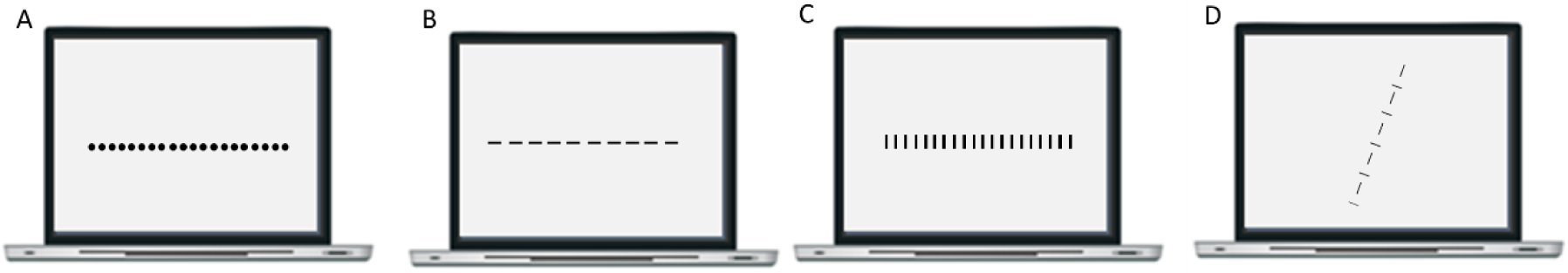
Stimuli used in Experiment 4.8.

##### 4.9. Judging the orientation of stimuli composed of unconnected dots

Davida was shown arrows implied by a series of unconnected small dots (Figure S21) and asked to name the orientation (left, right, up, down) in which that arrow was pointing. Davida named the correct orientation in 20/20 of the trials.

**Fig. S21.**
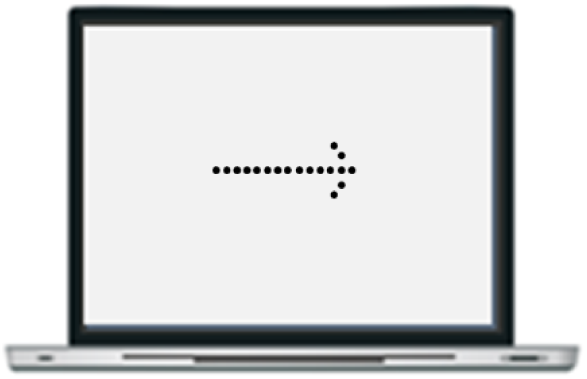
Stimuli used in Experiment 4.9.

##### 4.10. Pointing to the tip of an arrow composed of unconnected dots or of connected elements of different colors

In each trial of the first experiment Davida was shown an arrow implied by a series of unconnected small dots within an arrow composed of solid black lines (Figure S22A) and asked to use the computer mouse to move a small round cursor and click as precisely as possible on either the dot placed at the tip of the dotted arrow (16 first trials) or at the tip of the arrow made of solid black lines (16 other trials). The arrows were displayed in four different orientations (left, right, up, down). Davida was perfect in locating the tip of the dotted arrow but erred on 14/16 of the trials when asked to point to the tip of the arrow made of solid black lines. In this condition, she located the tip of the arrow approximately the place it would have been if it were rotated by 90, 180 or 270 degrees in 12.5, 37.5 and 37.5 % of the trials, respectively. The Movie S11, online, is a recording of Davida performing this task and illustrate her response profile.

In each trial of the second experiment, one of three types of a large black arrow (made of connected lines, of small or large unconnected dots, Figure S22 B-D) was displayed at the center of the computer screen pointing right, down, left or up for 200 ms; Davida was asked to use the computer mouse to move a small round cursor and click as precisely as possible on the tip of the arrow. When the arrow was composed of solid black lines (Figure S22B) Davida pointed to the tip of the arrow in 4/20 trials and often mislocated the tip at approximately the place it would have been if the arrow were rotated by 90 degrees (4/20), 180 degrees (10/20) or 270 degrees (4/20). In contrast, Davida made no errors when the arrows were composed of unconnected small or large dots (Figure S22C, D). The Movie S12, online, is a recording of Davida performing this task and illustrate her response profile.

In each trial of the third experiment, one large arrow composed of segments of different colors (Figure S22 E) was displayed at the center of the computer screen pointing right, down, left or up for 200 ms and Davida was asked to use the computer mouse to move a small round cursor and click as precisely as possible on the tip of the arrow. There were 40 trials, in which Davida made no error.

**Fig. S22.**
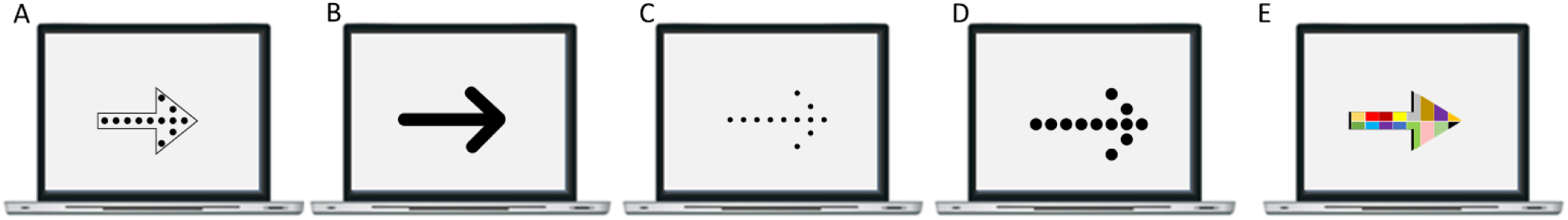
Stimuli used in Experiment 4.10.

##### 4.11. Stimulus–response compatibility task with dotted arrows

In each trial the first experiment, a fixation point was presented at the center of the screen for 2 sec, then the screen was cleared for 1 sec, and then a stimulus was displayed until a response was recorded. The stimuli were either a filled circle or a filled square displayed at the center of the computer screen and, below it, an arrow made of unconnected small black dots pointing either toward the left or toward the right (see Figure S23A-D). Thus, there were 4 different stimuli (2 shapes x 2 arrow orientations). Davida was asked to press a button on the keyboard (the “z” key) as fast as possible with her left index finger when she saw a circle and with her right index finger (the “m” key) when she saw a square, while ignoring the arrow. The experiment included 120 stimuli (30 repetition of each stimulus). To test whether Davida was implicitly influenced by the orientation of the arrow, we conducted analyses to compare Davida’s response latencies for trials in which the arrow pointed in the direction of the hand associated with the correct answer (congruent displays) and for trials in which the arrow pointed in the direction of the hand associated with the incorrect answer (incongruent displays). Response accuracy and response latency were analyzed separately. Response latency analyses were carried out over correct responses only. The distribution of Davida’s response latencies was homogeneous; there were no exceedingly fast (i.e., faster than 200 ms) or slow (slower than 1000 ms) responses. Davida was slightly less accurate for incongruent than congruent trials (95% and 98%), and an independent sample t-test carried out on Davida’s reaction latencies (RL) revealed that she was significantly faster on congruent (Mean = 521 ms; SD = 10.8 ms) than incongruent trials (Mean = 571 ms; SD = 11.9 ms) (tuni(114) = 2.33, p = 0.01).

In the second experiment, Davida was presented 15 times with each of the 8 displays depicted in Figure S23 A-H (randomly), and her task was the same as in the previous experiment. The aim was to replicate the results of the first experiment and of a previous stimulus–response compatibility experiment with solid arrows (see supplemental material and methods 1.11), and to compare in the same experiment the stimulus-response compatibility effect induced by a dotted arrow (S23 A-D) versus an arrow composed of solid black lines (S23 E-H). The analyses were as reported before. Davida was equally accurate for the incongruent and the congruent trials in both conditions (full arrow: 100%; dotted arrow: 96.7%). Latency analyses were conducted on the correct responses after responses faster than 200 ms (0%) and slower than 1000 ms (1 congruent and 1 incongruent trial for the solid arrow condition; 1 congruent and 2 incongruent trials in the dotted arrow condition) were excluded(Hommel, 1993). The results of two independent sample t-tests carried out over Davida’s RLs showed that she was significantly faster to respond to congruent (Mean = 517 ms; SD = 10 ms) than incongruent trials (Mean = 573 ms; SD = 10.1 ms) (tuni(53) = 2.05, p = 0.02) for the dotted arrows, but not for the solid arrows (Congruent: Mean = 525 ms; SD = 9.1 ms; Incongruent: Mean = 557 ms; SD = 14 ms; tuni(56) = 1.02).

**Fig. S23.**
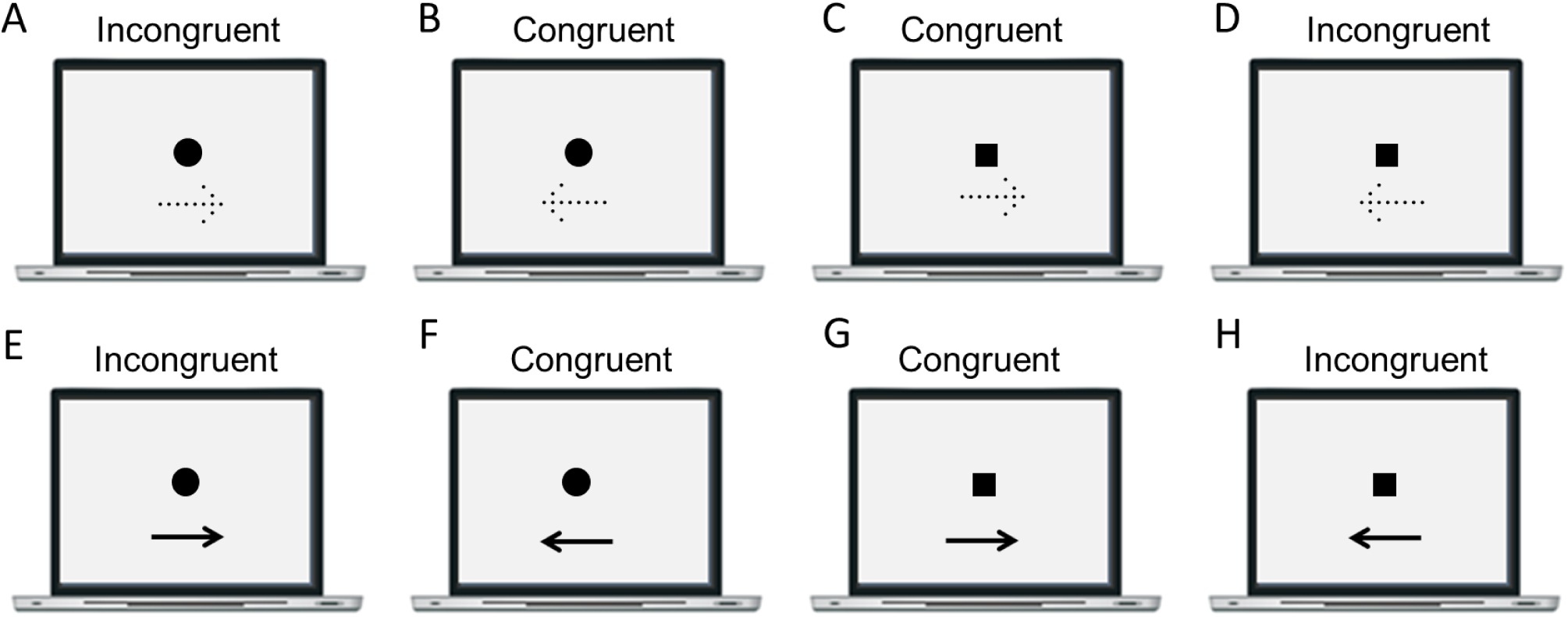
Stimuli used in Experiment 4.11. Stimuli were categorized as congruent (B, C, F, G) when the arrow points in the direction of the hand associated to the correct answer (left for a circle, right for a square) and incongruent (A, D, E, H) when the arrow points in the direction opposite to the hand associated to the correct answer (right for a circle, left for a square).

##### 4.12. Grouping by proximity

In a first experiment, Davida was presented five times for 100 ms with each of the 4 displays depicted in Figure S24 and asked to tell what she saw. In a second experiment the stimuli were displayed for only 16 msec. In both experiments, Davida systematically reported seeing vertical lines of dots when shown the stimuli depicted in Figure S24 A and C, and horizontal lines of dots when shown the stimuli depicted in Figure S24 B and D. Thus, Davida perceives accurately the orientation of lines made of unconnected elements, even when this type of stimulus is presented so briefly that it is unlikely that her response follows a conscious reconstruction of the orientation of the stimulus based on an analysis of the position of its parts.

**Fig. S24.**
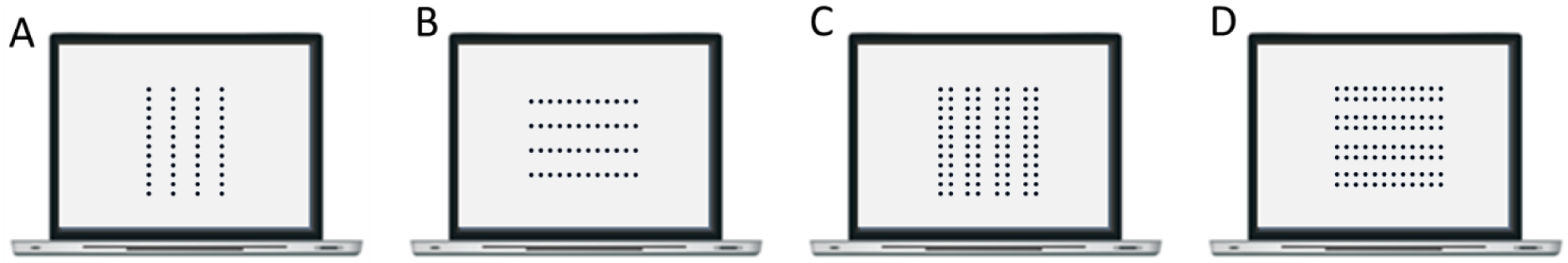
Stimuli used in Experiment 4.12.

##### 4.13. Copying stimuli composed of connected elements of different colors

In the first experiment, Davida was presented for 200 ms with a line (Figure S25 A) or an arrow (Figure S25 B) composed of segments of different colors and asked to copy the outline of stimuli with black ink on a separate sheet of paper. The line was either vertical or horizontal (six stimuli of each) and the arrow was either oriented to the right, left, up or down (three stimuli of each). In the second experiment, she was shown the same stimuli for only 16 msec. Davida performed these tasks easily and flawlessly.

**Fig. S25.**
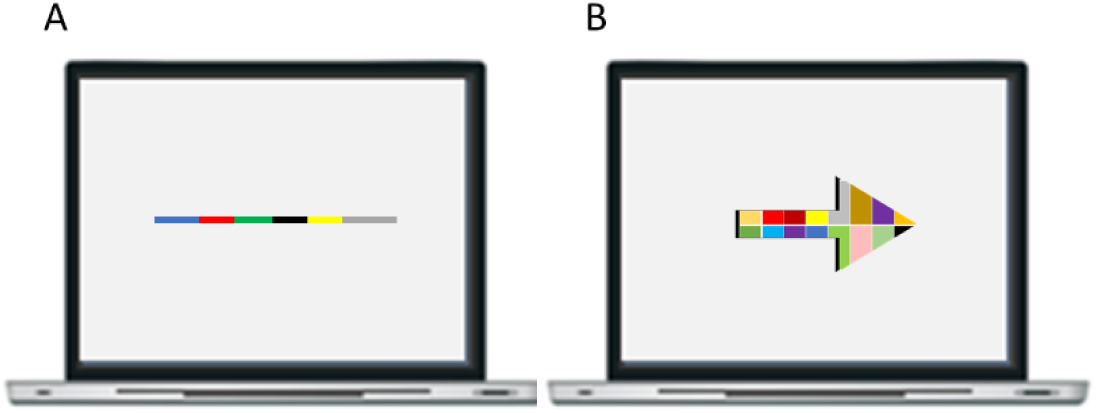
Stimuli used on Experiment 4.12.

##### 4.14. Naming letters composed of unconnected elements or connected elements of different colors

In each of the trials of the first experiment one of four possible orientation-sensitive letters (p, b, d or q) either drawn in black ink (Figure S26 A), composed of connected parts of different colors (Figure S26 B) or composed of small black dots (Figure S26 C) and subtending 166 x 240 pixels was shown at the center of the computer screen. Davida was asked to name the letter, which was displayed for as long as she needed. There were 20 trials by condition. Davida named accurately 2/20 letters drawn in black ink (Figure S26 A) but named easily and flawlessly (20/20) the letters composed of connected parts of different colors (Figure S26 B) and those composed of small black dots (Figure S26 C).

In the second experiment, the same stimuli were displayed for only 16 msec. Davida named accurately 0/20 letters drawn in black ink (Figure S26 A) but named easily and flawlessly the letters composed of connected parts of different colors (Figure S26 B) and those composed of small dots (Figure S26 C).

**Fig. S26.**
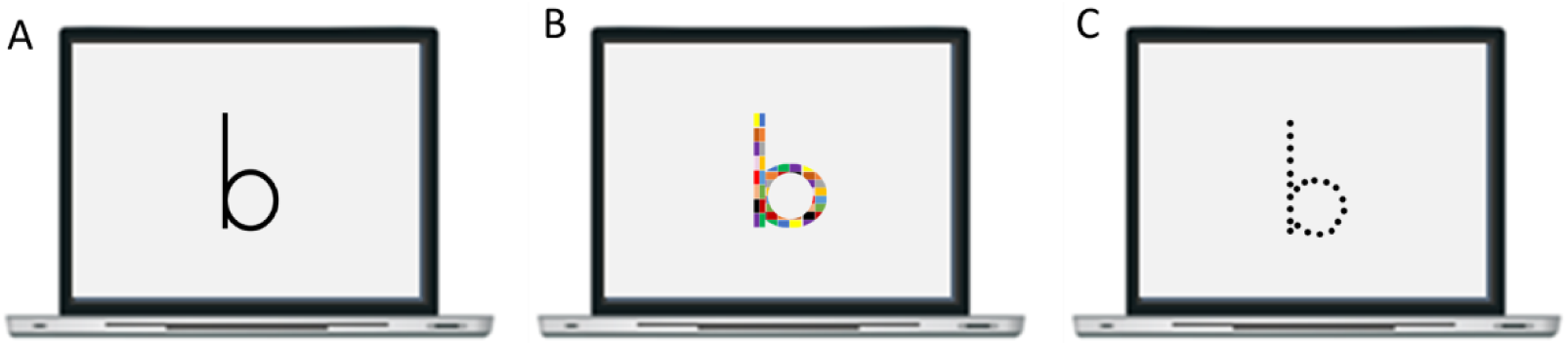
Examples of stimuli used in Experiment 4.14.

#### Materials and methods: set of results §5

##### 5.1. Copying a tilted asymmetrical shape

Davida was randomly presented with one of two asymmetrical shapes (see Figure S27) in one of 16 possible orientations (15, 30, 60, 75, 105, 120, 150, 165, 195, 210, 240, 255, 285, 300, 330 and 345 degrees) and was asked to copy it as precisely as possible on a separate sheet of paper. These shapes were ± 7 x 3 degrees of visual angle and drawn in black ink on white background. Davida was asked to copy a total of 296 shapes across six separate sessions. (in two of these sessions these stimuli were intermixed with stimuli displayed in lower contrast or blurred but Davida’s results for this type of stimuli will not be reported here). The Movie S13, online, is a recording of Davida performing this task and illustrate her response profile. Davida copied 13/296 stimuli accurately (4.4 %) and made the 7 types of errors displayed in Figure S27 on the other trials (see also Figure 4 and 6 in the main text).

**Fig. S27.**
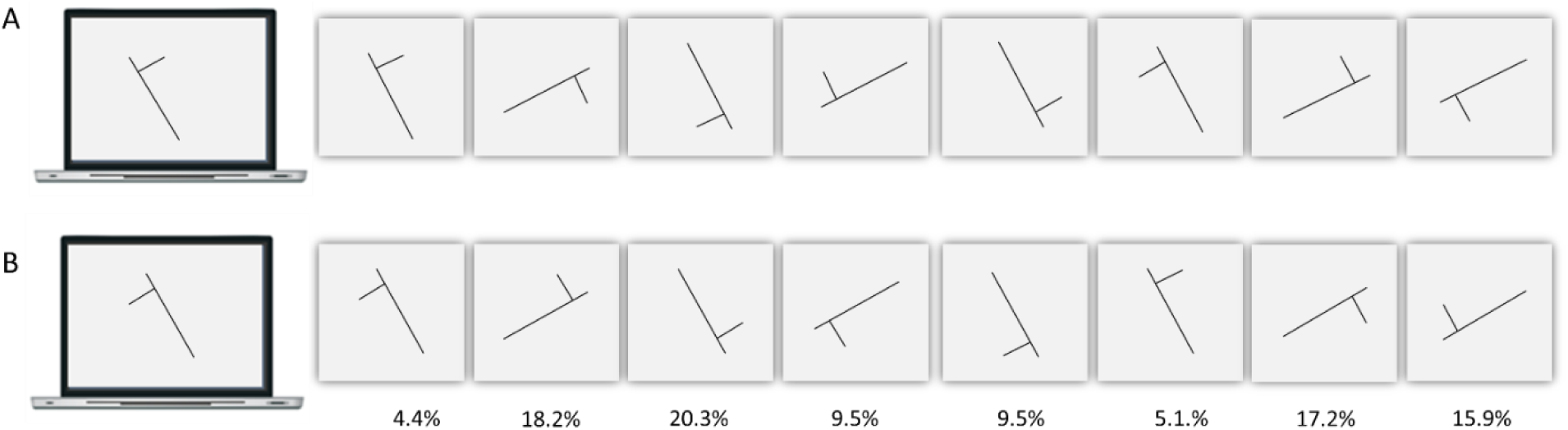
A, B. Examples of the two stimuli used in Experiment 5.1 (shown here tilted 330 degrees), an illustration of the 8 types of responses given by Davida for these types of stimuli, and their corresponding percentage. See Figure 6, in the main text, for more detail on these errors.

##### 5.2. Copying another tilted asymmetrical shape

Davida was randomly shown one of two asymmetrical shapes, the long axis of which was tilted 15 degrees from the vertical or horizontal (see Figure S28) in one of 8 possible orientation (15, 75, 105, 165, 195, 255, 285, and 345 degrees) and was asked to copy it as precisely as possible on a separate sheet of paper. These shapes were ± 7 x .05 degrees of visual angle and drawn in black ink on white background. Davida was asked to copy a total of 64 shapes. The Movie S14, online, is a recording of Davida performing this task and illustrate her response profile. Davida copied 1/64 stimulus accurately (1.6 %) and made the 7 types of errors displayed in Figure S28 on the other trials.

**Fig. S28.**
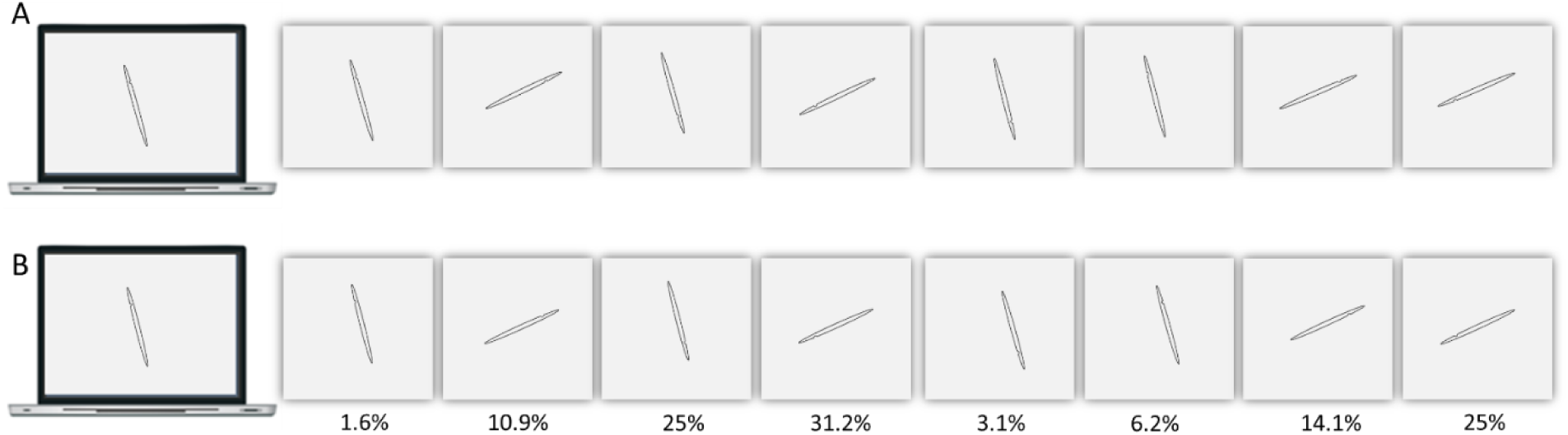
A, B. Examples of the two stimuli used in Experiment 5.2 (shown here tilted 345 degrees), an illustration of the 8 types of responses given by Davida for these types of stimuli, and their corresponding percentage. See Figure 6, in the main text, for more detail on these errors.

##### 5.3. Drawing all that she sees

Davida was randomly presented with one of two asymmetrical shapes (Figure S29) tilted 30 degrees from the vertical or horizontal in one of 8 possible orientations (30, 60, 120, 150, 210, 240, 300, and 330 degrees) for 2 seconds and asked, after each presentation, to draw “all she saw”. These shapes were ± 7 x 3 degrees of visual angle and drawn in black ink on white background. There was one trial by condition (2 shapes, 8 orientations). The Movie S15 illustrates Davida drawing the different orientations that she perceived when shown a similar stimulus. Davida drew each shape in 6.06 different orientations on average, including the correct orientation 81.25 % of the time (13/16) and 7 errors types displayed in Figure S29 on the other trials (see also Figure 3 in the main text) in various proportions but made no other error.

**Fig. S29.**
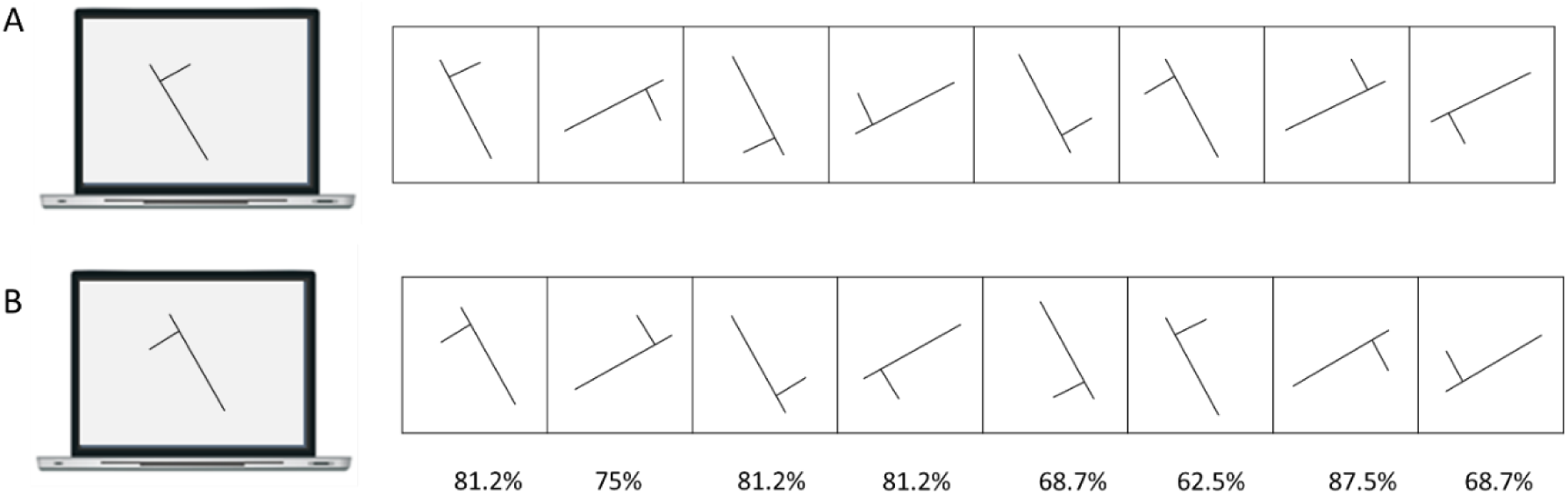
A, B. Examples of the two stimuli used in Experiment 5.3 (shown here tilted 330 degrees), an illustration of the 8 types of responses given by Davida for these types of stimuli, and the corresponding percentage of trials in which this orientation was drawn. See Figure 6, in the main text, for more detail on these errors.

##### 5.4. Tracing a tilted asymmetrical shape

Davida was randomly shown one of two asymmetrical shape (Figure S30) tilted 15 degrees from the vertical or horizontal in one of 8 possible orientations (15, 75, 105, 165, 195, 255, 285, 345 degrees) on a sheet of paper and asked to trace the shape with ink. The Movie S16, online, is a recording of Davida performing this task and illustrate her response profile. On a total of 80 trials, Davida traced the displayed shape only 2.5% of the time (2/80 trials). On the other trials, she inked the sheet of paper as if the shape was transformed by one of the 7 error types displayed on the Figure S30 (see also Figure 4 and 6 in the main text). She made no other type of error.

**Fig. S30.**
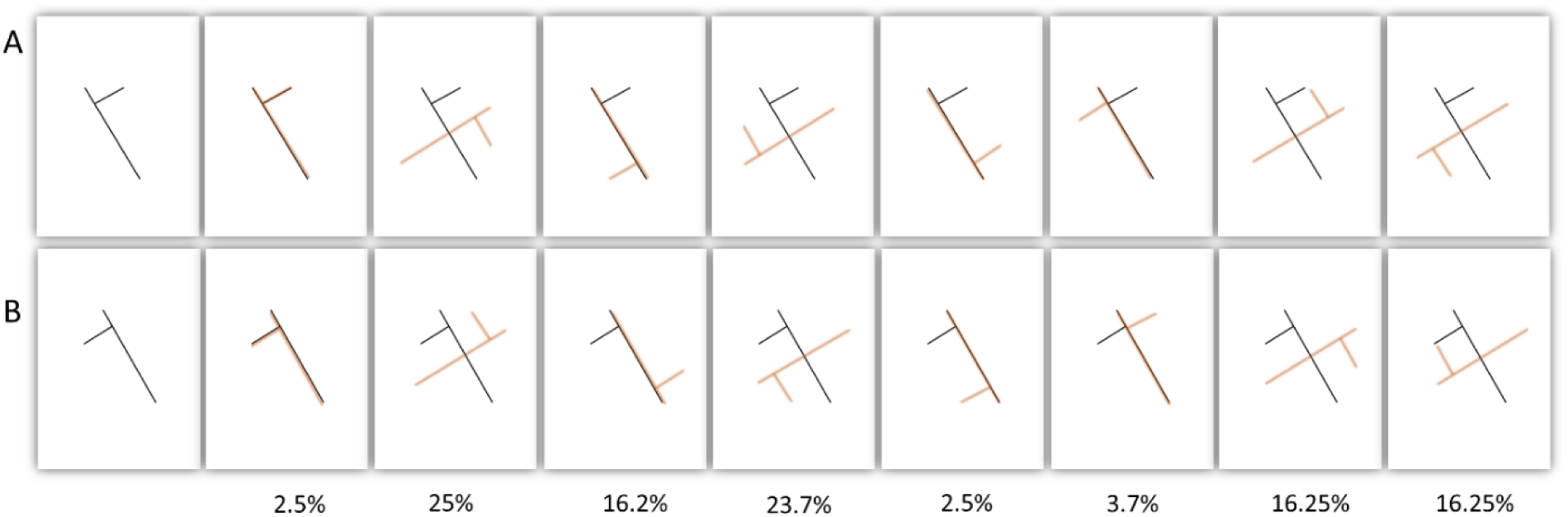
A, B. Examples of the two stimuli used in Experiment 5.4 (shown here tilted 345 degrees), an illustration of the 8 types of responses given by Davida for these types of stimuli (in red ink), and the percentage of trials corresponding to the different types of responses.

##### 5.5. Matching tilted asymmetrical shapes

In this task, Davida was shown two exemplars of a tilted asymmetrical shape displayed on the left and right side of the center of a computer screen (see Figure S31) and asked to report verbally (yes/no) whether she saw the two shapes in the same orientations or not. The longer segment of the shape subtended approximately 5 degrees of visual angle. The shapes were displayed in black ink on white background. The left (target) shape was always tilted 15 degrees counterclockwise from vertical. The right shape could be the same as the target (48 trials, Figure S31A), it could be tilted 10 degrees more or 10 degrees less than the target (i.e., tilted 5 or 25 degrees; 24 trials, Figure S31 B, C), it could be a vertical (12 trials, Figure S31 D) or a horizontal (12 trials, Figure S31 E) mirror reflection of the target; or correspond to any of the 7 error types (Figure S31 F-L) found in Davida for the same type of stimuli in previous experiments (see Supplemental method and Material 5.1 – 5.4, see also Figure 2 in the main text for a discussion of these types of errors). The two shapes were displayed until Davida gave a response; there was no time limit to respond and accuracy was emphasized. Davida was allowed to shift her gaze between the two shapes as many times as needed. Davida made no errors on the identical trials (48/48), on the trials in which the two shapes differed by 10 degrees (24/24) or on the vertical and horizontal mirroring (24/24) (Figure S31 A-E). However, she made numerous errors in all the other conditions (Figure S31 F-L), judging most of the time that the two shapes were displayed in the same orientation.

**Fig. S31.**
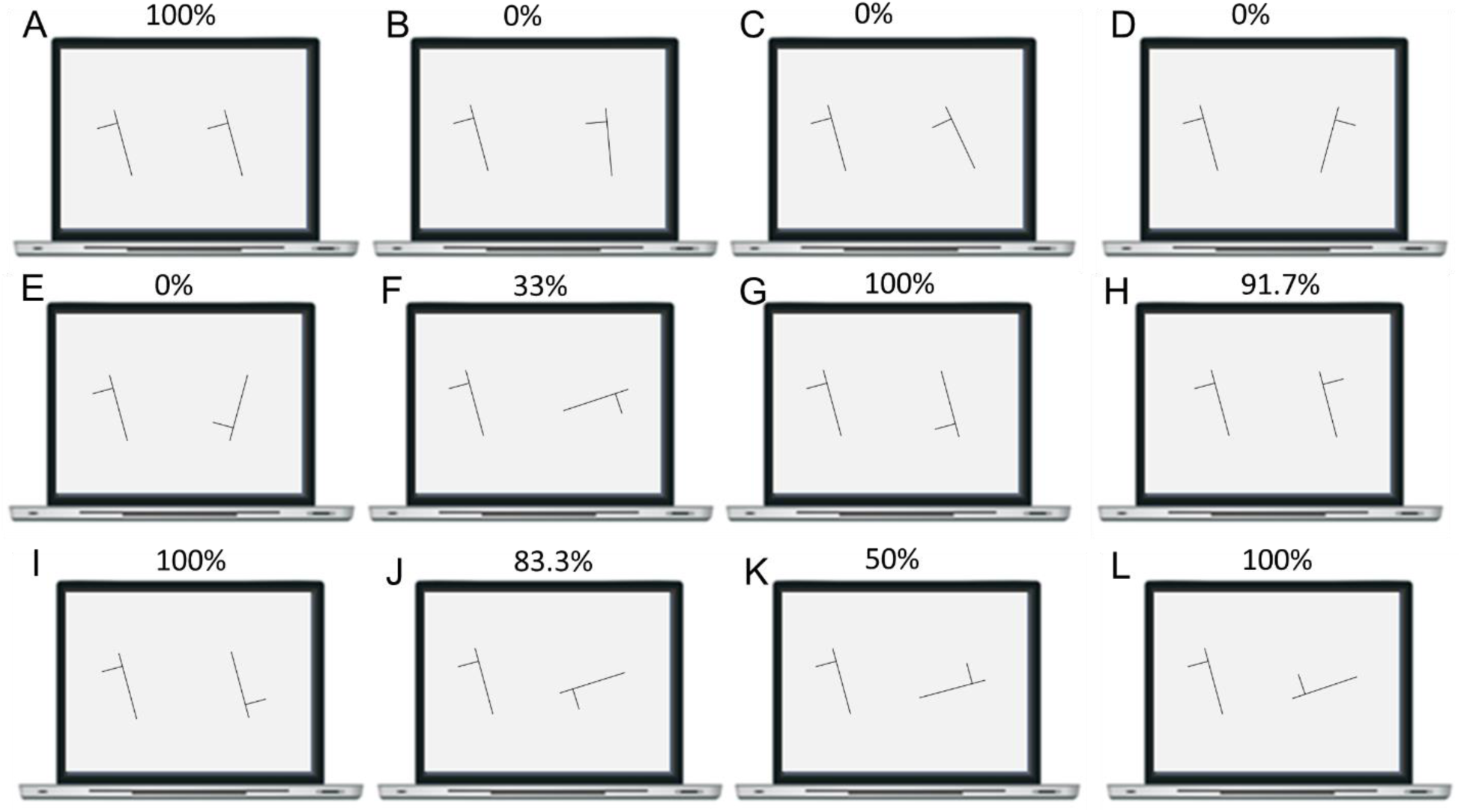
Stimuli used in Experiment 5.5 (shown here tilted 345 degrees), and for each of them the percentage of trials in which Davida reported seeing the two shapes in the same orientations.

##### 5.6. Matching tilted shapes to probe the nature of the long axis

This task aimed at discriminating the role of three geometrical properties of a shape for determining the shape’s long axis (Figure S32 A): the axis corresponding to the shape’s longest straight segment (Chaisilprungraung et al., 2019), the axis relating the two most distant points of the shape (longest span axis; Sekuler & Swimmer, 2000), or the axis that minimizes the sum of squared distances to all points of the shape (the axis of least second moment; Haralick & Shapiro, 1991).

In each of the 24 trials of this shape association task, Davida was shown a shape (target) and, below, an array of three shapes displayed at very low contrast with respect to the white background (the probes). The contrast of the probes was determined by Davida herself and corresponded to a very light grey level (RGB of 252) at which she reported seeing these stimuli, including their orientation, perfectly, and their orientation did not appear to change over time (see the main text and supplemental method and 6.3 – 6.9, for more detail on the effect of luminance contrast on Davida’s perception).The three probes were mirror reflections of the target across an axis aligned on either the shapes’ longest straight segment (Figure S32 B), (B) the axis of least second moment (Figure S32 C), or the longest span axis (Figure S32D). Across the 24 trials, the target was displayed twice as displayed the Figure S29 and twice in each of 11 rotations of that stimulus by steps of 30 degrees (30, 60, 90, 120, 150, 180, 210, 240, 270, 300, 330). Davida’s task was to point to the probe corresponding to a possible orientation of the target (that she sees in several orientations). Davida pointed to the probe corresponding to a mirror reflection of the target across the shape’s longest straight segment (Figure S32 B) in 100% of the trials.

**Fig. S32.**
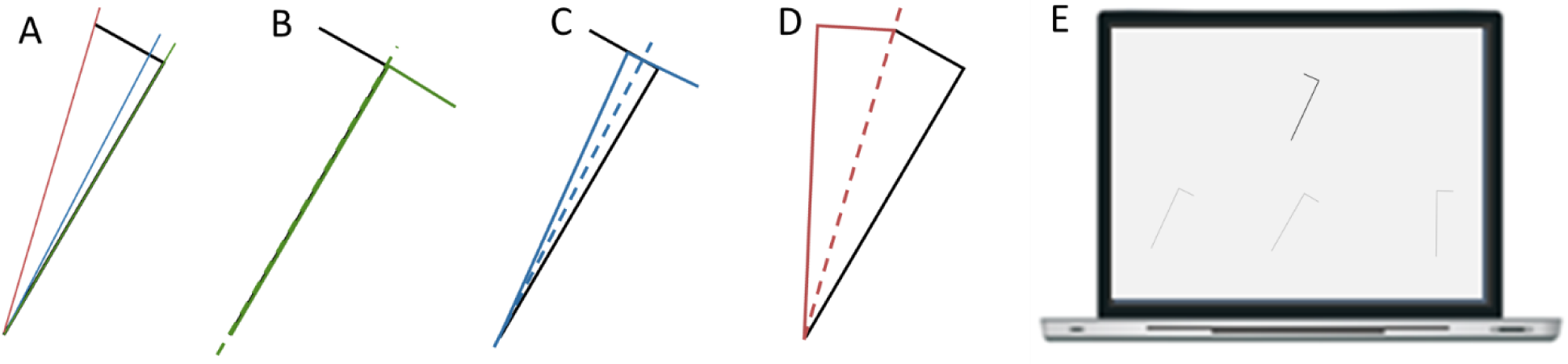
A. Illustration of the shape’s longest straight segment (green), longest span axis (red) and axis of least second moment (blue). B. Mirror reflection (in green) of the shape (in black) across an axis aligned on the shape’s longest straight segment. C. Mirror reflection (in blue) of the shape (in black) across an axis aligned on the shape’s axis of least second moment. D. Mirror reflection (in red) of the shape (in black) across an axis aligned on the shape’s longest span axis. E. Illustration of the experimental display. Top: the target. Bottom: mirror reflections of the target across an axis determined by the shape’s longest straight segment (left), axis of least second moment (center) and longest span axis (right). The contrast of the probes was largely increased in this example to make them more visible.

##### 5.7. Exploring the “center” of elongated shapes’ representational frame

Here, we aimed at discriminating whether the center of the representational frame (i.e., the intersection of the shapes’ long and secondary axes) of an elongated shape is the center of the shape’s long axis (red dot in Figure S33A) or the intersection of the shape’s long axis and a secondary axis passing through the shapes’ centroid (mean coordinate of all the points in the shape, green dot in Figure S33A). To discriminate between these possibilities, we re-analyzed Davida’s responses in the 560 trials of an experiment in which she was asked to localize the tip of a large black arrow (Supplemental Methods and Results 19, experiment 1). For each of Davida’s error in this experiment (N = 550), we calculated the distance between the coordinates of Davida’s response (where she reported seeing the tip of the arrow) and the place where the tip of the arrow would have been if the center of the representational frame was the center of the long axis (Figure S33 A, red) or of the arrow’s centroid (Figure S33A, green). As shown in Figure S33, she located the tip of the arrow (illustrated by the black triangle in Figure S33B-E) closer on average (38 pixels, 6 mm; ± 0.6 degrees of visual angle) from where it would have been if the arrow were rotated by 90, 180 or 270 degrees around the center of the long axis (illustrated by the red dot in Figure S33B-E), than from where it would have been if the center of the representational frame was the arrow’s centroid (84 pixels; 14 mm; ± 1.4 degrees of visual angle; illustrated by the green dot in Figure S33B-E).

**Fig. S33.**
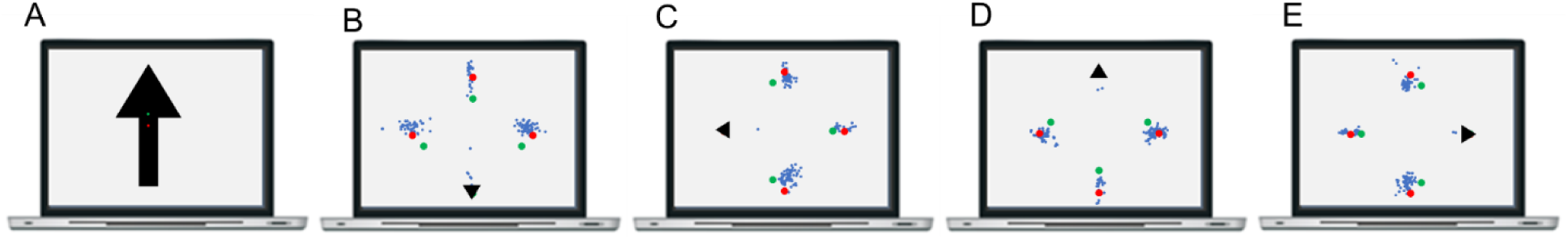
A. Illustration of the center of the arrow’s long axis (red dot) and of the arrow’s centroid (green dot). B-E. Each blue dot represents the coordinates of one attempt of Davida to localize the tip (illustrated by the black triangle) of an arrow pointing down (B), left (C), up (D) or right (E). The red and green dots illustrate where the tip of the arrow would have been if the center of the representational frame were the center of the long axis (red) or of the arrow’s centroid (green).

##### 5.8. Exploring the “center” of tilted asymmetrical elongated shapes’ representational frame

This experiment aimed at replicating the results of Experiment 5.7 with a new type of stimulus. To discriminate whether the center of the representational frame (i.e., the intersection of the shapes’ long and secondary axes) of an elongated asymmetrical shape is the center of the shape’s long axis (red dot in Figure S34A, B) or the shapes’ centroid (green dot in Figure S34A, B) this experiment relied on the fact that Davida almost systematically perceives elongated shapes in several orientations, in which the shapes’ main (elongation) axis is always either accurately oriented or rotated by 90 degrees. We investigated whether, in this case, the main (elongation) axis of these two orthogonal percepts intersect at the level of the center of the shape’s centroid (Figure S34 C) or at the center of the shape’s elongation axis (Figure S34 D).

In each trial of this experiment, Davida was presented with either a tilted asymmetrical shape or a simple line corresponding to the shape’s longest straight segment (Figure S34 A, B) and was asked to either use the computer mouse to move a small round cursor and click as precisely as possible on the place where she saw two lines crossing each other or to press the space bar if she did not see any line crossing. There was no time constraint. Both stimuli were displayed three times in 12 different orientations (0 to 330 degrees by steps of 30 degrees) for a total of 72 trials (2 stimuli x 12 orientations x 3 repetitions).

If the center of elongated shapes’ representational frame is defined by the shapes’ centroid, then, when presented with the asymmetrical shape, Davida should perceive lines intersecting at the level of that shape’s centroid (the green dot in Figure S34 C). In addition, she should perceive lines intersecting at a different place along the main axis of the simple line and of the asymmetrical stimuli (compare the position of the green dot in Figure S34 A and B). In contrast, if the center of elongated shapes’ representational frame is defined by the center of the shapes’ most elongated part (the red dot in Figure S34 A, D), then, she should perceive lines intersecting on the center of the shapes’ most elongated part (Figure S34 D) and there should be no difference between the simple line and of the asymmetrical stimuli.

During the task, Davida never used the space bar to indicate that she did not see lines crossing. The coordinates at which she indicated seeing two lines crossing was on average at 14.3 pixels from the center/centroid of the line stimulus (2 mm, ± 0.2 degrees of visual angle) and at 14.9 pixels (2 mm, ± 0.2 degrees of visual angle) from the center/but 45 pixels from the centroid of the asymmetrical elongated shape. Thus, the center of the elongated shapes’ representational frame is defined by the center of their longest straight part.

**Fig. S34.**
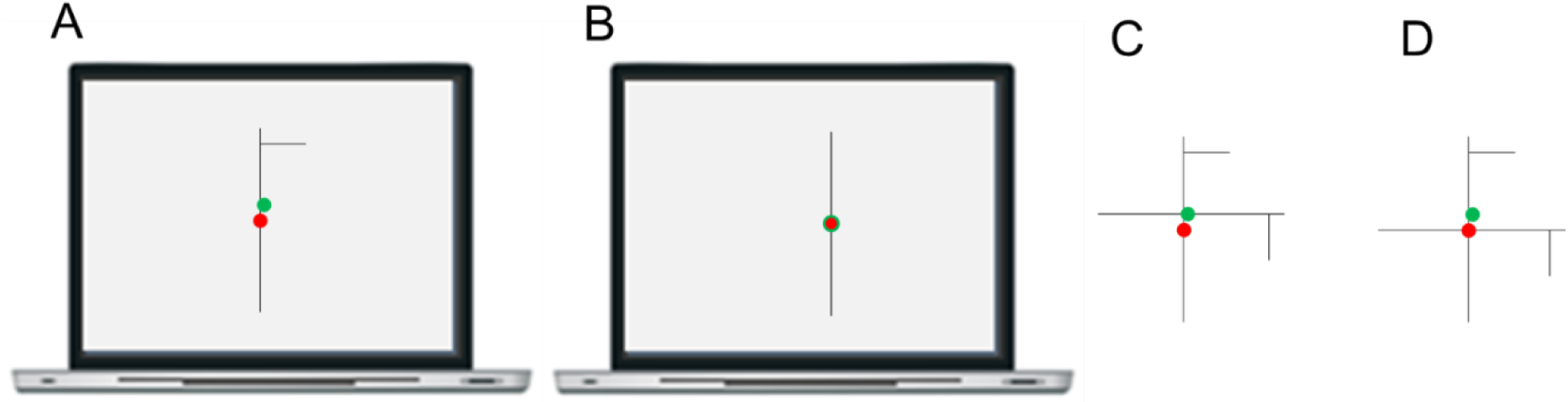
A, B. Illustration of the stimuli used in Experiment 5.8 and of the position of the center of their longest straight part (the red dot) and of their centroid (the green dot). The center and centroid of the simple line being located at the same position, they are depicted as overlapping circles. C, D. Illustrations of possible percepts the center of these shapes’ representational frames is defined by their centroid (C) or at the level of the center of their most elongated part (D).

##### 5.9. The structure of the representational frame: circles

Here, we aimed at characterizing the representational frame for non-elongated symmetrical shapes (Figure S35). In each trial of this experiment Davida was shown a large black circle (640 x 640 pixels; 10.32 degrees of visual angle) with a small semicircular indent of the same color as the background (see Figure S35) for as long as needed, and she was asked to use the computer mouse to move a small round cursor and click as precisely as possible on the place “where she sees the indent” before pressing the space bar to launch the next trial. She was free to click any number of times on the stimulus. The circle was displayed with the indent rotated either 10 (e.g., Figure S35), 20, 30, 60, 70, 80, 150, 160, 170, 190, 200, 210, 240, 250, 260, 280, 290, 300, 330, 340 or 350 degrees clockwise from the vertical midline. Davida acted on two or three stimuli at each orientation for a total of 47 stimuli (the task had to be aborted due to time constraint). Across the trials, Davida reported seeing the indent at either two (1/47), three (35/47) or four (11/47) different locations on the screen, providing a total of 152 coordinates. The Movie S17, online, is a recording of Davida performing this task and illustrate her response profile. Among these responses, 30.9% consisted in localizing the indent at its’ correct location (i.e., Davida clicked at less than1 degree of visual angle, 62 pixels, of the center of the indent). Her errors consisted mainly in localizing the ident erroneously at less than1 degree of visual angle from where it would have been if the circle had been rotated by 90 (19.7%), 180 (26.9%) or 270 (16.4%) degrees.

Among the 9 other errors, one error (0.6%) consisted in localizing the ident erroneously at less than1 degree of visual angle from where it would have been if the circle had been mirrored across a vertical axis, and 8 (5.2%) errors were at a distance of more than 1 degree of visual angle from any interpretable landmark. The results thus indicated the existence of a frame composed by the shape’s axis of symmetry and it’s perpendicular, intersecting either at the center of the axis of symmetry or at the centroid of the shape.

**Fig. S35.**
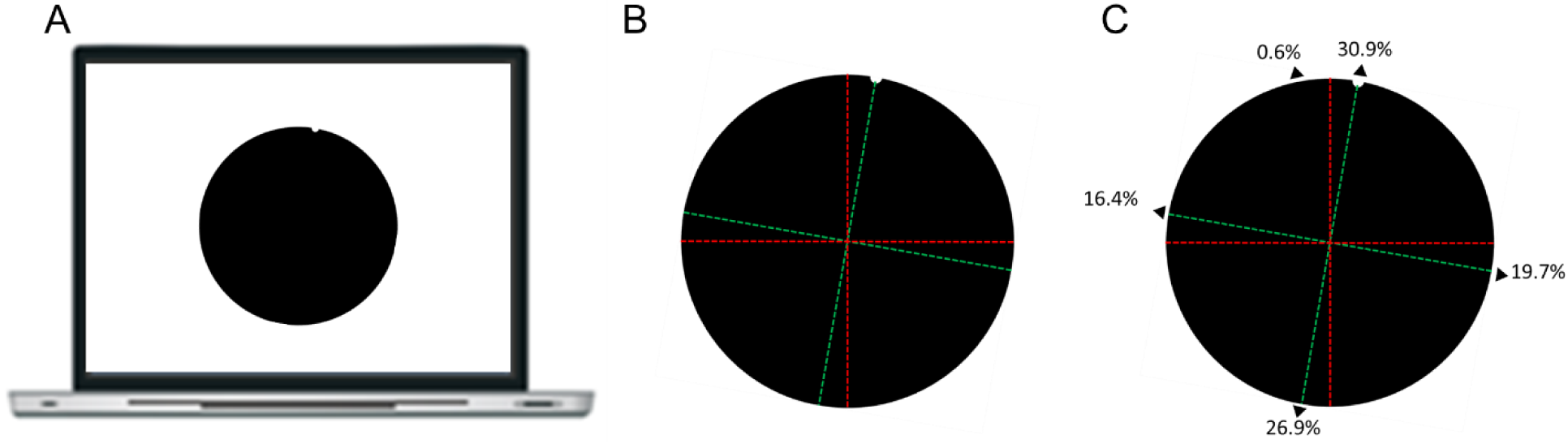
A. Illustration of a stimulus used in Experiment 5.9 oriented 10 degrees from the vertical clockwise. B. Illustration of a shape-centered representational frame composed of an axis aligned on the circle’s axis of symmetry and of its perpendicular (in green), and of a shape-centered representational frame composed of extrinsic vertical and horizontal axes (in red). C. The percentages refer to the proportion of Davida’s responses corresponding to the different locations identified by the black triangles.

##### 5.10. The structure of the representational frame: semi-circles and arcs

This experiment aimed at confirming the finding from Experiment 5.9 with a new type of stimulus and at gathering data pertaining to the question of the center of the representational frame for symmetrical shapes. We first presented Davida with 40 semicircles (see Figure S36 A) on a sheet of paper and asked her to trace the shape with ink. The semi-circles were oriented 30 (Figure 36 A), 60, 120, 150, 210, 240, 300 or 330 degrees and there were 5 trials per orientation. Davida traced the displayed shape accurately (Figure S36B) in 10% of the trials (4/40). In most of the other trials, she inked the sheet of paper as if the shape were rotated by 90 (Figure S36C), 180 (Figure S36D) or 270 (Figure S36E) degrees around the shape’s centroid (illustrated by the green dot in Figure 36 A-F). She also made one error consisting in inking the sheet of paper as if the shape were rotated by 180 degrees around a point located at the intersection of the shape and its’ axis of symmetry (Figure S36F). Then, in each of the 36 trial of the next experiment Davida saw the arc of a circle (corresponding to ± 82 degrees; Figure S36G) on a sheet of paper and asked her to trace the shape with ink. The arcs were oriented 15 (see Figure S36 G), 45, 75, 105, 135, 165, 195, 225, 255, 285, 315, 345 degrees clockwise from the vertical and there were 3 trials per orientation. Davida accurately traced the displayed shape in 5.5% of the trials (Figure S36 H). In the other trials, she inked the sheet of paper as if the shape were rotated by 90 (Figure S36 H), 180 (Figure S36 I) or 270 (Figure S36 K) degrees around the shape’s centroid (illustrated by the green dot in Figure 36 G-K). These results confirmed that Davida’s errors emerged in a representational frame based on these shapes’ axis of symmetry and indicated that center of the frame is the shapes’ centroid (except for one answer in the first experiment, Figure S32 E).

**Fig. S36.**
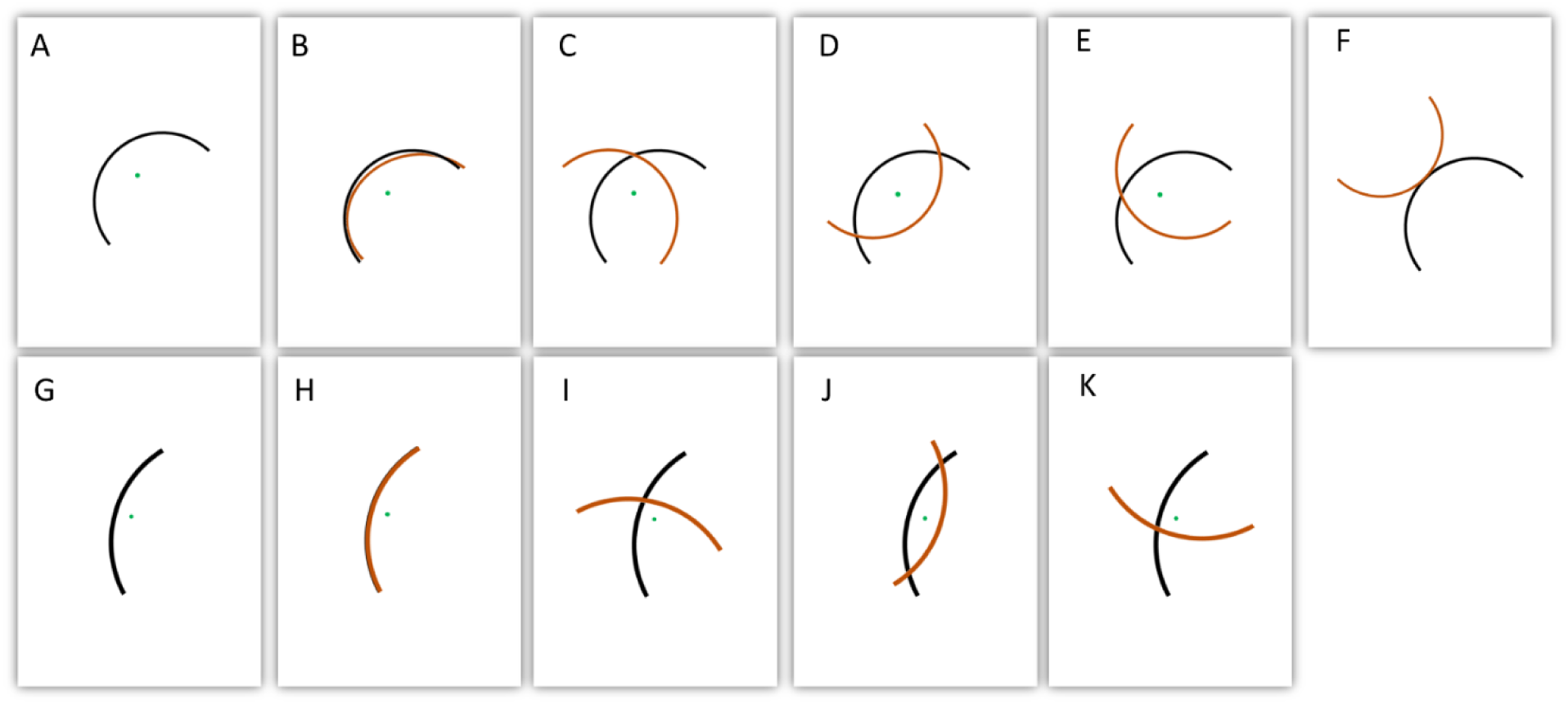
A – F. Illustration of a semicircle stimulus used in Experiment 5.10 (A), of its’ centroid (green dot), which was not shown during the experiment and of Davida’s different types of responses when asked to trace the depicted shape with ink (B-F, in red ink). G – K. Illustration of an arc stimulus used in Experiment 5.10(G), and of its’ centroid (green dot), which was not shown during the experiment, and of Davida’s different types of responses when asked to trace the depicted shape with ink (H-K, in red ink). A-K. These shapes’ centroid is illustrated by a green dot. This green dot was not depicted in the experiment.

##### 5.11. The structure of the representational frame: semi-circles and arcs (2)

We conducted an additional experiment in order to confirm the nature of the errors reported by Davida in Experiment 5.10 with a new type of measure. In each trial of this experiment Davida was presented with a semi-circular arrow (diameter: 415 pixels; 6.7 degrees of visual angle; see Figure S37) for as long as needed and asked to use the computer mouse to move a small round cursor and click as precisely as possible on the tip of the arrow. The arrow was displayed 5 times in each of 12 different orientations: 25 (Figure S37 A), 55, 85, 115, 145, 175, 205, 235, 265, 295, 325 and 355 degrees clockwise from the vertical. Davida clicked at less than 1 degree of visual angle (62 pixels) from the tip of the arrow in 3.3.% of the trials. In most of the other trials she made 7 types of errors, consisting in clicking at less than 1 degree of visual angle from the place where the tip of the arrow is depicted in red in Figure S37 B-H. In 5 last trials she clicked at a distance of more than 1 degree of visual angle from any interpretable landmark. These 7 error types confirmed that this type of shape is represented with respect to a representation frame composed of their axis of symmetry and a perpendicular axis intersecting the axis of symmetry at the level of the shape’s centroid (see main text and Figure 6 for more detail on these error types).

**Fig. S37.**
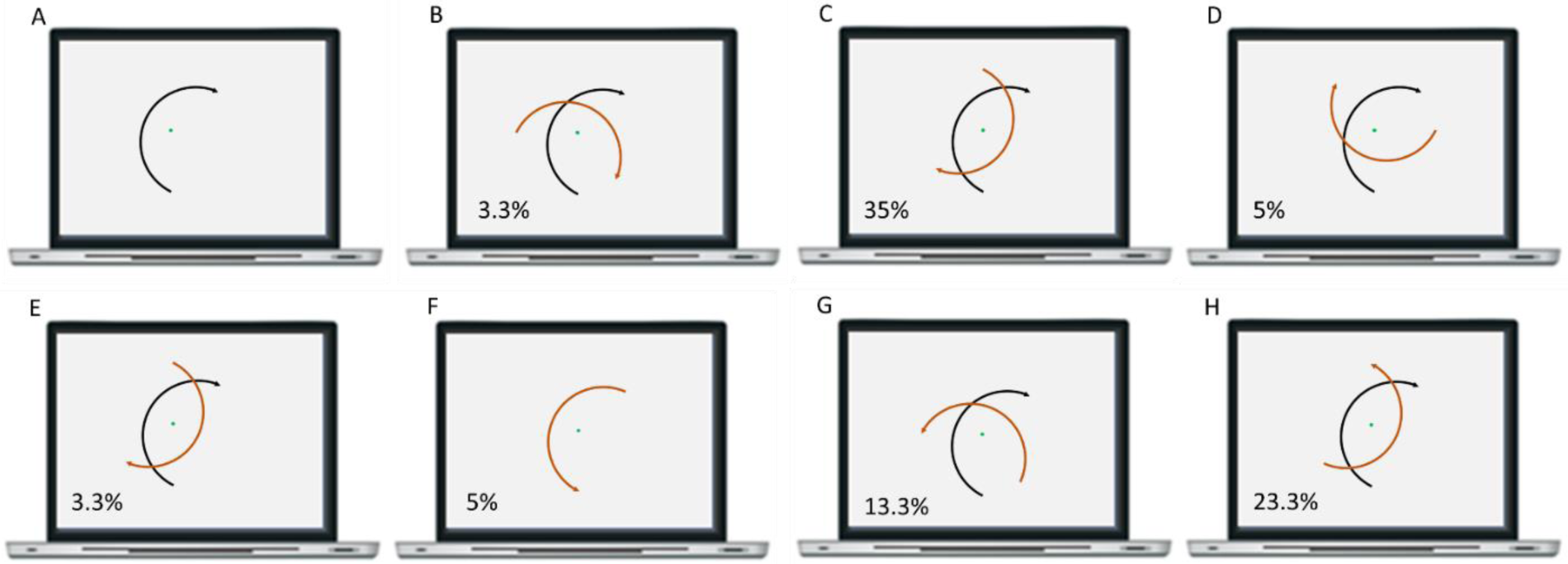
A. Illustration of a semicircular arrow stimulus used in Experiment 5.11 and of its centroid (green dot). The green dot was not presented during the experiment. B-H. The tip of semicircular arrows in red ink represent the 7 places in which Davida typically reported perceiving the tip of the depicted stimulus (the arrow shown here in black ink). The percentages correspond to the proportion of trials in which Davida clicked at less than one degree of visual angle from the place where the tip of the arrow in red ink is depicted in each figure (see main text and Figure 6 for more detail on these error types).

#### Materials and methods: set of results §6

##### 6.1. Naming the orientation of arrows displayed on an isoluminant background

Davida was shown 24 arrows pointing up, down, left or right and asked to indicate the orientation of these arrows by naming it (right, left, up, down). Stimuli were displayed one at the time, at the center of the screen, as long as needed by Davida, and consisted in large (8 x 3.5 degrees of visual angle), blue, red or green, arrows on an isoluminant (20 cd/m2, measured by a Konica Minolta LS-100) background of a different color (blue, red or green). Davida named the orientation of 1/24 stimuli accurately. Her errors consisted in responding as if the arrow were rotated by 90 (6/24), 180 (9/24) or 270 (8/24) degrees.

##### 6.2. Naming the letters p, b, d and q displayed on an isoluminant background

Davida was presented 6 times with the letters b, d, p and q and was asked to name them. Stimuli were displayed one at the time, at the center of the screen, as long as needed by Davida, and were composed of large (± 3.5 degrees of vertical visual angle) lower-case blue, red or green letters drawn in the Calibri font on an isoluminant (20 cd/m2, measured by a Konica Minolta LS-100) background of a different color (blue, red or green). She almost systematically named the letters as if they were inverted or rotated (see Table S4).

**Table S4.** Davida’s responses in the isoluminant letters reading task

| Targets | Davida's responses |  |  |  |
| --- | --- | --- | --- | --- |
|  | p | b | d | q |
| p | 1 | 1 | 4 |  |
| b | 2 |  | 4 |  |
| d | 6 |  |  |  |
| q | 2 | 1 | 3 |  |

##### 6.3. Arrow orientation judgment task with various luminance contrast

In a first experiment, Davida was shown arrows pointing up, down, left or right (randomly) and asked to indicate the orientation of these arrows by pressing on the corresponding key on a computer keyboard. These arrows were large (4 degrees of visual angle), of 7 different shades of grey (RGB of 0, 40, 93, 148, 202, 228 or 242; corresponding to 5.8, 9, 35, 88, 170, 221 and 244 cd/m2, measured by a Konica Minolta LS-100) and displayed one at the time, for as long as needed by Davida, at the center of the screen on white background (RGB of 255; 270 cd/m2).

Thus, they were 7 levels of luminance contrast (Background/Figure: 46.5, 30, 7.71, 3.07, 1.59, 1.22, 1.11). This experiment was carried out twice: The first session contained 140 stimuli (7 contrasts x 4 orientations x 5 repetitions). The second session was terminated at Davida’s request after 91 stimuli. As a result, Davida responded to 32, 33 or 34 stimuli displayed at each contrast level. As shown in Figure S38 A, Davida’s performance was influenced by the luminance contrast of the stimuli, varying from 0% correct responses at the two highest levels of luminance contrast to 84.3% correct responses at the lowest level.

In a second experiment, Davida was shown arrows pointing up, down, left or right (randomly) and asked to indicate the orientation of these arrows by pressing on the corresponding arrow key on the computer keyboard. These arrows were large (11.5 degrees of visual angle), of 10 different shades of light grey (178, 180, 182, 184, 186, 188, 190, 192, 194 and 196 cd/m2, measured by a Konica Minolta LS-100) and displayed one at the time, as long as needed by Davida, at the center of the screen on a light grey background (198 cd/m2). Thus, there were 10 different levels of luminance contrast (Background/Figure: 1.11, 1.10, 1.09, 1.08. 1.07, 1.06, 1.05, 1.04, 1.03, 1.02, 1.01). This experiment included 200 stimuli: 10 contrasts x 4 orientations x 5 repetition. As shown in Figure S38 B, Davida’s performance was influenced by the luminance contrast of the stimuli, varying from 70% of correct responses at the highest levels of contrast (1.11) to 100% correct responses at the three lowest level (1.03 and lower).

**Fig. S38.**
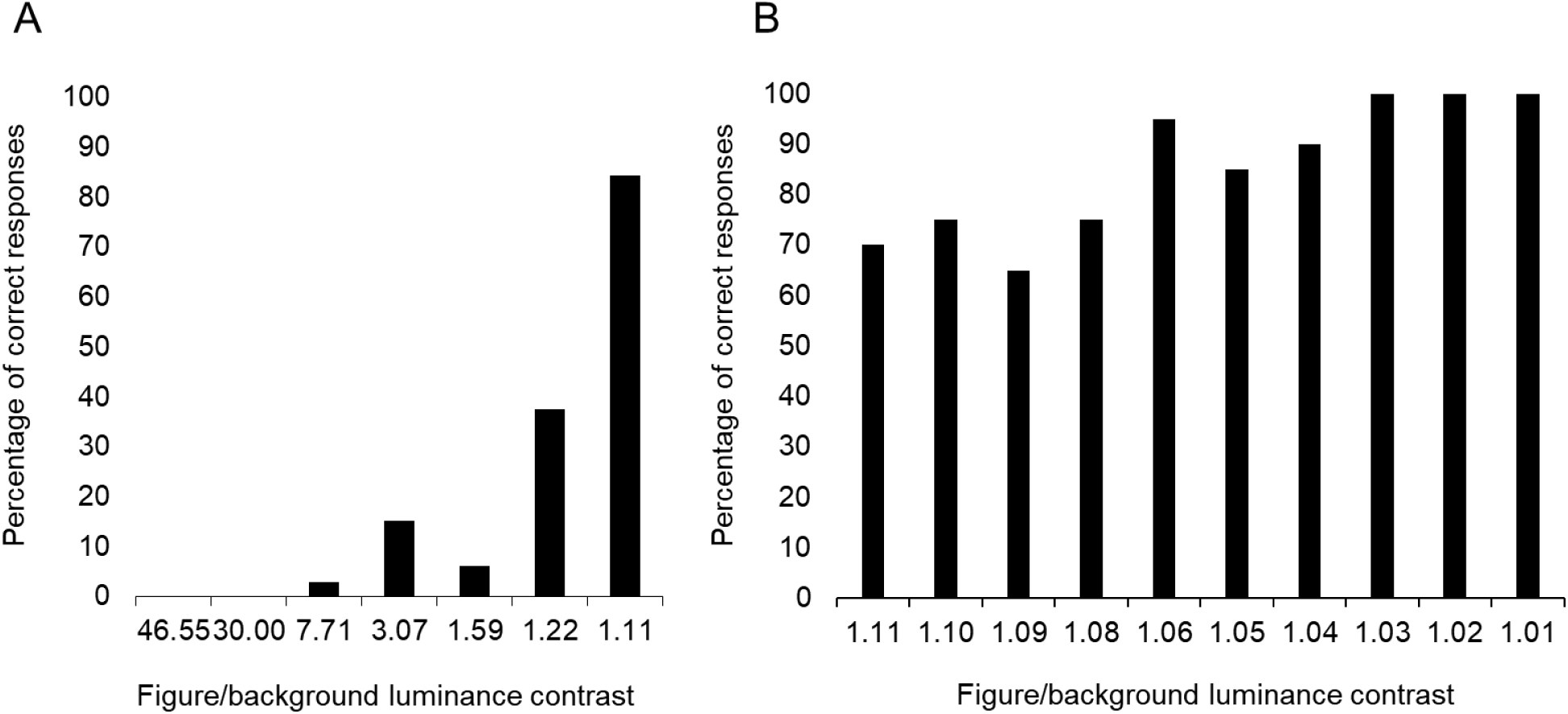
Davida’s percentage of correct responses in Experiment 6.3 for arrows of different levels of luminance contrast with the background.

##### 6.4. Pointing to the tip of a high or low-contrast arrow

In this task, a large (±10 x 6 degrees of visual angle) arrow pointing left, right, up or down was displayed at the center of the computer screen on white background (RBG 255) for an unlimited duration. In each trial, Davida was asked to use the computer mouse to move a small round cursor and click as precisely as possible on the tip of the arrow. In 50 trials, the arrow was black (RGB 0). In 50 additional trials, the arrow was colored in very light gray (RGB 253). The Movie S18, illustrates Davida’s performance in this task. Davida clicked approximately on the tip of the arrow (less than 50 pixels = 8 mm) in 100% of the trials when the arrow and the background had a low luminance contrast (light grey arrow), but only 2% of the trials when the arrow and the background had a high luminance contrast (black arrow). In the later condition, she almost systematically mislocated the position of the tip to approximately the place it would have been if the arrow were rotated by 90 degrees (34%), 180 degrees (24%) or 270 degrees (40%).

##### 6.5. Reading high and low contrast letters

In two separate sessions, Davida was asked to read 30 times the letters “b”, “p”, “d” and “q” displayed at the center of the computer screen on white background (RBG 255) for an unlimited duration. The letters were displayed in the Calibri font with a size of 166 (+- 3.7 x 2 degrees of visual angle) and were either colored in black (RGB 0), light grey (RGB 250) or very light grey (RGB 253). The Movie S19, illustrates Davida’s performance in this type of experiment. Davida made systematic errors when naming the letters displayed in black (0/40), in light grey (0/40) but read without any error or difficulty the letters displayed in very light grey (40/40).

##### 6.6. Low contrast objects orientation decision task

Davida was shown 40 line-drawings of objects from the Snodgrass and Vanderwart (60) set displayed once in their typical upright orientation and once upside-down. The line drawings were depicted in very light grey on white background to decrease their luminance contrast. Davida was presented with each stimulus one at a time, for as long as needed, and asked to decide whether the object was in its typical or atypical orientation. She performed the task perfectly (100% correct responses).

##### 6.7. Ponzo illusion task with high and low contrast stimuli

In this task, Davida was shown the same Ponzo illusion display and experiment as above (Experiment 1.12) but also, in addition, with 20 additional trials in which the display was shown at a very low luminance contrast with the background. When the display was displayed in black, Davida performed like in the first experiment with the same stimuli: she did not show the typical visual illusion (t (19) = −1.07, p = 0.3), drawing the lower line on average 6.1 pixels shorter (SD = 25.5) than the upper (reference) line, and was significantly better than the controls at matching the length of the two lines (modified t test: t (13) = −2.08, p = 0.05, see Figure S39). However, when the display was shown at very low luminance contrast with the background her performance became similar to that of the control participants: she showed the typical effect of the illusion (t (19) = 1.88, p = 0.07), drawing the drawing the lower line on average 12 pixels longer (SD = 30.4) than the upper (reference) line, and became significantly less accurate at matching the length of the two lines (paired t test (19) = 2.14, p = 0.046).

**Fig. S39.**
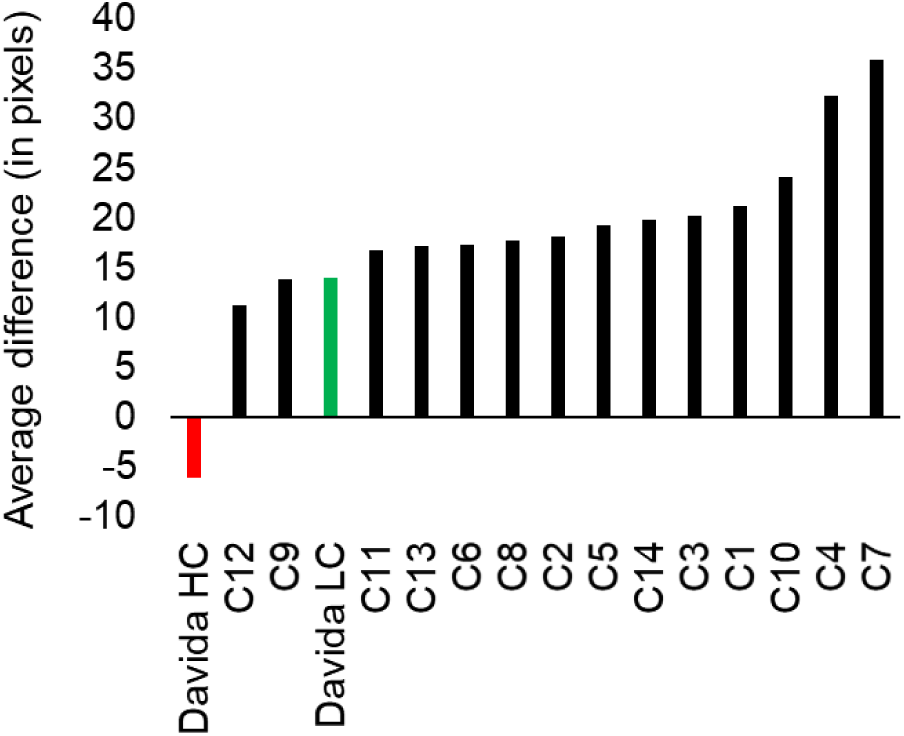
Control participants’ (C1 – C14, in black) and Davida’s average difference between the length of the two horizontal lines in the Ponzo illusion task (in pixels) in the high contrast (HC, in red) and low contrast (LC, in green) conditions. Individual data are aligned in ascending order on the horizontal axis as a function of the size of the difference between the length of the two lines (average length of the lower line – length of the upper reference line).

##### 6.8. Copying a tilted asymmetrical shape displayed at 6 levels of luminance contrast

In each trial of this task, Davida was randomly presented with one of two asymmetrical shapes (11.5 x 3.5 of visual angle, see Figure S29 A, B) displayed on white background (RGB 255) tilted from upright toward one of 4 possible orientations (45, 135, 225, 315) and was asked to copy it as precisely as possible on a separate sheet of paper. In a total of 288 trials, the stimuli were displayed 48 times (2 stimuli x 4 orientations x 8 repetitions) at 6 levels of grey (RGB of 0, 218, 230, 240, 245 and 252), resulting in 6 levels of luminance contrast with the white background. As shown in Figure S40, Davida made many errors when the stimuli were displayed with a high level of luminance contrast, but her error rate decreased when the luminance contrast between the shape and the background decreased. The Movie S20, illustrates Davida’s performance in this type of experiment with high and low levels of luminance contrast.

**Fig. S40.**
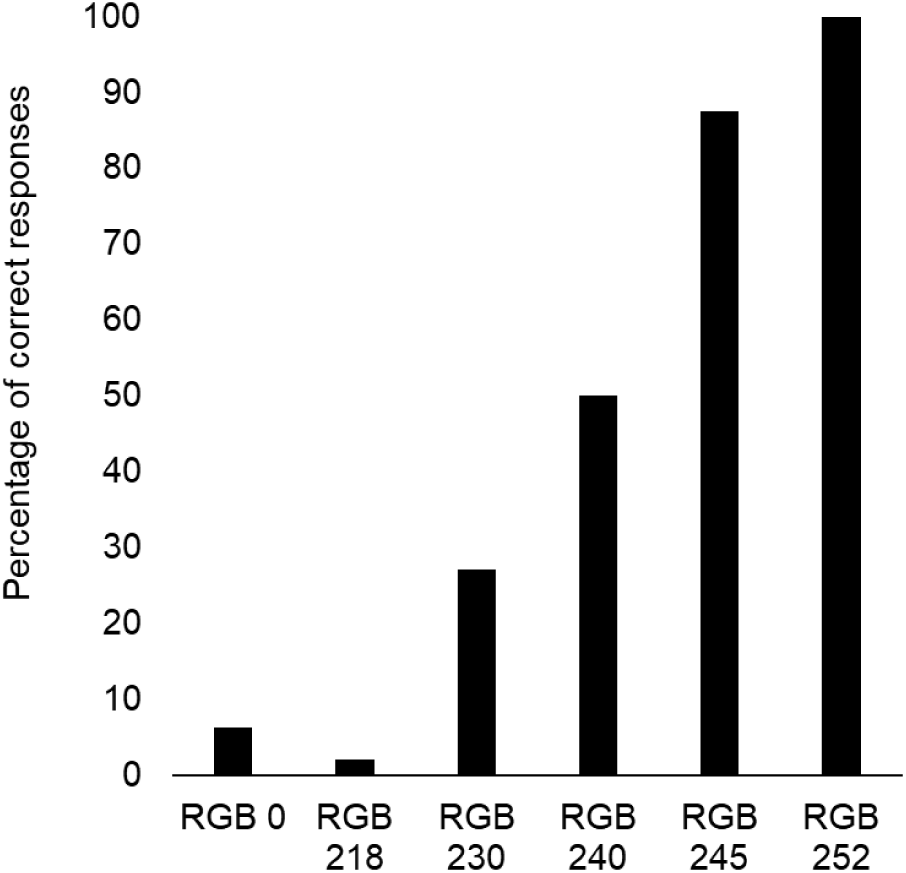
Davida’s percentage of correct responses for the different conditions of Experiment 6.8.

##### 6.9. Drawing all that she sees from a tilted asymmetrical shape displayed at 5 levels of luminance contrast

In each trial of this task, Davida was randomly presented with one of two asymmetrical shapes (11.5 x 3.5 of visual angle, see Figure S29 A, B) displayed on white background (RGB 255) tilted from upright toward one of 4 possible orientations (45, 135, 225, 315) for 2 seconds and asked, after each presentation, to draw all she saw. In a total of 80 trials, the stimuli were displayed 16 times (2 shapes x 4 orientations x 2 repetitions) at 5 levels of grey (RGB of 0, 230, 240, 245 and 252), resulting in 5 levels of luminance contrast with the white background.

Figure S41 A shows the average number of orientations drawn for each stimulus in all the conditions. As one can see from this figure, Davida perceived numerous different orientations of stimuli displayed with a high level of luminance contrast, but this number decreased when the luminance contrast between the shape and the background decreased and she reported seeing only one orientation of each stimulus shown at the lowest luminance contrast. Figure S41 B shows the percentage of trials in which Davida’s response included the correct orientation of the stimulus. As one can see from this figure, Davida’s response often included the correct orientation at all contrast levels. Nevertheless, the percentage of trials that included the correct response increased with the decrease of the luminance contrast.

**Fig. S41.**
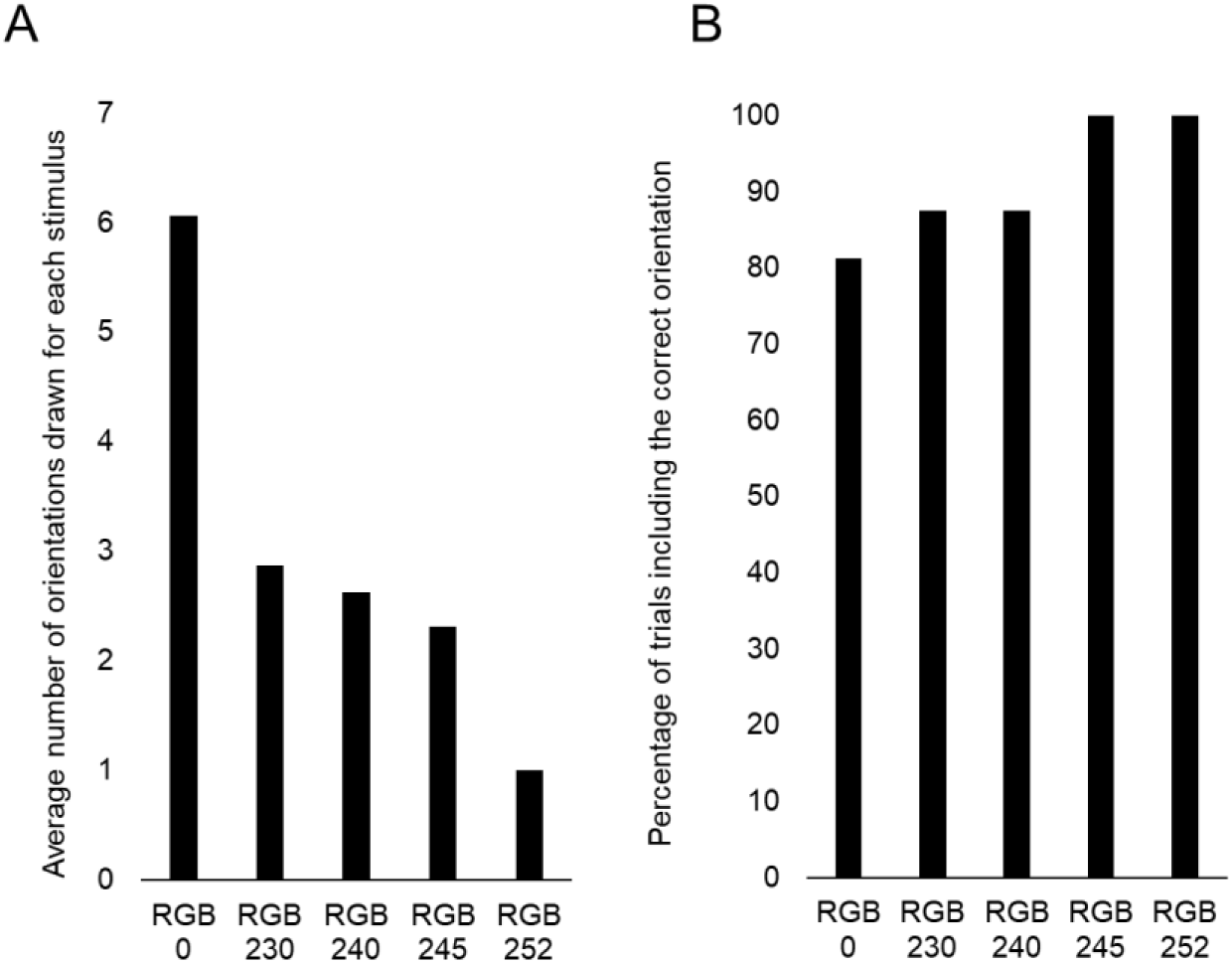
A. Average number of orientations drawn per stimulus in the five levels of luminance contrast. B. Percentage of trials in which Davida’s response included the correct orientation in the five levels of luminance contrast.

##### 6.10. Davida’s perception of shapes from motion

We tested Davida’s perception of arrows and letters in a shape from motion experiments. In these experiments, the shape and the background were composed of white (1/6), black (1/6) and grey (2/3) pixels and the shapes were visible only because the motion (60 pixels/second) of the white and black dots within the shape region were in a different direction (right) from that of their motion on the background (left). In the first experiment Davida saw 20 large arrows in one of four different orientations (left, right, up, down) and had to report verbally their orientation. In the second task, she saw 20 letters displayed one at a time (b, p, d, q) and had to name each of them. Davida was flawless in both tasks.

##### 6.11. Naming the orientation of high and low spatial frequency arrows

Davida was randomly shown arrows pointing up, down, left or right and asked to indicate the orientation of these arrows by pressing on the corresponding arrow key on the computer keyboard. These arrows were large (±10 x 6 degrees of visual angle), of 3 different levels of gaussian blur (0, 50, 80) and displayed one at the time, at the center of the screen on white background for as long as needed (see Figure S42). In total, there were 20 stimuli in each condition (5 of each orientation). Davida’s performance was 0% correct responses at the two lowest levels of blur and 100% of correct responses at the highest level.

**Fig. S42.**
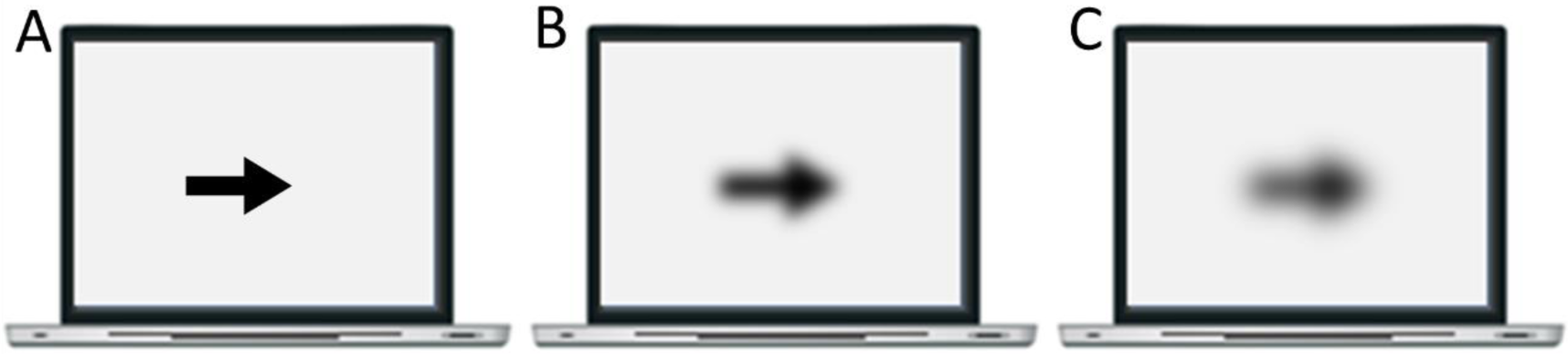
Illustration of high (A), medium (B) and low (C) spatial frequency stimuli used in Experiment 6.11-6.12.

##### 6.12. Pointing to the tip of high and low spatial frequency arrow

In this task, a large (±10 x 6 degrees of visual angle) arrow pointing left, right, up or down was displayed at the center of the computer screen on white background (RBG 255) for an unlimited duration. On each trial, Davida was asked to use the computer mouse to move a small round cursor and click as precisely as possible on the tip of the arrow. In 20 trials, the arrow had a high spatial frequency (See Figure S42 A). In 20 other trials, the arrow was edited with a gaussian blur with a radius of 80 pixels (See Figure S42 C). The Movie S21 illustrates Davida’s performance in this type of experiment. Davida systematically mislocated the position of the tip of the high spatial frequency arrows to approximately the place where it would have been if the arrow were rotated by 90 degrees (40%), 180 degrees (20%) or 270 degrees (40%). When the arrow was blurred, however, she correctly localized the tip of the arrow (less than 50 pixels) in 100% of the trials.

##### 6.13. Reading high and low spatial frequency letters

In this experiment, Davida was asked to read the letters “b”, “p”, “d” and “q” displayed at the center of the computer screen on white background (RBG 255) for an unlimited duration. The letters were displayed in black, in the Calibri font with a size of +- 3.7 x 2 degrees of visual angle (166) and were either not further edited or edited with a gaussian blur of 50 or 80 pixels (See Figure S43). One session comprised 20 trials in each condition (60 stimuli). Davida participated in 2 sessions. She named 2 letters accurately when the letters were displayed in black (2/40), 0 when reading the letter edited with a gaussian blur with a radius of 50 pixels (0/40), but she read all the letters accurately when the letter was edited with a gaussian blur with a radius of 80 pixels (40/40). The Movie S22 illustrates Davida’s performance in this type of experiment.

**Fig. S43.**
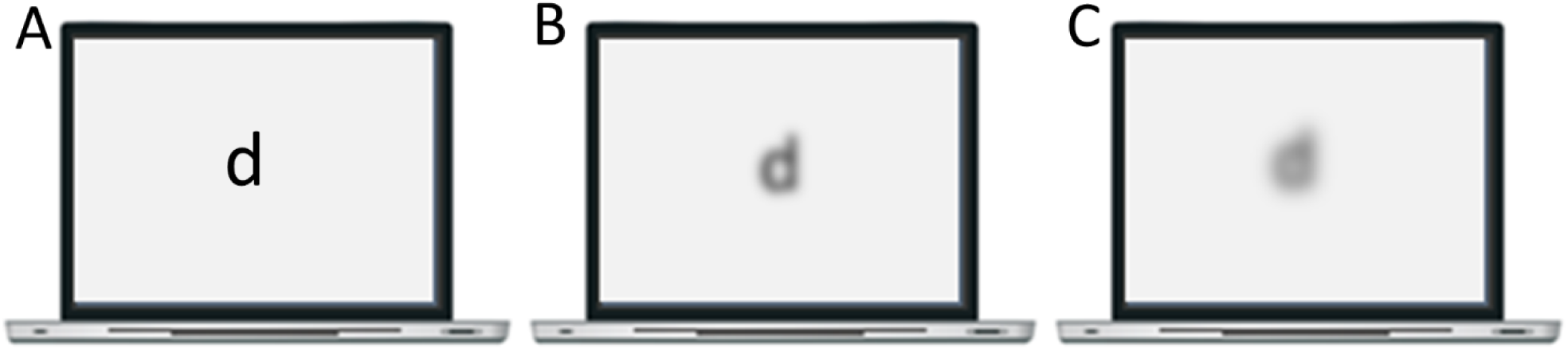
Illustration of high (A), medium (B) and low (C) spatial frequency stimuli used in Experiment 6.13.

##### 6.14. Copying a tilted asymmetrical shape displayed at 6 levels of spatial frequency

In each trial of this task, Davida was randomly presented with one of two asymmetrical shapes (11.5 x 3.5 of visual angle, see Figure S29 A, B) displayed on white background (RGB 255) tilted from upright toward one of 4 possible orientations (45, 135, 225, 315) and was asked to copy it as precisely as possible on a separate sheet of paper. In a total of 400 trials, the stimuli were displayed 88 times (2 stimuli x 4 orientations x 11 repetitions) at each of four levels of gaussian blur (radius of 0, 20, 40 or 60) and 24 times at each of two levels of gaussian blur (2 stimuli x 4 orientations x 3 repetitions). As shown in Figure S44, Davida made many errors when the stimuli were displayed at a high level of spatial frequency (lower levels of gaussian blur), but her error rate decreased when the spatial frequency decreased (the gaussian blur increased) and she eventually became flawless at the lowest levels of spatial frequency (highest levels of blur). The Movie S23, illustrates Davida’s performance in this type of experiment with arrows presented at a high and low level of spatial frequency.

**Fig. S44.**
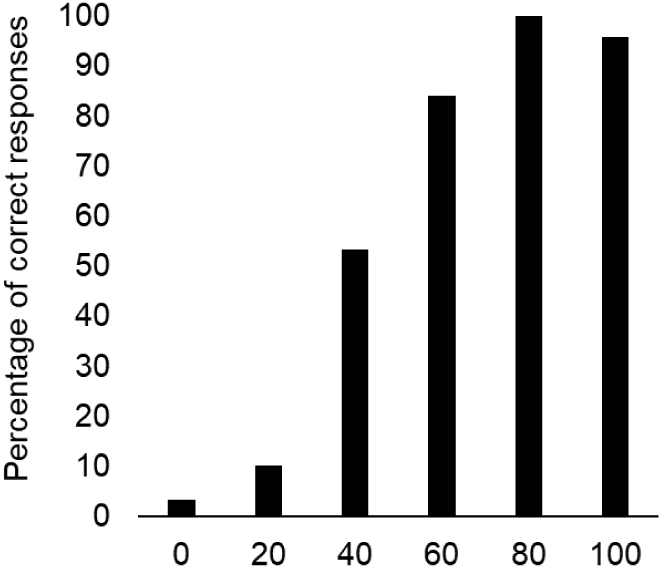
Davida’s percentage correct responses by level of gaussian blur (from 0 to 100).

##### 6.15. Judging the orientation of 3D stimuli defined by binocular and monocular cues

In one experiment, Davida was shown 20 trials in which a 3D wooden black arrow was positioned in front of her on white sheet of paper in one of four possible orientations (right, left, up, down) and asked to tell the orientation (right, left, up, down). In a second experiment, she was presented with 20 trials in which a 3D “b” shaped wooden letter was positioned in front of her on white sheet of paper in one of four possible orientations (b, d, p, q) and asked to name the letter (b, d, p, q). Davida performed both tasks easily and flawlessly. Then, she was asked to perform the same two experiments monocularly for both eyes. She performed the two tasks flawlessly with either eyes.

**Movie S1 (separate file).** Part I: Davida, interviewed by G.V., describes what she perceives when shown an arrow on the computer screen. Part II: Davida, interviewed by G.V., describes what she perceives when shown an abstract shape on the computer screen.

**Movie S2 (separate file).** Davida copies arrows.

**Movie S3 (separate file).** Davida copies letters.

**Movie S4 (separate file).** Davida reads orientation-sensitive letters.

**Movie S5 (separate file).** Davida uses the computer mouse to move a small round cursor and click as precisely as possible on the tip of the arrow. Part I: The arrows are pointing left, right, up or down. Part II: The arrows are tilted.

**Movie S6 (separate file).** Davida places her right thumb and index finger on the extremities of a series of black lines, black lines ending with a dot at each extremity or two dots displayed on a sheet of paper hanged at a comfortable distance in front of her. Part I: The importance of accuracy was stressed. Part II: The importance of speed was stressed.

**Movie S7 (separate file).** Davida is standing in front of a sheet of paper with her eyes closed, and, at the experimenter’s signal, she opens her eyes and places her index finger as fast as possible where she sees the tip of a large black arrow.

**Movie S8 (separate file).** Davida, blindfolded, is asked to decide which of four possible letters (b, d, p, q) was traced on the back of her hand. She performed this task easily and perfectly.

**Movie S9 (separate file).** Part I: Davida uses the computer mouse to move a small round cursor and click as precisely as possible on the tip of the arrow. In half of the trials the arrow is fully depicted. In the other half, the center of the arrow is hidden behind a mask. Part II: Davida sees two red or two black rectangles separated from each other by three pixels or connected by a 4-pixels line and uses the computer mouse to move a small round cursor and click as precisely as possible on the little “indent” at the extremity of the rectangle.

**Movie S10 (separate file).** Davida uses the computer mouse to move a small round cursor and click as precisely as possible on the tip of a bicolor arrow. In half of the trials the blending of the colors occurred over a short area. In the other half, the blending of the colors occurred over a large area.

**Movie S11 (separate file).** Davida sees an arrow implied by a series of unconnected small dots within an arrow composed of connected solid black straight lines and uses the computer mouse to move a small round cursor and click as precisely as possible on either the dot placed at the tip of the dotted arrow (Part I) or at the tip of the arrow made of solid black lines (part II).

**Movie S12 (separate file).** Davida sees one of three types of large black arrow for 200 ms (made of solid lines, of small or large unconnected dots) and uses the computer mouse to move a small round cursor and click as precisely as possible on the tip of these arrows.

**Movie S13 (separate file).** Davida sees a tilted asymmetrical shape and copies it as precisely as possible on a separate sheet of paper.

**Movie S14 (separate file).** Davida sees a tilted asymmetrical shape and copies it as precisely as possible on a separate sheet of paper.

**Movie S15 (separate file).** Davida sees a tilted asymmetrical shape for 2 seconds and then draws all the orientations of that shape that she has seen on a separate sheet of paper.

**Movie S16 (separate file).** Davida is given a drawing of an asymmetrical shape printed on a sheet of paper and is asked to overlap the shape with ink.

**Movie S17 (separate file).** Davida is shown a large black circle with a small semicircular indent and uses the computer mouse to move a small round cursor and click as precisely as possible on the places “where she sees the indent” before pressing the space bar to launch the next trial. She was free to click any number of times on the stimulus.

**Movie S18 (separate file).** Davida uses the computer mouse to move a small round cursor and click as precisely as possible on the tip of black (RGB 0) and light grey (RGB 253) arrows presented on white background (RGB 255).

**Movie S19 (separate file).** Davida reads black (RGB 0) and very light grey (RGB 253) letters (“b”, “p”, “d” and “q”) displayed on white background (RBG 255).

**Movie S20 (separate file).** Davida copies a black (RBG 0) or a very light grey (RGB 253) tilted asymmetrical shape on a separate sheet of paper.

**Movie S21 (separate file).** Davida uses the computer mouse to move a small round cursor and click as precisely as possible on the tip of high spatial frequency and blurred arrows.

**Movie S22 (separate file).** Davida reads high spatial frequency and blurred letters (“b”, “p”, “d” and “q).

**Movie S23 (separate file).** Davida copies high spatial frequency and blurred asymmetrical shapes as precisely as possible on a separate sheet of paper.

## References

Blake, R. (2001). A primer on binocular rivalry, including current controversies. Brain and Mind, 2(1), 5–38. https://doi.org/10.1023/A:1017925416289

Bryant, B. R., Wiederholt, J. L., & Bryant, D. P. (1991). Gray Diagnostic Reading Tests–Second Edition (GRDT-2). Austin, TX: Pro-Ed.

Burr, D. C., & Morrone, M. C. (2011). Spatiotopic coding and remapping in humans. Philosophical Transactions of the Royal Society B: Biological Sciences. https://doi.org/10.1098/rstb.2010.0244

Bushnell, B. N., Harding, P. J., Kosai, Y., Bair, W., & Pasupathy, A. (2011). Equiluminance cells in visual cortical area V4. Journal of Neuroscience. https://doi.org/10.1523/JNEUROSCI.1890-11.2011

Cadieu, C., Kouh, M., Pasupathy, A., Connor, C. E., Riesenhuber, M., & Poggio, T. (2007). A model of V4 shape selectivity and invariance. Journal of Neurophysiology, 98(3), 1733– 1750. https://doi.org/10.1152/jn.01265.2006

Caramazza, A., & Hillis, A. E. (1990). Levels of representation, co-ordinate frames, and unilateral neglect. Cognitive Neuropsychology, 7(5–6), 391–445.

Chaisilprungraung, T., German, J., & McCloskey, M. (2019). How are object shape axes defined? Evidence from mirror-image confusions. Journal of Experimental Psychology: Human Perception and Performance, 45(1), 111.

Clifford, C. W. G., Spehar, B., Solomon, S. G., Martin, P. R., & Zaidi, Q. (2003). Interactions between color and luminance in the perception of orientation. Journal of Vision. https://doi.org/10.1167/3.2.1

Colby, C. L. (1998). Action-Oriented Spatial Reference Frames in Cortex. Neuron, 20(1), 15–24. https://doi.org/10.1016/S0896-6273(00)80429-8

Conners, C. K. (2010). Conners comprehensive behavior rating scales (Conners CBRS). Multi-Health Systems.

Connor, C. E., & Knierim, J. J. (2017). Integration of objects and space in perception and memory. Nature Neuroscience, 20(11), 1493–1503. https://doi.org/10.1038/nn.4657

Cooper, A. C. G., & Humphreys, G. W. (2000). Task-specific effects of orientation information: Neuropsychological evidence. Neuropsychologia, 38(12), 1607–1615. https://doi.org/10.1016/S0028-3932(00)00070-1

Cowey, A., & Gross, C. G. (1970). Effects of foveal prestriate and inferotemporal lesions on visual discrimination by rhesus monkeys. Experimental Brain Research, 11(2), 128–144. https://doi.org/10.1007/BF00234318

Crawford, J. R., & Garthwaite, P. H. (2005). Testing for suspected impairments and dissociations in single-case studies in neuropsychology: evaluation of alternatives using monte carlo simulations and revised tests for dissociations. Neuropsychology, 19(3), 318–331. https://doi.org/10.1037/0894-4105.19.3.318

Crawford, J. R., & Howell, D. C. (1998). Comparing an individual’s test score against norms derived from small samples. The Clinical Neuropsychologist, 12(4), 482–486.

Davidoff, J., & Warrington, E. K. (1999). The bare bones of object recognition: Implications from a case of object recognition impairment. Neuropsychologia, 37(3), 279–292. https://doi.org/10.1016/S0028-3932(98)00076-1

Davidoff, Jules, & Warrington, E. K. (2001). A particular difficulty in discriminating between mirror images. Neuropsychologia, 39(10), 1022–1036. https://doi.org/10.1016/S0028-3932(01)00039-2

De Valois, R. L., Albrecht, D. G., & Thorell, L. G. (1982). Spatial frequency selectivity of cells in macaque visual cortex. Vision Research, 22(5), 545–559. https://doi.org/10.1016/0042-6989(82)90113-4

Delis, D. C., Kaplan, E., & Kramer, J. H. (2001). Delis-Kaplan Executive Function System®(D-KEFS®): Examiner’s Manual: Flexibility of Thinking, Concept Formation, Problem Solving, Planning, Creativity, Impluse Control, Inhibition. Pearson.

DiCarlo, J. J., Zoccolan, D., & Rust, N. C. (2012). How Does the Brain Solve Visual Object Recognition? Neuron, 73(3), 415–434. https://doi.org/10.1016/j.neuron.2012.01.010

Driver, J., Baylis, G. C., Goodrich, S. J., & Rafal, R. D. (1994). Axis-based neglect of visual shapes. Neuropsychologia, 32(11), 1353–1356. https://doi.org/10.1016/0028-3932(94)00068-9

Driver, J., & Pouget, A. (2000). Object-Centered Visual Neglect, or Relative Egocentric Neglect? Journal of Cognitive Neuroscience, 12(3), 542–545. https://doi.org/10.1162/089892900562192

Duhamel, Colby, C., & Goldberg, M. (1992). The updating of the representation of visual space in parietal cortex by intended eye movements. Science, 255(5040), 90–92. https://doi.org/10.1126/science.1553535

Eacott, M. J., & Gaffan, D. (1991). The role of monkey inferior parietal cortex in visual discrimination of identity and orientation of shapes. Behavioural Brain Research, 46(1), 95–98. https://doi.org/10.1016/S0166-4328(05)80100-7

El-Shamayleh, Y., & Pasupathy, A. (2016). Contour curvature as an invariant code for objects in visual area V4. Journal of Neuroscience, 36(20), 5532–5543. https://doi.org/10.1523/JNEUROSCI.4139-15.2016

Felleman, D. J., & Van Essen, D. C. (1991). Distributed Hierarchical Processing in the Primate Cerebral Cortex. Cerebral Cortex, 1(1), 1–47. https://doi.org/10.1093/cercor/1.1.1

Ferrera, V., Nealey, T., & Maunsell, J. (1994). Responses in macaque visual area V4 following inactivation of the parvocellular and magnocellular LGN pathways. The Journal of Neuroscience, 14(4), 2080–2088. https://doi.org/10.1523/JNEUROSCI.14-04-02080.1994

Flanagan, P., Cavanagh, P., & Favreau, O. E. (1990). Independent orientation-selective mechanisms for the cardinal directions of colour space. Vision Research. https://doi.org/10.1016/0042-6989(90)90102-Q

Gainotti, G., Messerli, P., & Tissot, R. (1972). Qualitative analysis of unilateral spatial neglect in relation to laterality of cerebral lesions. Journal of Neurology, Neurosurgery & Psychiatry, 35(4), 545–550. https://doi.org/10.1136/jnnp.35.4.545

Gallant, J. L., Braun, J., & Van Essen, D. (1993). Selectivity for polar, hyperbolic, and Cartesian gratings in macaque visual cortex. Science, 259(5091), 100–103. https://doi.org/10.1126/science.8418487

Gallant, J. L., Connor, C. E., Rakshit, S., Lewis, J. W., & Van Essen, D. C. (1996). Neural responses to polar, hyperbolic, and Cartesian gratings in area V4 of the macaque monkey. Journal of Neurophysiology, 76(4), 2718–2739. https://doi.org/10.1152/jn.1996.76.4.2718

Gioia, G. A., Isquith, P. K., Guy, S. C., & Kenworthy, L. (2000). BRIEF: Behavior rating inventory of executive function. Psychological Assessment Resources.

Goodale, M. A., Milner, A. D., Jakobson, L. S., & Carey, D. P. (1991). A neurological dissociation between perceiving objects and grasping them. Nature, 349(6305), 154–156. https://doi.org/10.1038/349154a0

Gregory, E., & McCloskey, M. (2010). Mirror-image confusions: Implications for representation and processing of object orientation. Cognition, 116(1), 110–129. https://doi.org/10.1016/j.cognition.2010.04.005

Gross, C. G. (1978). Inferior temporal lesions do not impair discrimination of rotated patterns in monkeys. Journal of Comparative and Physiological Psychology. https://doi.org/10.1037/h0077515

Haralick, R. M., & Shapiro, L. G. (1991). Glossary of computer vision terms. Pattern Recognition, 24(1), 69–93.

Harris, I. M., Harris, J. A., & Caine, D. (2001). Object orientation agnosia: A failure to find the axis? Journal of Cognitive Neuroscience, 13(6), 800–812. https://doi.org/10.1162/08989290152541467

Holmes, E., & Gross, C. (1984). Effects of inferior temporal lesions on discrimination of stimuli differing in orientation. The Journal of Neuroscience, 4(12), 3063–3068. https://doi.org/10.1523/JNEUROSCI.04-12-03063.1984

Hommel, B. (1993). The relationship between stimulus processing and response selection in the Simon task: Evidence for a temporal overlap. Psychological Research, 55(4), 280–290.

Hubel, D. H., & Wiesel, T. N. (1962). Receptive fields, binocular interaction and functional architecture in the cat’s visual cortex. The Journal of Physiology, 160(1), 106–154. https://doi.org/10.1113/jphysiol.1962.sp006837

Ishihara, S. (1987). Test for colour-blindness. Kanehara Tokyo, Japan.

Karnath, H. O., Ferber, S., & Bülthoff, H. H. (2000). Neuronal representation of object orientation. Neuropsychologia, 38(9), 1235–1241. https://doi.org/10.1016/S0028-3932(00)00043-9

Kim, T., Bair, W., & Pasupathy, A. (2019). Neural Coding for Shape and Texture in Macaque Area V4. The Journal of Neuroscience, 39(24), 4760–4774. https://doi.org/10.1523/JNEUROSCI.3073-18.2019

Kolster, H., Peeters, R., & Orban, G. A. (2010). The retinotopic organization of the human middle temporal area MT/V5 and its cortical neighbors. Journal of Neuroscience. https://doi.org/10.1523/JNEUROSCI.2069-10.2010

Larsson, J., & Heeger, D. J. (2006). Two retinotopic visual areas in human lateral occipital cortex. Journal of Neuroscience. https://doi.org/10.1523/JNEUROSCI.1657-06.2006

Livingstone, M. S., & Hubel, D. H. (1987). Psychophysical evidence for separate channels for the perception of form, color, movement, and depth. Journal of Neuroscience, 7(11), 3416– 3468.

Logothetis, N. K., Pauls, J., & Poggio, T. (1995). Shape representation in the inferior temporal cortex of monkeys. Current Biology, 5(5), 552–563. https://doi.org/10.1016/S0960-9822(95)00108-4

Marr, D., & Nishihara, H. K. (1978). Representation and recognition of the spatial organization of three-dimensional shapes. Proceedings of the Royal Society of London. Series B. Biological Sciences, 200(1140), 269–294. https://doi.org/10.1098/rspb.1978.0020

Martinaud, O., Mirlink, N., Bioux, S., Bliaux, E., Champmartin, C., Pouliquen, D., … Gérardin, E. (2016). Mirrored and rotated stimuli are not the same: A neuropsychological and lesion mapping study. Cortex, 78, 100–114. https://doi.org/10.1016/j.cortex.2016.03.002

Martinaud, O., Mirlink, N., Bioux, S., Bliaux, E., Lebas, A., Gerardin, E., & Hannequin, D. (2014). Agnosia for Mirror Stimuli: A New Case Report with a Small Parietal Lesion. Archives of Clinical Neuropsychology, 29(7), 724–728. https://doi.org/10.1093/arclin/acu032

McCarney, D., & Greenberg, L. M. (1990). Test of Variables of Attention (TOVA). St. Paul: Attention Technology.

McCloskey, M. (2004). Spatial representations and multiple-visual-systems hypotheses: Evidence from a developmental deficit in visual location and orientation processing. Cortex. https://doi.org/10.1016/S0010-9452(08)70164-3

McCloskey, M. (2009). Visual Reflections. In Visual Reflections: A Perceptual Deficit and Its Implications. https://doi.org/10.1093/acprof:oso/9780195168693.001.0001

McCloskey, M., Rapp, B., Yantis, S., Rubin, G., Bacon, W. F., Dagnelie, G., … Palmer, E. (1995). A Developmental Deficit in Localizing Objects from Vision. Psychological Science, 6(2), 112–117. https://doi.org/10.1111/j.1467-9280.1995.tb00316.x

McCloskey, M., Valtonen, J., & Cohen Sherman, J. (2006). Representing orientation: A coordinate-system hypothesis and evidence from developmental deficits. Cognitive Neuropsychology, 23(5), 680–713. https://doi.org/10.1080/02643290500538356

McKyton, A., & Zohary, E. (2007). Beyond Retinotopic Mapping: The Spatial Representation of Objects in the Human Lateral Occipital Complex. Cerebral Cortex, 17(5), 1164–1172. https://doi.org/10.1093/cercor/bhl027

Melcher, D., & Morrone, M. C. (2015). Nonretinotopic visual processing in the brain. Visual Neuroscience, 32, E017. https://doi.org/10.1017/S095252381500019X

Merigan, W. H., & Maunsell, J. H. R. (1993). How parallel are the primate visual pathways? Annual Review of Neuroscience, 16(1), 369–402.

Milner, D., & Goodale, M. (2006). The Visual Brain in Action. In The Visual Brain in Action. https://doi.org/10.1093/acprof:oso/9780198524724.001.0001

Mozer, M. C. (2002). Frames of reference in unilateral neglect and visual perception: A computational perspective. Psychological Review, 109(1), 156–185. https://doi.org/10.1037/0033-295X.109.1.156

Nassi, J. J., & Callaway, E. M. (2009). Parallel processing strategies of the primate visual system. Nature Reviews Neuroscience, 10(5), 360.

Oleskiw, T. D., Nowack, A., & Pasupathy, A. (2018). Joint coding of shape and blur in area V4. Nature Communications. https://doi.org/10.1038/s41467-017-02438-8

Olson, C. R. (2003). Brain representation of object-centered space in monkeys and humans. Annual Review of Neuroscience, 26(1), 331–354. https://doi.org/10.1146/annurev.neuro.26.041002.131405

Palmer, S. E. (1985). The role of symmetry in shape perception. Acta Psychologica, 59(1), 67– 90. https://doi.org/10.1016/0001-6918(85)90042-3

Palmer, S., & Rock, I. (1994). Rethinking perceptual organization: The role of uniform connectedness. Psychonomic Bulletin & Review, 1(1), 29–55. https://doi.org/10.3758/BF03200760

Pasupathy, A. (2015). The neural basis of image segmentation in the primate brain. Neuroscience, 296, 101–109. https://doi.org/10.1016/j.neuroscience.2014.09.051

Pasupathy, A., & Connor, C. E. (2001). Shape Representation in Area V4: Position-Specific Tuning for Boundary Conformation. Journal of Neurophysiology, 86(5), 2505–2519. https://doi.org/10.1152/jn.2001.86.5.2505

Peirce, J. W. (2007). PsychoPy—psychophysics software in Python. Journal of Neuroscience Methods, 162(1–2), 8–13.

Peirce, J. W. (2009). Generating stimuli for neuroscience using PsychoPy. Frontiers in Neuroinformatics, 2, 10.

Peirce, J. W. (2015). Understanding mid-level representations in visual processing. Journal of Vision, 15(7), 5. https://doi.org/10.1167/15.7.5

Pelli, D. G., & Robson, J. G. (1988). The design of a new letter chart for measuring contrast sensitivity. Clinical Vision Sciences. Citeseer.

Pflugshaupt, T., Nyffeler, T., von Wartburg, R., Wurtz, P., Lüthi, M., Hubl, D., … Müri, R. M. (2007). When left becomes right and vice versa: Mirrored vision after cerebral hypoxia. Neuropsychologia, 45(9), 2078–2091. https://doi.org/10.1016/j.neuropsychologia.2007.01.018

Priftis, K., Rusconi, E., Umiltà, C., & Zorzi, M. (2003). Pure agnosia for mirror stimuli after right inferior parietal lesion. Brain, 126(4), 908–919. https://doi.org/10.1093/brain/awg075

Quinlan, P. T., & Humphreys, G. W. (1993). Perceptual Frames of Reference and Two-Dimensional Shape Recognition: Further Examination of Internal Axes. Perception, 22(11), 1343–1364. https://doi.org/10.1068/p221343

Riddock, M. J., Humphreys, G. W., Jacobson, S., Pluck, G., Bateman, A., & Edwards, M. (2004). Impaired orientation discrimination and localisation following parietal damage: On the interplay between dorsal and ventral processes in visual perception. Cognitive Neuropsychology, 21(6), 597–623. https://doi.org/10.1080/02643290342000230

Robinson, G., Cohen, H., & Goebel, A. (2011). A case of complex regional pain syndrome with agnosia for object orientation. Pain, 152(7), 1674–1681. https://doi.org/10.1016/j.pain.2011.02.010

Rock, I. (1973). Orientation and form. Academic Press.

Roe, A. W., Chelazzi, L., Connor, C. E., Conway, B. R., Fujita, I., Gallant, J. L., … Vanduffel, W. (2012). Toward a Unified Theory of Visual Area V4. Neuron, 74(1), 12–29. https://doi.org/10.1016/j.neuron.2012.03.011

Rollenhagen, J. E., & Olson, C. R. (2000). Mirror-image confusion in single neurons of the macaque inferotemporal cortex. Science, 287(5457), 1506–1508. https://doi.org/10.1126/science.287.5457.1506

Rust, N. C., & DiCarlo, J. J. (2010). Selectivity and Tolerance (“Invariance”) Both Increase as Visual Information Propagates from Cortical Area V4 to IT. Journal of Neuroscience, 30(39), 12978–12995. https://doi.org/10.1523/JNEUROSCI.0179-10.2010

Sandoval, J., & Echandia, A. (1994). Behavior assessment system for children. Journal of School Psychology, 32(4), 419–425. https://doi.org/10.1016/0022-4405(94)90037-X

Sebastian, S., Burge, J., & Geisler, W. S. (2015). Defocus blur discrimination in natural images with natural optics. Journal of Vision. https://doi.org/10.1167/15.5.16

Sekuler, A. B. (1996). Axis of elongation can determine reference frames for object perception. Canadian Journal of Experimental Psychology/Revue Canadienne de Psychologie Expérimentale, 50(3), 270–279. https://doi.org/10.1037/1196-1961.50.3.270

Sekuler, A. B., & Swimmer, M. B. (2000). Interactions between symmetry and elongation in determining reference frames for object perception. Canadian Journal of Experimental Psychology/Revue Canadienne de Psychologie Expérimentale, 54(1), 42.

Serre, T., Oliva, A., & Poggio, T. (2007). A feedforward architecture accounts for rapid categorization. Proceedings of the National Academy of Sciences, 104(15), 6424–6429. https://doi.org/10.1073/pnas.0700622104

Sheslow, D., & Adams, W. (2003). Wide Range Assessment of Memory and Learning–Second Edition (WRAML2). Wide Range Inc., Delaware.

Silson, E. H., McKeefry, D. J., Rodgers, J., Gouws, A. D., Hymers, M., & Morland, A. B. (2013). Specialized and independent processing of orientation and shape in visual field maps LO1 and LO2. Nature Neuroscience, 16(3), 267–269. https://doi.org/10.1038/nn.3327

Sincich, L. C., & Horton, J. C. (2005). The circuitry of V1 and V2: integration of color, form, and motion. Annu. Rev. Neurosci., 28, 303–326.

Subbiah, I., & Caramazza, A. (2000). Stimulus-centered neglect in reading and object recognition. Neurocase, 6(1), 13–31. https://doi.org/10.1080/13554790008402754

Tanigawa, H., Lu, H. D., & Roe, A. W. (2010). Functional organization for color and orientation in macaque V4. Nature Neuroscience, 13(12), 1542–1548. https://doi.org/10.1038/nn.2676

Tipper, S. P., & Behrmann, M. (1996). Object-centered not scene-based visual neglect. Journal of Experimental Psychology: Human Perception and Performance, 22(5), 1261–1278. https://doi.org/10.1037/0096-1523.22.5.1261

Tootell, R. B. H., & Nasr, S. (2017). Columnar segregation of magnocellular and parvocellular streams in human extrastriate cortex. Journal of Neuroscience, 37(33), 8014–8032.

Torgesen, J. K., Wagner, R. K., & Rashotte, C. A. (2012). TOWRE-2 Examiner’s Manual. Austin, TX: Pro-Ed.

Townsend, J. T., & Ashby, F. G. (1983). Stochastic modeling of elementary psychological processes. CUP Archive.

Tse, P. U., & Palmer, S. E. (2012). Visual Object Processing. In Handbook of Psychology, Second Edition. https://doi.org/10.1002/9781118133880.hop204007

Turnbull, O. H., Beschin, N., & Della Sala, S. (1996). Agnosia for object orientation: Implications for theories of object recognition. Neuropsychologia, 35(2), 153–163. https://doi.org/10.1016/S0028-3932(96)00063-2

Turnbull, O. H., Laws, K. R., & McCarthy, R. A. (1995). Object Recognition without Knowledge of Object Orientation. Cortex, 31(2), 387–395. https://doi.org/10.1016/S0010-9452(13)80371-1

Turnbull, O. H., & McCarthy, R. A. (1996). Failure to Discriminate between Mirror-image Objects: A Case of Viewpoint-independent Object Recognition? Neurocase, 2(1), 63–72. https://doi.org/10.1080/13554799608402390

Valtonen, J., Dilks, D. D., & McCloskey, M. (2008). Cognitive representation of orientation: A case study. Cortex, 44(9), 1171–1187. https://doi.org/10.1016/j.cortex.2007.06.005

Vaughn-Blount, K., Watson, S. T., Kokol, A. L., Grizzle, R., Carney, R. N., Rich, S. S., … Maricle, D. E. (2011). Wechsler Intelligence Scale for Children, Fourth Edition. In Encyclopedia of Child Behavior and Development (pp. 1553–1555). https://doi.org/10.1007/978-0-387-79061-9_3066

Vernon, R. J. W., Gouws, A. D., Lawrence, S. J. D., Wade, A. R., & Morland, A. B. (2016). Multivariate Patterns in the Human Object-Processing Pathway Reveal a Shift from Retinotopic to Shape Curvature Representations in Lateral Occipital Areas, LO-1 and LO-2. Journal of Neuroscience, 36(21), 5763–5774. https://doi.org/10.1523/JNEUROSCI.3603-15.2016

Wagemans, J., Elder, J. H., Kubovy, M., Palmer, S. E., Peterson, M. A., Singh, M., & von der Heydt, R. (2012). A century of Gestalt psychology in visual perception: I. Perceptual grouping and figure–ground organization. Psychological Bulletin, 138(6), 1172–1217. https://doi.org/10.1037/a0029333

Wagner, R., Torgesen, J., Rashotte, C., & Pearson, N. A. (1999). CTOPP-2: Comprehensive Test of Phonological Processing–Second Edition. Pro-ed Austin, TX.

Wandell, B. A., Dumoulin, S. O., & Brewer, A. A. (2007). Visual Field Maps in Human Cortex. Neuron, 56(2), 366–383. https://doi.org/10.1016/j.neuron.2007.10.012

Wiig, E. H., Semel, E. M., & Secord, W. (2003). CELF 5: Clinical evaluation of language fundamentals. Pearson/PsychCorp.

Yabuta, N. H., Sawatari, A., & Callaway, E. M. (2001). Two functional channels from primary visual cortex to dorsal visual cortical areas. Science. https://doi.org/10.1126/science.1057916

Yamins, D. L. K., Hong, H., Cadieu, C. F., Solomon, E. A., Seibert, D., & DiCarlo, J. J. (2014). Performance-optimized hierarchical models predict neural responses in higher visual cortex. Proceedings of the National Academy of Sciences, 111(23), 8619–8624. https://doi.org/10.1073/pnas.1403112111

